# Mathematical genomics of hidden signatures in NCF1: implications for chronic granulomatous disease

**DOI:** 10.64898/2026.11.12.766202

**Authors:** Sk. Sarif Hassan, Kharerin Hungyo, Vladimir N. Uversky

## Abstract

Inborn errors of immunity (IEIs), such as Chronic Granulomatous Disease (CGD), impose a disproportionately severe clinical burden in populations with high rates of consanguinity. CGD is caused by defects in the phagocyte NADPH oxidase complex, frequently due to mutations in the *NCF1* gene encoding the p47phox protein. While the functional domains of NCF1 are well characterized, the quantitative architectural constraints governing its structural stability remain elusive. This study aimed to decipher the multidimensional quantitative signatures of NCF1, across diverse organisms in order to understand its conserved spatial sequence architecture and the structural basis of variant severity. We analyzed 41 NCF1 sequences using ten distinct quantitative features, integrating primary sequence metrics, biophysical chemistry, and abstract topological geometry. The synthesis of these features revealed a lack of a unified proximity matrix, demonstrating that NCF1 is defined by feature-dependent, multidimensional individual signatures. Primary sequence metrics exhibited rigid phylogenetic conservation, revealing invariant zero-entropy blocks that correlated strictly with severe CGD-causing mutations, while topological metrics exposed a deep spatial architectural convergence. These multidimensional baselines confirmed that the protein requires a highly specific spatial packing to assemble the NADPH oxidase complex; pathogenic mutations therefore disrupt precise structural signatures rather than globally destroying the protein. Consistent with this, the AlphaFold model places the folded PX and tandem SH3 domains in a high-confidence core and the regulatory C-terminal tail in an exposed, low-confidence region, supporting a tail-mediated autoinhibitory mechanism. Ultimately, this quantitative framework offers a useful perspective on NCF1 architecture and may serve as a supportive tool for evaluating the severity of clinical variants, contributing to precision medicine efforts for NCF1-associated diseases.

## 1. Introduction

Inborn errors of immunity (IEIs) form a heterogeneous group of genetic disorders that disrupt the development and function of the immune system, leaving patients highly susceptible to infections, autoimmunity, and autoinflammation [1, 2, 3, 4, 5]. In populations with high rates of consanguinity, the clinical burden of IEIs is disproportionately high, since consanguineous unions strongly favor the manifestation of autosomal recessive forms of these disorders [6, 7, 8]. Among the most severe IEIs is Chronic Granulomatous Disease (CGD), a phagocytic immunodeficiency caused by defects in the NADPH oxidase complex [9, 10, 11]. Epidemiological studies in populations with elevated consanguinity have reported incidence rates as high as 5.2 cases per 100,000 person-years, with the autosomal recessive form being exclusively observed [12, 13]. Genotyping of affected cohorts has specifically identified pathogenic mutations in the Neutrophil cytosol factor 1 (*NCF1*) gene, including the c.75_76delGT variant, underscoring its critical clinical relevance [14, 15].

The *NCF1* gene encodes the p47phox protein, a 390-amino-acid cytosolic subunit essential for the assembly and activation of the phagocyte NADPH oxidase (NOX2) complex [16, 17, 18]. Structurally, p47phox features an N-terminal Phox homology (PX) domain, tandem Src homology 3 (SH3) domains, an autoinhibitory region (AIR), and a C-terminal proline-rich region (PRR) [19, 16, 20]. In resting cells, intramolecular interactions between the SH3 domains and the AIR keep the protein in an auto-inhibited state [19, 20]. Upon cellular activation, phosphorylation relieves this autoinhibition, allowing p47phox to translocate to the phagosome membrane and assemble the active oxidase complex that produces reactive oxygen species (ROS) [19, 21]. Mutations that disrupt this delicate spatial and structural balance abolish ROS production, leading to the fatal immunodeficiency characteristic of CGD or, in the case of hypomorphic variants, contributing to autoimmune dysregulation such as Systemic Lupus Erythematosus (SLE) [22].

While the functional domains of NCF1 are well characterized, a deeper quantitative understanding of its sequence architecture across diverse organisms is needed to identify the fundamental biophysical constraints that govern its structural stability [23]. To address this, we retrieved the protein sequence for NCF1 (p47phox) from the [UniProtKB](https://www.uniprot.org/uniprotkb?query=%28protein_name%3A%22Neutrophil+cytosol+factor+1%22%29+AND+%28gene%3ANCF1%29+AND+%28fragment%3Afalse%29+AND+%28length%3A%5B380+TO+400%5D%29). This yielded a robust dataset of 41 unique protein entries across various vertebrate species, encompassing mammals, birds, reptiles, amphibians, and fish.

Deciphering the hidden structural constraints within these sequences requires moving beyond simple linear comparisons and employing mathematical genomics. A couple of tools such as Chaos Game Representation (CGR), fractal dimension analysis, and hydropathy-based n-gram profiling translate symbolic sequences into geometric representations, capturing long-range correlations and organizational complexity that standard homology searches miss. By integrating ten distinct quantitative features—ranging from primary sequence metrics and biophysical chemistry to abstract topological geometry—this study aimed to construct a multidimensional proximity landscape of NCF1. We hypothesized that a single, unified proximity matrix would not exist; instead, NCF1 would be defined by feature-dependent individual signatures, wherein phylogenetic history, local biophysical chemistry, and topological geometry operate independently. The outcomes of this study provide a comprehensive quantitative framework for evaluating the structural severity of clinical NCF1 variants. Conclusively, this multidimensional profiling offers a useful perspective on how genetic variations disrupt protein function and may provide a helpful foundation for assessing disease severity and informing future diagnostic and therapeutic directions for NCF1-associated diseases.

## 2. Data acquisition

The protein sequence dataset for Neutrophil cytosolic factor 1 (NCF1/p47-phox) was retrieved from the [UniProt Knowledgebase (UniProtKB)](https://www.uniprot.org/uniprotkb?query=%28protein_name%3A%22Neutrophil+cytosol+factor+1%22%29+AND+%28gene%3ANCF1%29+AND+%28fragment%3Afalse%29+AND+%28length%3A%5B380+TO+400%5D%29) using the Advanced Search function. The search was performed on 2nd November, 2026 using the following specific criteria:

~~~
Protein Name [DE]: “Neutrophil cytosolic factor 1, p47-phox”
AND Gene Name [GN]: NCF1
AND Sequence length: 380 to 400
AND Fragment: No
~~~

This stringent filtering ensured the retrieval of full-length (non-fragment) NCF1 orthologs within the 380–400 amino acid range. The search yielded a total of 41 unique protein entries across various vertebrate species. For each entry, the UniProt entry name, source organism, and sequence length were manually extracted and compiled (Table 1). Because multiple entries were identified for certain species, the fundamental unit of analysis for this study is the individual protein entry rather than the species.

**Table 1:**
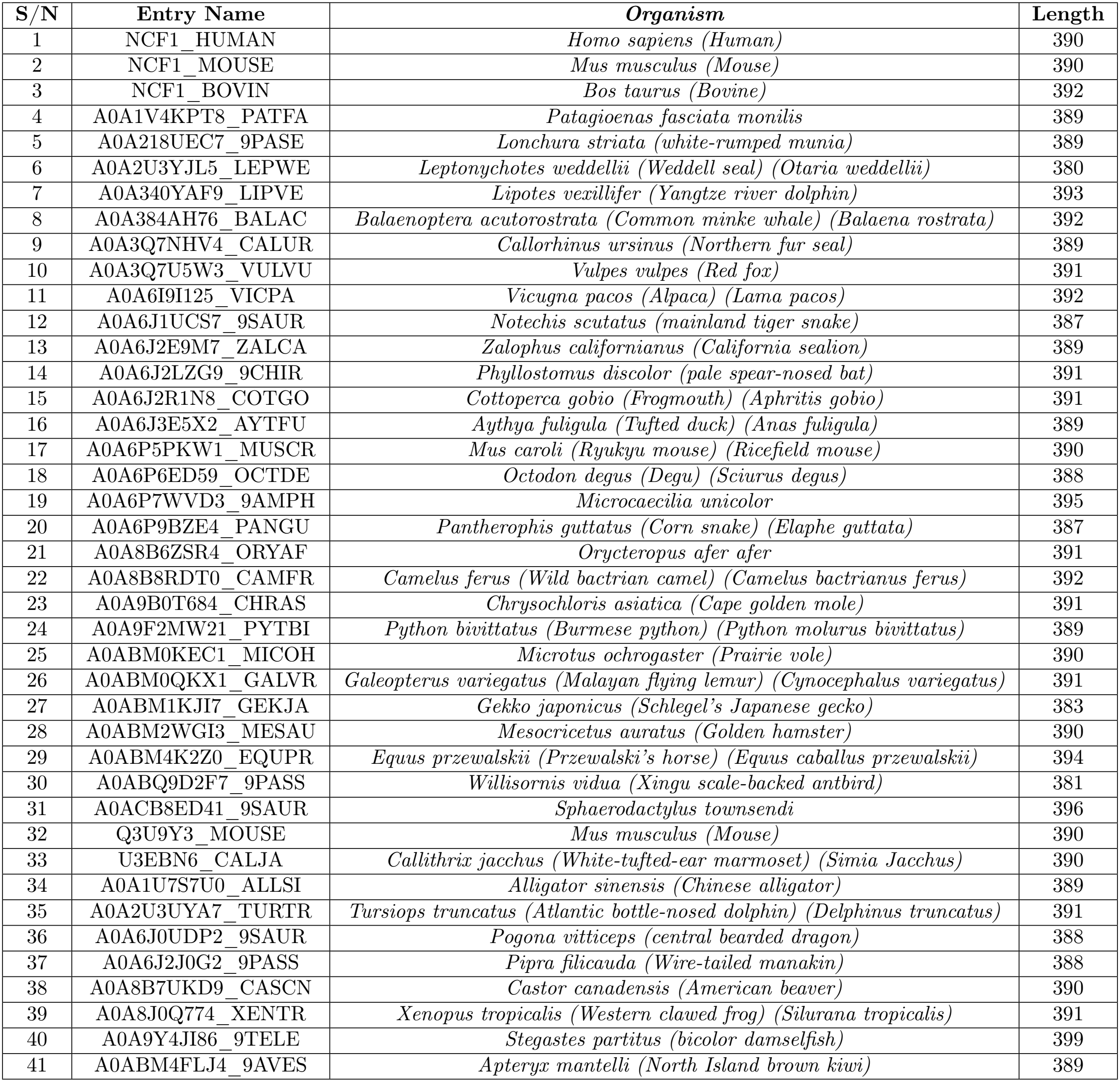
List of NCF1 (p47-phox) protein entries analyzed in the present study. Columns represent the serial number (S/N), UniProt entry name, source organism, and protein sequence length (in amino acids).

## 3. Methods

### 3.1. Amino acid frequency distribution of NCF1 proteins across various organisms

#### 3.1.1. Relative amino acid frequency of NCF1 Proteins across organisms

For each NCF1 protein sequence, we quantified amino acid composition by counting how often each of the 20 standard amino acids occurred. To account for differences in sequence length, we converted these counts into percentages of the total sequence length, which gave the relative abundance of each amino acid within the corresponding NCF1 protein sequence. Each NCF1 protein sequence was therefore represented as a 20-dimensional feature vector containing the relative frequencies of the standard amino acids.

#### 3.1.2. Shannon entropy-based analysis of compositional diversity in NCF1 protein sequences

To quantify compositional diversity in the NCF1 protein sequences, we used Shannon entropy, which measures the uncertainty in amino acid frequency distributions [24, 25, 26]. For each sequence, let *p*_*i*_ be the relative frequency of the *i*th amino acid (*i* = 1, …, 20). Shannon entropy (*H*) is calculated as

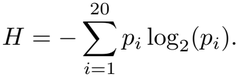

*H* gives a quantitative measure of amino acid diversity within a sequence. Lower values indicate reduced diversity and a predominance of certain amino acids, whereas higher values indicate a more even distribution of residues. For proteins made up of the 20 standard amino acids, the theoretical maximum entropy is 4.322, which occurs when all amino acids are equally represented [24, 26]. Shannon entropy therefore provides a useful way to compare compositional complexity and conservation among NCF1 protein sequences.

#### 3.1.3 Dendrogram relationships among NCF1 protein sequences based on amino acid composition

To explore compositional relationships among the NCF1 protein sequences, we represented each sequence as a 20-dimensional vector of relative amino acid frequencies [27, 28]. We then calculated pairwise Aitchison distances between all sequence vectors to quantify compositional dissimilarity. This distance measure is well suited to amino acid frequency data because it accounts for their compositional nature. The resulting distance matrices were symmetric and had dimensions of 41 *×*41 for the NCF1 protein sequences, thereby summarizing the pairwise compositional differences among them. Using these Aitchison distance matrices, we constructed dendrograms with the MATLAB phytree function to visualize compositional clustering and dendrogram relationships among the NCF1 proteins.

### 3.2. Frequency-dominant amino acid distribution across consecutive sequence windows in NCF1 protein sequences

Each NCF1 protein sequence was divided into consecutive, non-overlapping 20-residue windows to examine local patterns in amino acid composition [27]. When the terminal segment of an NCF1 protein sequence contained fewer than five residues, those remaining residues were merged with the preceding window to avoid bias from very short segments. Terminal segments containing more than five residues were treated as independent windows.

For each window, we calculated the frequencies of the 20 standard amino acids and identified the amino acid with the highest occurrence as the *dominant residue*. The dominance of a residue within a window was quantified as

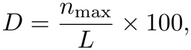

where *n*_max_ is the frequency of the most abundant amino acid in the window and *L* is the length of the corresponding window. Thus, *D* measures the percentage contribution of the dominant amino acid to the local composition of the NCF1 protein sequence.

For every window, we recorded the dominant amino acid, its percentage abundance, and its positions within the original sequence. The resulting dominance profiles were visualized as window-wise distributions, in which the percentage of the dominant amino acid was plotted across consecutive sequence windows. To allow direct comparison among different sequences, all plots used a standardized *y*-axis scale ranging from 0% to 100%. Background color coding was used to distinguish dominant amino acid types across windows, making it possible to visualize local compositional heterogeneity and residue dominance patterns within the NCF1 protein sequences.

### 3.3 Multiple sequence alignment-based phylogenetic analysis of NCF1 sequences

To investigate the evolutionary relationships among the NCF1 proteins, we performed a multiple sequence alignment using [Clustal Omega](https://www.ebi.ac.uk/jdispatcher/msa/clustalo) [29]. This alignment allowed us to map conserved residues, variable positions, and amino acid substitutions across the dataset, providing a solid foundation for assessing sequence homology and evolutionary divergence.

Alignments were run using the default Clustal Omega settings, employing the *Gonnet* substitution matrix with a gap opening penalty of 10 and a gap extension penalty of 0.20. End-gap penalties were disabled to match the standard alignment procedure implemented by the [Clustal Omega](https://www.ebi.ac.uk/jdispatcher/msa/clustalo) platform. These processed alignments were then used to reconstruct the phylogenetic relationships and to survey sequence conservation patterns across the NCF1 family.

### 3.4. Dendrogram relationships among NCF1 sequences based on dipeptide composition analysis

To capture the frequency of adjacent amino acid pairs, we calculated the dipeptide composition for each NCF1 protein sequence. For a sequence of length *L*, this approach yields a 400-dimensional vector representing all possible ordered dipeptides derived from the twenty standard amino acids [30, 31, 32]. By extracting overlapping dipeptides across each sequence, we computed the frequency of each dipeptide *ij* as:

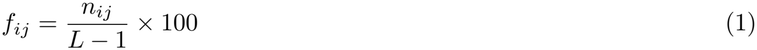

where *n*_*ij*_ represents the count of dipeptide *ij*, and *L* − 1 denotes the total number of dipeptide bonds.

To assess dipeptide prevalence across the NCF1 dataset, we defined presence as a non-zero frequency within a given sequence. For each dipeptide, we calculated several descriptive statistics: the number of sequences containing it, its prevalence (%), and its mean, median, standard deviation, minimum, and maximum frequencies (%). Prevalence was determined using:

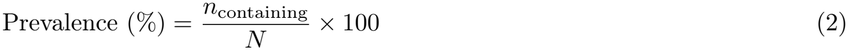

where *n*_containing_ is the number of sequences containing the dipeptide and *N* is the total number of NCF1 sequences. Based on these values, we classified the dipeptides into five prevalence categories: absent (0%), rare (*≤* 25%), moderately prevalent (26–50%), common (51–75%), and highly prevalent (>75%), while treating those absent from all sequences separately. Finally, to explore the compositional relationships among the proteins, we computed pairwise Aitchison distances (Euclidean distances between the dipeptide frequency vectors) for all NCF1 sequence pairs. We then applied hierarchical clustering using average linkage to generate a dendrogram mapping the structural relationships among the sequences.

### 3.5. Longest polystring analysis and dendrogram relationships across NCF1 sequences

To characterize homopolymer runs, we determined the longest polystring for each of the 20 standard amino acids across all NCF1 sequences [30]. The longest polystring was defined as the longest consecutive stretch of a single residue; where multiple runs shared the same maximum length, the first occurrence was retained. For each sequence and amino acid, we recorded the longest run length and its corresponding polystring, alongside a normalized longest-run percentage. This percentage was calculated as *L*_run_*/L×* 100, where *L*_run_ is the longest run length and *L* is the total sequence length.

Using these normalized longest-run percentages, we constructed a dendrogram to explore the relationships among the NCF1 sequences. We assembled the percentages for the 20 amino acids into a 20-dimensional feature vector for each sequence and calculated pairwise Euclidean distances to generate a comprehensive distance matrix. Hierarchical clustering was then performed using average linkage. Clusters were defined by cutting the tree at a distance threshold of 0.5, with cluster membership recorded for each sequence. This same threshold was applied for branch colouring to visually group the sequences, though the threshold line itself was omitted, as the cutoff served purely for visual grouping and cluster assignment.

### 3.6. Hydropathy-based n-gram frequency analysis of NCF1 sequences

To capture local hydropathy patterns, we divided the 20 standard amino acids (*A*) into two distinct groups: polar (*P*) and nonpolar/hydrophobic (*ℋ*):

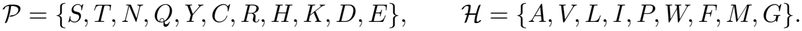

Each amino acid sequence *S* = (*s*_1_, *s*_2_, …, *s*_*L*_) of length *L* was then translated into a binary string *E* = (*e*_1_, *e*_2_, …, *e*_*L*_), where each residue was encoded as:

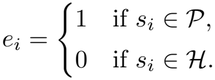

An *n-gram* in this context is any contiguous pattern of length *n* within the binary string, representing a specific local hydropathy sequence (i.e., a word in *{*0, 1*}*^*n*^)) [27]. To analyze these patterns, we slid a window of length *n* across each encoded sequence, enumerating all overlapping *n*-grams. This generates a set of possible patterns *K*_*n*_ of size 2^*n*^. For each pattern *w ∈ K*_*n*_, we defined its raw count *N* (*w*) and relative frequency *f* (*w*) as:

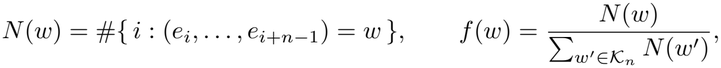

meaning *f* (*w*) represents the proportion of pattern *w* among all observed overlapping *n*-grams in the sequence. To illustrate this process, consider the short binary sequence *E* = (1, 0, 1, 1, 0, 0, 1, 0, 1, 0). Using a window of length 2,we extract the overlapping 2-grams: 10, 01, 11, 10, 00, 01, 10, 01, 10. Tallying these yields counts of *N* (10) = 4, *N* (01) = 3, *N* (11) = 1, and *N* (00) = 1, corresponding to relative frequencies of 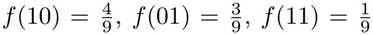, and 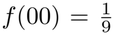 Similarly, a window of length 3 yields the overlapping 3-grams: 101, 011, 110, 100, 001, 010, 101, 010. This results in counts of *N* (101) = 2, *N* (010) = 2, *N* (011) = 1, *N* (110) = 1, *N* (100) = 1, and *N* (001) = 1, with the remaining 3-grams having a count of zero. The same sliding-window logic applies directly to higher values of *n*.

Finally, we used these relative frequencies to compare the sequences. Specifically, for *n* = 2, 3, and 4, we computed pairwise Euclidean distances between all 41 NCF1 sequences based on their independent *n*-gram frequency profiles. This produced symmetric distance matrices of dimension 41*×* 41 for each *n*. We then applied hierarchical clustering to these distance matrices to construct dendrograms, visually mapping the sequence proximities. By applying a distance threshold to the resulting trees, we delineated clusters of NCF1 sequences that share similar hydropathy-based *n*-gram profiles.

### 3.7. Three-dimensional Chaos Game Representation and fractal dimension analysis

To capture the spatial heterogeneity and long-range correlations of the NCF1 protein sequences, we encoded each sequence into a three-dimensional geometric space using a Chaos Game Representation (CGR). Because proteins comprise twenty standard amino acids, we mapped each residue to the 20 unique vertices of a regular dodecahedron, a Platonic solid naturally suited for this dimensionality. This geometric arrangement ensures maximal spatial separation between chemically distinct residues, distributing the 20 amino acids across the vertices in the 3D Cartesian space as previously established [33, 34].

For a protein sequence *S* = (*s*_1_, *s*_2_, …, *s*_*L*_) of length *L*, the CGR trajectory was generated iteratively. The spatial position **r**_*n*_ *∈* ℝ^3^ of the *n*-th step was defined by the contraction mapping:

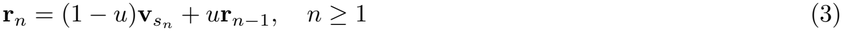

where 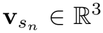 is the dodecahedron coordinate of the amino acid *s*_*n*_, *u* is the contraction factor, and **r**_0_ was initialized at the geometric center of the coordinate space inscribed within a unit ball, **r**_0_ = (1, 1, 1). The parameter *u* dictates the degree to which the trajectory is pulled toward the new vertex versus how much it retains its previous position (i.e., its “memory”).

To comprehensively explore how different degrees of spatial contraction influence the resulting geometric complexity, we evaluated four specific contraction factors: *u ∈ {*0.15, 0.25, 0.35, 0.50}. For each value of *u*, the amino acid sequence was iterated cyclically to generate a dense, invariant point cloud of *N* = 50, 000 points per protein. This cyclic iteration ensures that the compositional biases of the sequence fully manifest as a geometric attractor rather than a sparse, transient trajectory.

The spatial complexity of the resulting three-dimensional point clouds was quantified using the box-counting fractal dimension (*D*_B_). The coordinates of each attractor were first normalized to the unit cube [0, 1]^3^. The cube was then partitioned into cubic grids of varying side length *ε*, and the number of occupied boxes *N* (*ε*) was counted across a scaling range of *ε∈* [10^*−*1.5^, 10^*−*0.3^]. The fractal dimension was estimated as the absolute slope of the linear regression of log *N* (*ε*) against log(1*/ε*):

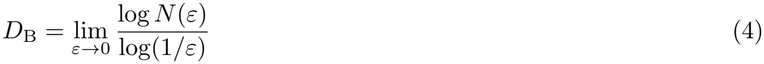

This *D*_B_ provided a quantitative measure of the structural heterogeneity encoded within each NCF1 sequence, revealing how the spatial distribution of amino acids scales with the resolution of observation.

### 3.8. Hydropathy dance of NCF1 sequences

An amino-acid sequence of NCF1 protein *s* = *s*_1_*s*_2_ *· · · s*_*N*_ is reduced to a binary sequences

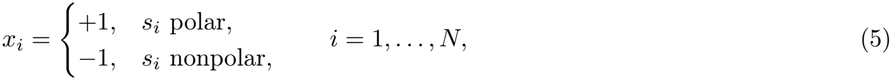

using a fixed hydrophobicity partition applied identically to all sequences. A fixed-direction walk, in which *x*_*i*_ = +1 and *x*_*i*_ = *−*1 are assigned two constant step directions, advances one coordinate per residue and therefore records global composition only [35]. The dance replaces the constant direction set by a position-indexed heading field, so that the local composition and the local arrangement of residues around position *i* determine the step direction at *i*.

#### Local composition and texture fields

For an window of width *w ∈* ℤ_*≥*1_ the local polarity is

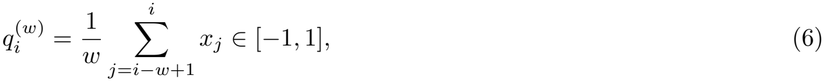

and, for *w ≥* 2, the local alternation (texture) index is

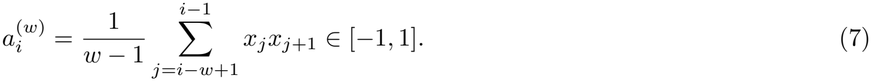

The field *q*^(*w*)^ measures net local composition; *a*^(*w*)^ measures local order, taking the value +1 for contiguous runs, *−*1 for strict alternation, and 0 for uncorrelated arrangement. The pair 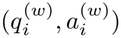 therefore separates sequences that share a composition: PPPPNNNN, PNPNPNPN and an equal-composition random arrangement all have 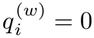, and differ in 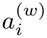 by sign and magnitude.

#### Heading field and trace

Fix a number of scales *K ∈* ℤ_*≥*1_, a base window *w*_0_ *∈* ℤ_*≥*2_, an exponent *H*_*s*_ *∈* (0, 1], couplings *κ*_0_ *>* 0, *λ ≥* 0, *µ ≥* 0, and irrational frequencies *γ, γ*^*′*^ *∈* (0, 1). Set

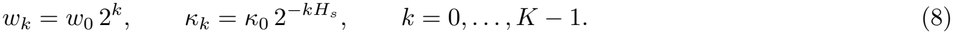

The heading at position *i* is

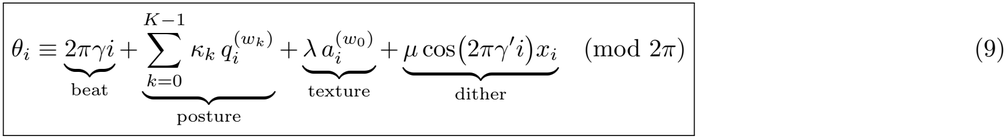

and the trace is the cumulative unit-step path

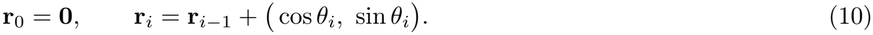

The four terms of Eq. (9) have distinct functions. The *beat* 2*πγi*, with *γ* algebraic irrational, provides an aperiodic carrier that prevents lattice locking and resonance with the dyadic windows. The *posture* sum steers the heading by local composition measured over *K* octaves with self-similar amplitudes *κ*_*k*_, so that a domain of length *l* perturbs the geometry at every scale *w*_*k*_ ≳ *l*; spatial composition, rather than residue identity alone, thereby determines the trace. The *texture* term makes the map sensitive to arrangement at fixed composition. The *dither* term couples residue identity to a second incommensurate rotation, so that two sequences differing at a single position cannot produce identical headings. The exponent *H*_*s*_ controls the scale similarity of the steering: at 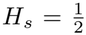 the summed perturbation from a domain of size *l* is independent of *l* across scales *w*_*k*_*≥l*, which is the condition under which the trace admits a nontrivial scaling range rather than a single characteristic length.

#### Curvature and the boundary variant

The primitive quantity of the encoding is the curvature

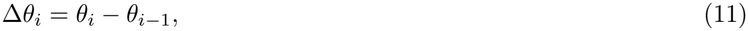

from which the trace (10) is recovered by integration; Eq. (9) defines the *steering dance*. A variant, the *boundary dance*, prescribes the curvature directly,

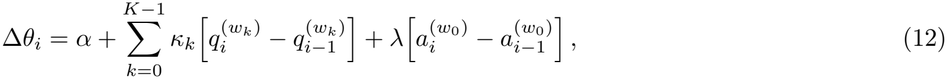

so that the heading responds to changes in local composition and the geometry marks domain boundaries rather than domain interiors; periodic amphipathic structure then appears as a characteristic loop in the trace.

#### Fractal and multifractal exponents

Let *N* (*ε*) be the number of boxes of side *ε* required to cover 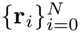. The box-counting dimension is

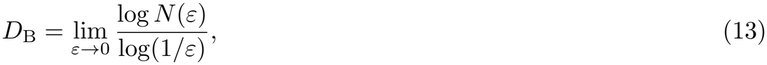

estimated by least-squares regression of log *N* (*ε*) on log(1*/ε*) over *ε* [1, *ρ*], where *ρ* is the trace diameter. For a self-affine trajectory with Hurst exponent *H*,

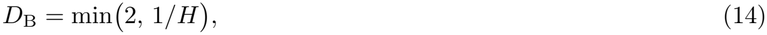

so that *D*_B_ is a geometric readout of the persistence of the local composition field: smooth drift gives *H →* 1 and *D*_B_ *→* 1;uncorrelated arrangement gives 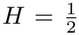 and *D*_B_ *→* 2; anti-persistent alternation saturates *D*_B_ = 2. As an independent dynamical estimate we regress the mean-square displacement

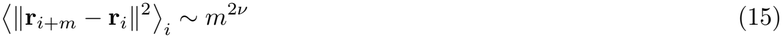

over *m* ∈ [1, *N*/4].

To resolve hierarchical clustering that a single exponent averages out, assign each box *B* the visit measure *µ*(*B*) =*N*_*B*_*/N*, where *N*_*B*_ is the number of trace points in *B*, and form the partition function

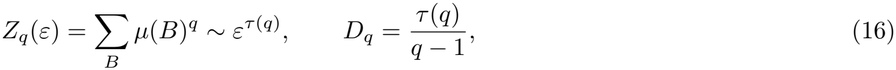

with *q∈* [*−*5, 5] and the *q →*1 limit obtained by interpolation of *τ* (*q*). The generalised dimensions *D*_0_ (support), *D*_1_(information) and *D*_2_ (correlation) constitute a multifractal fingerprint of the spatial composition of the sequence.

*Algorithm*

##### Algorithm 1

Dance encoding of a binary sequence

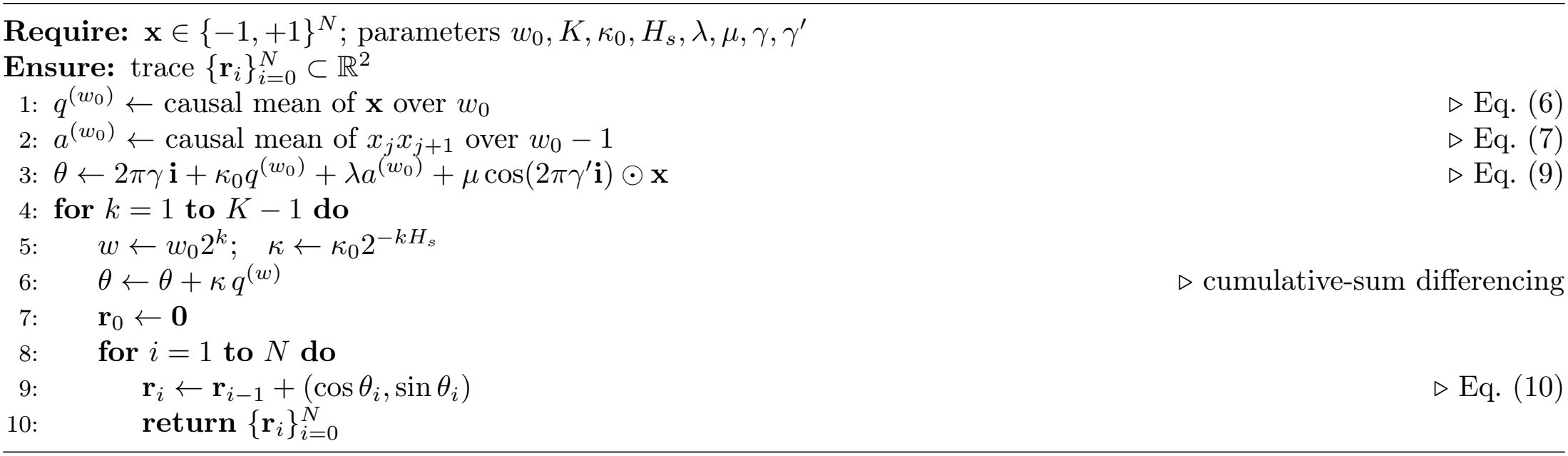

All sliding sums in Eqs. (6)–(7) are evaluated by cumulative-sum differencing, so the encoding costs *O*(*KN*) time and *O*(*N*) memory. Box counting over *B* scales costs *O*(*BN*), and the multifractal partition function *O*(*BN*) for all *q*.

Furthermore, it was noted that the parameters were fixed a priori and applied unchanged to every NCF1 sequences. Defaults are listed in Table 2.

**Table 2:**
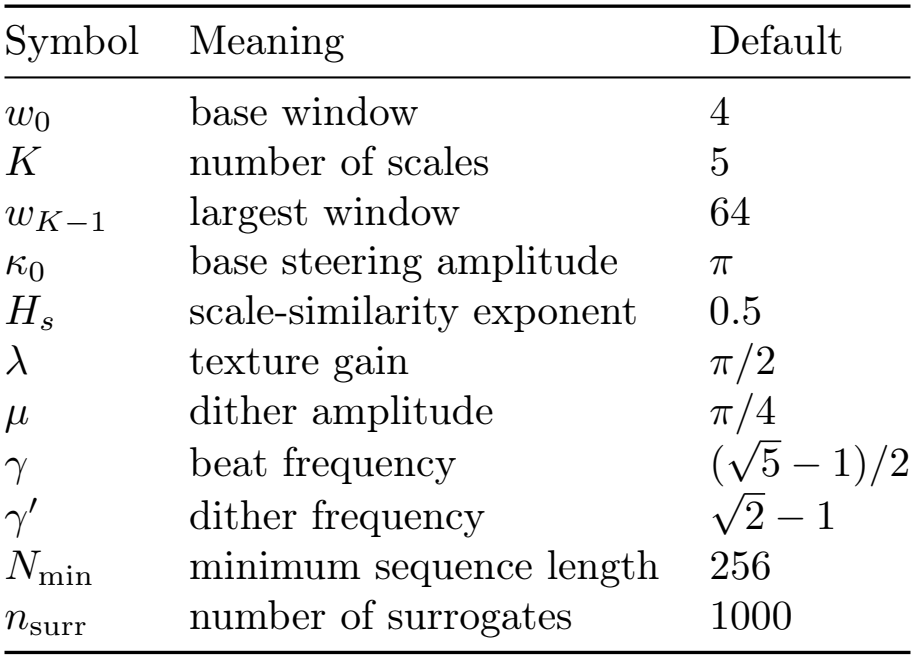
Default parameter values for the dance encoding considered in this study.

| Symbol | Meaning | Default |
|---|---|---|
| *w*<sub>0</sub> | base window | 4 |
| *K* | number of scales | 5 |
| *w*<sub>*K*−1</sub> | largest window | 64 |
| κ<sub>0</sub> | base steering amplitude | π |
| *H*<sub>*s*</sub> | scale-similarity exponent | 0.5 |
| λ | texture gain | π/2 |
| μ | dither amplitude | π/4 |
| γ | beat frequency | (√5 − 1)/2 |
| γ′ | dither frequency | √2 − 1 |
| *N*<sub>min</sub> | minimum sequence length | 256 |
| *n*<sub>surr</sub> | number of surrogates | 1000 |

With *κ*_0_ = *π* a fully polar window (*q* = 1) rotates the heading by *−π* and a fully nonpolar window by *π*, so that composition is mapped onto the full circle rather than onto a half-plane.

Therefore, each NCF1 sequence was summarised by the triple (*D*_B_, *{D*_*q*_*}, v)*, together with the scaling range used for each estimate. The dance converted the spatial composition field of the polar/nonpolar encoding into a geometric exponent: long polar or nonpolar domains yielded smooth arcs with *D*_B_ *→*1; blocky, but finite domains yielded 1 *< D*_B_ *<* 2; shuffled and strongly alternating sequences both saturated *D*_B_ = 2, but were separated by their multifractal spectra. Sequences of identical composition and different arrangement were thereby distinguished by their (*D*_B_, *D*_*q*_) fingerprint.

Finally, to establish a hierarchical relationship among the 41 NCF1 sequences based on their dance metrics, the computed exponents were used to define a feature space. For each sequence, a feature vector was constructed from the fractal and dynamical exponents, ***ϕ*** = (*D*_B_, *v, D*_0_, *D*_1_, *D*_2_). Pairwise Euclidean distances were then calculated between the standardised feature vectors of all 41 sequences. Using this distance matrix, agglomerative hierarchical clustering was performed, and the resulting hierarchical relationships were visualised as a dendrogram. This clustering grouped NCF1 sequences with similar spatial composition profiles, thereby revealing distinct structural relationships within the set of all NCF1 protein sequences across organisms.

#### 3.8.1. Illustration of the dance encoding on a toy example

To demonstrate that the dance encodes spatial arrangement rather than merely global composition, we apply the method to four synthetic 128-residue sequences that all contain exactly 64 polar (P) and 64 nonpolar (N) residues but differ in their sequence order:

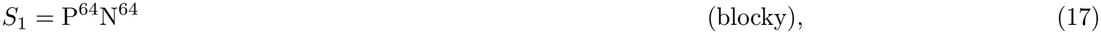

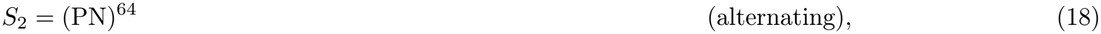

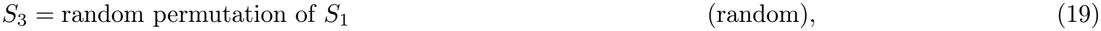

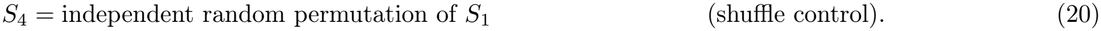

The sequences were processed using the default parameters of Table 2 (with beat frequency 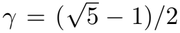 and the resulting traces are shown in Figure 1.

**Figure 1.**
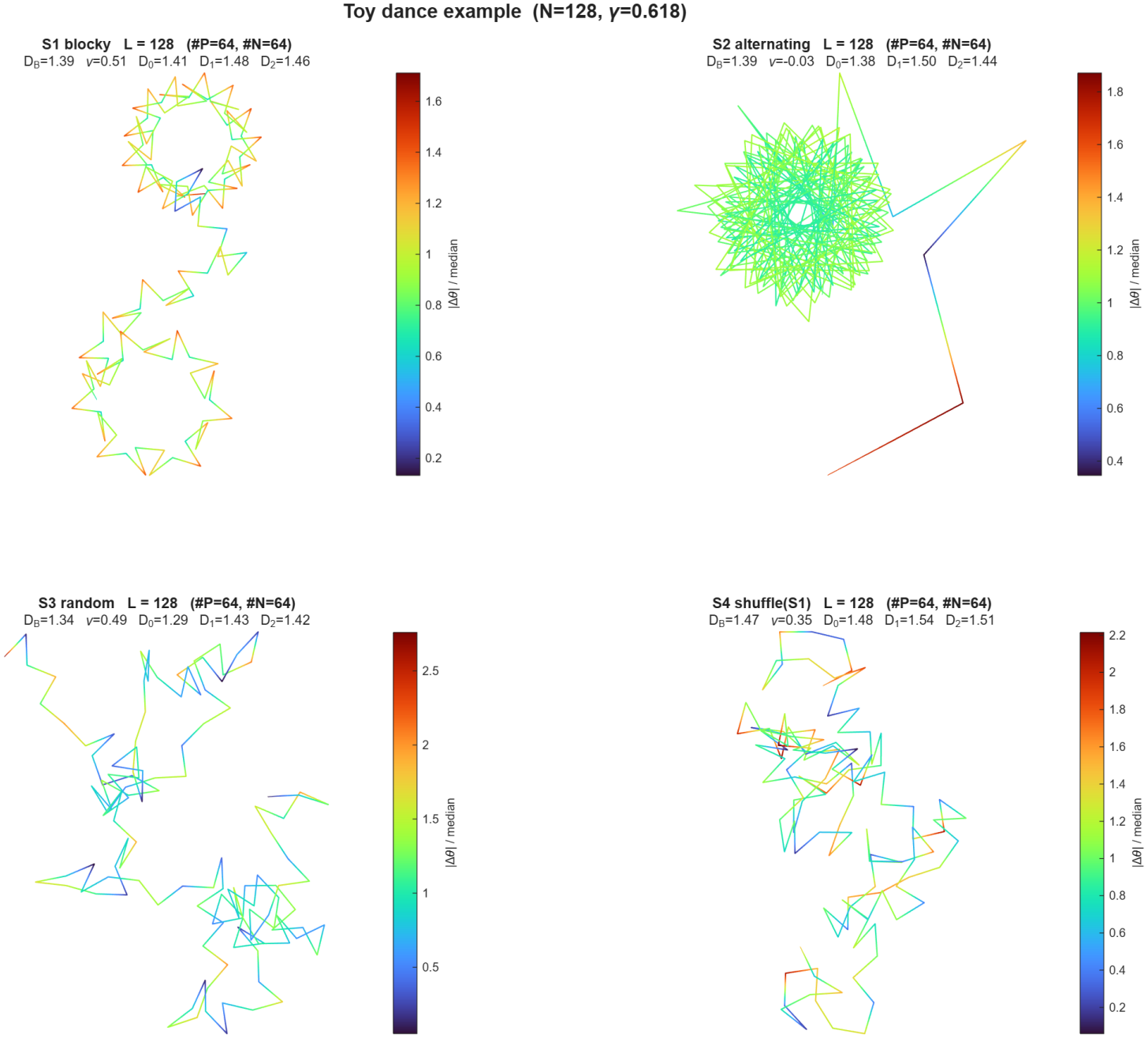
Toy dance example. Traces for the four sequences of Eqs. (17)–(20), colored by the normalized curvature |Δ*θ*_*i*_|/median. All sequences have *L* = 128 and identical composition. (a) Blocky *S*_1_: long coherent loops. (b) Alternating *S*_2_: a tightly bounded, high-curvature rosette.(c) Random *S*_3_ and (d) shuffle *S*_4_: diffusive, area-filling paths. The metrics are listed in Table 3.

The four sequences yield visually distinct geometric trajectories. Because *S*_1_ consists of long contiguous blocks, the heading field changes slowly within the homopolymer stretches, producing two large, coherent, smooth loops joined by a sharp transition at the P/N boundary. In contrast, the rapidly alternating sequence *S*_2_ causes the posture term to cancel, leaving the heading dominated by the beat and dither. This results in a tightly bounded, high-frequency rosette (star-like) trajectory. The random sequences *S*_3_ and *S*_4_ produce diffusive, area-filling paths that lack the coherent large-scale structure of *S*_1_.

**Table 3:** Metrics for the toy sequences of Eqs. (17)–(20). All sequences have identical composition (*N* = 128, #*P* = 64, #*N* = 64).

| Sequence | *D*<sub>B</sub> | *ν* | *D*<sub>0</sub> | *D*<sub>1</sub> | *D*<sub>2</sub> |
|---|---|---|---|---|---|
| *S*<sub>1</sub> blocky | 1.39 | 0.51 | 1.41 | 1.48 | 1.46 |
| *S*<sub>2</sub> alternating | 1.39 | −0.03 | 1.38 | 1.50 | 1.44 |
| *S*<sub>3</sub> random | 1.34 | 0.49 | 1.29 | 1.43 | 1.42 |
| *S*<sub>4</sub> shuffle(*S*<sub>1</sub>) | 1.47 | 0.35 | 1.48 | 1.54 | 1.51 |

Table 3 reports the fractal and dynamical exponents obtained from these traces. The metric most sensitive to the difference between the blocky and alternating regimes is the displacement exponent *v*, derived from the mean-square displacement. The blocky sequence *S*_1_ exhibits *v* = 0.51, characteristic of a persistent, directed trajectory over the length of the blocks, whereas the alternating sequence *S*_2_ yields *v* = *−*0.03, reflecting a bounded oscillator whose mean-square displacement saturates (and thus lacks a positive scaling slope). The random sequences *S*_3_ and *S*_4_ yield intermediate positive values (*v* = 0.49 and *v* = 0.35, respectively).

The box-counting dimensions *D*_B_ for these sequences range between 1.34 and 1.47, with the two independent shuffles of the same composition (*S*_3_ and *S*_4_) differing by Δ*D*_B_ *≈* 0.13. This spread reflects the finite-size variance inherent to the box-counting estimator on random, area-filling walks and highlights the necessity of using shuffled surrogates as baselines when comparing real protein sequences. The generalized dimensions (*D*_0_, *D*_1_, *D*_2_) provide an additional fingerprint; for example, the alternating sequence *S*_2_ has a higher information dimension (*D*_1_ = 1.50) than the blocky sequence *S*_1_ (*D*_1_ = 1.48), reflecting the dense, uniform filling of its bounded rosette, even though their *D*_B_ values are nearly identical.

### 3.9. Functional intrinsic disorder analysis of NCF1 sequences

The intrinsic disorder propensity of human NCF1 (UniProt ID: P14598; 390 amino acids) was analyzed using the [Rapid Intrinsic Disorder Analysis Online (RIDAO)](https://ridao.app/) platform, which assembles the outputs of six commonly used predictors: PONDR^®^ VLS2, PONDR^®^ VL3, PONDR^®^ VLXT, PONDR^®^ FIT, IUPred-Long, and IUPred-Short [36, 37]. The mean disorder profile (MDP) was calculated by averaging the per-residue scores of the individual predictors, and the standard deviation across predictors was plotted to represent profile variability. Residues with disorder scores *≥* 0.5 were classified as intrinsically disordered, whereas residues with scores between 0.15 and 0.5 were considered flexible. The overall disorder content of NCF1 was quantified using the PONDR^®^ VLS2-predicted disorder content (PDC, expressed as a percentage of residues classified as disordered) and the mean disorder score (MDS, reported as mean *±* standard deviation). Proteins were classified according to the accepted field criteria: highly ordered (PDC *<* 10%), moderately disordered (10% *≤* PDC *<* 30%), and highly disordered (PDC*≥* 30%).

To identify putative disorder-based interaction sites, the ANCHOR2 profile was generated via the IUPred2A web server [38]. ANCHOR2 scores above 0.5 were taken to indicate a propensity for intrinsically disordered residues to gain stabilizing energy upon binding to a globular protein partner, thereby identifying putative molecular recognition features (MoRFs) [38].

In parallel, the functional disorder profile of NCF1 was obtained from the D^2^P^2^ platform, which integrates predictions from multiple disorder predictors and annotates disorder-based binding sites (MoRFs) and post-translational modification sites, including phosphorylation sites.

### 3.10. Determining proximal sets of NCF1 sequences

For each NCF1 sequence, six types of quantitative features were computed: (i) amino acid frequency distribution, (ii) amino acid sequence homology, (iii) dipeptide composition, (iv) longest polystring analysis, and (v) hydropathy-based *n*-gram frequency distribution (vi) four-dimensional fractal dimension (FD) profiles derived from the three dimensional chaos game representations, and (vii) hydropathy-based dance translated the spatial arrangement of polar and nonpolar residue [27, 30]. Each feature type yielded a feature vector for every NCF1 sequence, with the dimensionality depending on the feature (e.g., 20 for longest polystring of amino acids, and so on). To compare sequences, pairwise Euclidean distances were calculated between their feature vectors for each feature type separately. For a given feature, the distance between two sequences reflected their dissimilarity in that feature space. These distances were then used to identify NCF1 sequences that were proximal to one another. For each feature, a suitable distance threshold was applied to define the most proximal sets of NCF1 sequences. Sequences falling within the threshold were considered part of the same proximal set for that feature.

### 3.11. Cumulative feature-based clustering of NCF1 sequences

Robust proximal NCF1 sequences were identified by performing nine independent feature-based clustering analyses. The features used were amino acid frequency, sequence homology, dipeptide composition, longest polystring, 2-, 3-, and 4-gram hydropathy profiles, four-dimensional fractal dimension (FD) profiles, and hydropathy-based dance. For each feature, NCF1 sequences were clustered using hierarchical clustering with Euclidean distance and average linkage, and clusters were defined at a fixed threshold. The clusters obtained from all nine analyses were then intersected, and only those groups of sequences that co-clustered in all eight analyses were retained as cumulative proximal clusters [27, 30]. This intersection ensured that the final clusters were robust to the choice of feature and reflected consistent sequence similarity across multiple compositional and hydropathy descriptors. This integrative approach provides a multidimensional perspective on sequence similarity, capturing complementary signals from composition, hydropathy, and local order.

## 4. Results and analyses

### 4.1. Amino acid compositions and derived proximilty of NCF1 protein sequences

#### 4.1.1. Amino acid frequency distribution across NCF1 sequences

The analysis set out to characterize the amino acid frequency distribution across the 41 NCF1 sequences and to determine whether any sequence carried a compositional profile that stood apart from the rest. It was found that the highest-frequency positions were occupied mainly by arginine (R), glutamic acid (E), lysine (K), and proline (P), with serine (S) appearing as the maximum in a smaller number of NCF1 sequences. Cysteine (C), by contrast, was almost always the lowest-frequency residue (Figure 2).

**Figure 2.**
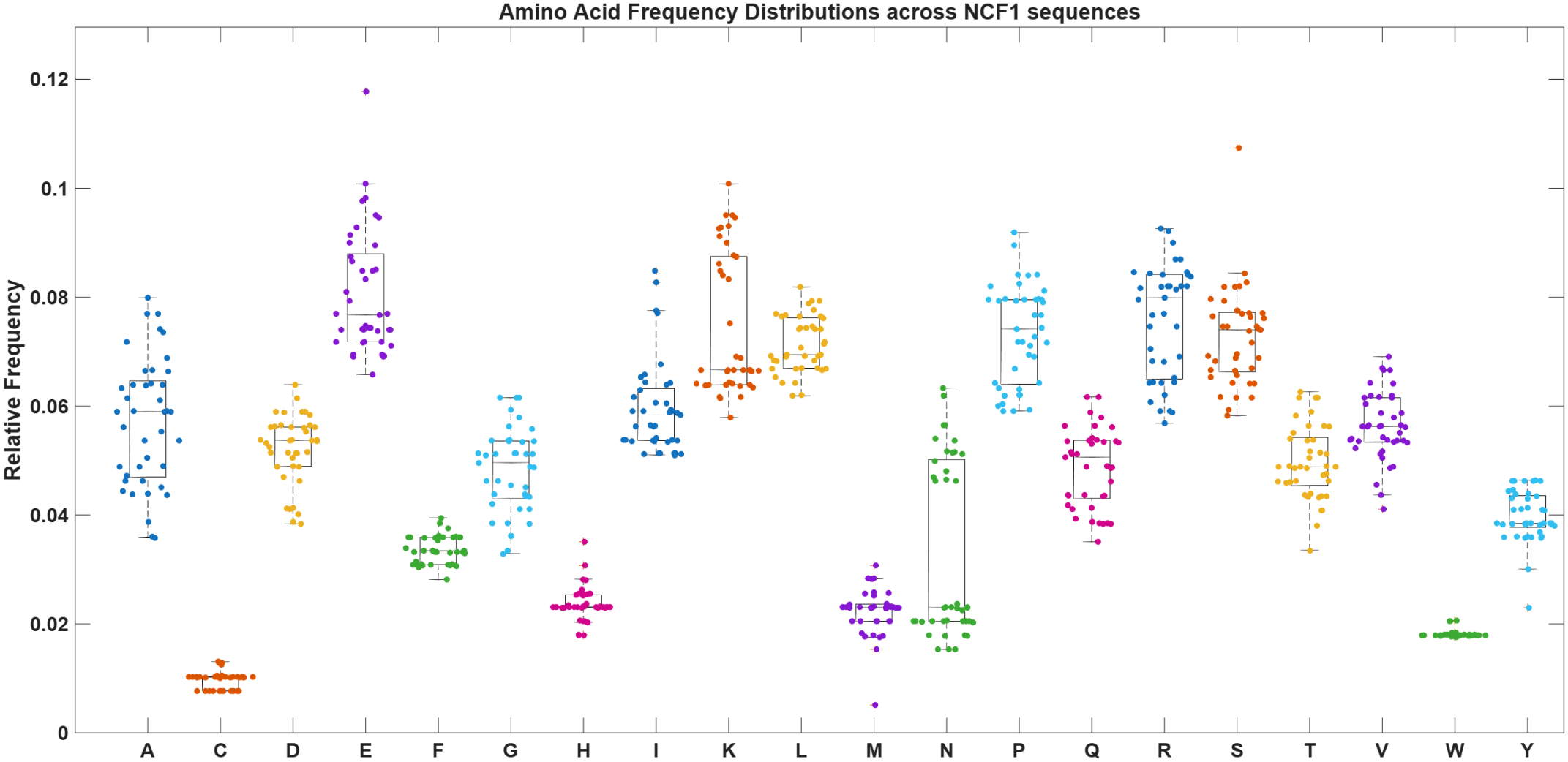
Amino acid frequency distribution across NCF1 sequences

The amino acid R was the most frequent maximum across NCF1 sequences. It accounted for 19 of the 51 top-frequency assignments (37.3%) and was the sole highest residue in 13 of the 41 sequences (31.7%). E followed, with 10 assignments (19.6%) and 7 unique maxima (17.1%). K contributed 9 assignments (17.6%) and 6 unique maxima (14.6%), P contributed 7 assignments (13.7%) and 4 unique maxima (9.8%), and S contributed 4 assignments (7.8%) and 2 unique maxima (4.9%). A and L appeared as highest residues only in tied positions, each with 1 assignment (2.0%). No NCF1 sequence had C, D, F, G, H, I, M, N, Q, T, V, W, or Y as its highest-frequency amino acid. The largest single percentage in the set of NCF1 sequences was E at 11.8% in A0A9Y4JI86_9TELE, followed by S at 10.7% in A0A6J2R1N8_COTGO and E/K at 10.1% in A0A6J1UCS7_9SAUR. Most other per-sequence maxima fell between about 7.7% and 9.3%. Across all sequences, the residues that frequently occupied the maximum position ranged as follows: R from 5.7% to 9.3%, E from 6.6% to 11.8%, K from 5.8% to 10.1%, P from 5.9% to 9.2%, and S from 5.8% to 10.7%.

The lower end of the distribution was more consistent. C was the lowest amino acid in 40 of the 41 sequences, with values between 0.8% and 1.3%. The exception was A0A6J2R1N8_COTGO, where M was the lowest residue at 0.5%; C was 1.0% in that sequence (Figure 3). The global minimum was therefore M at 0.5%, followed by C at 0.8%. Other residues that remained consistently low were W (1.8–2.1%), H (1.8–3.5%), F (2.8–3.9%), and Y (2.3–4.6%). M was usually low as well, but in most sequences it was not the minimum because of occasional values as high as 3.1%.

**Figure 3.**
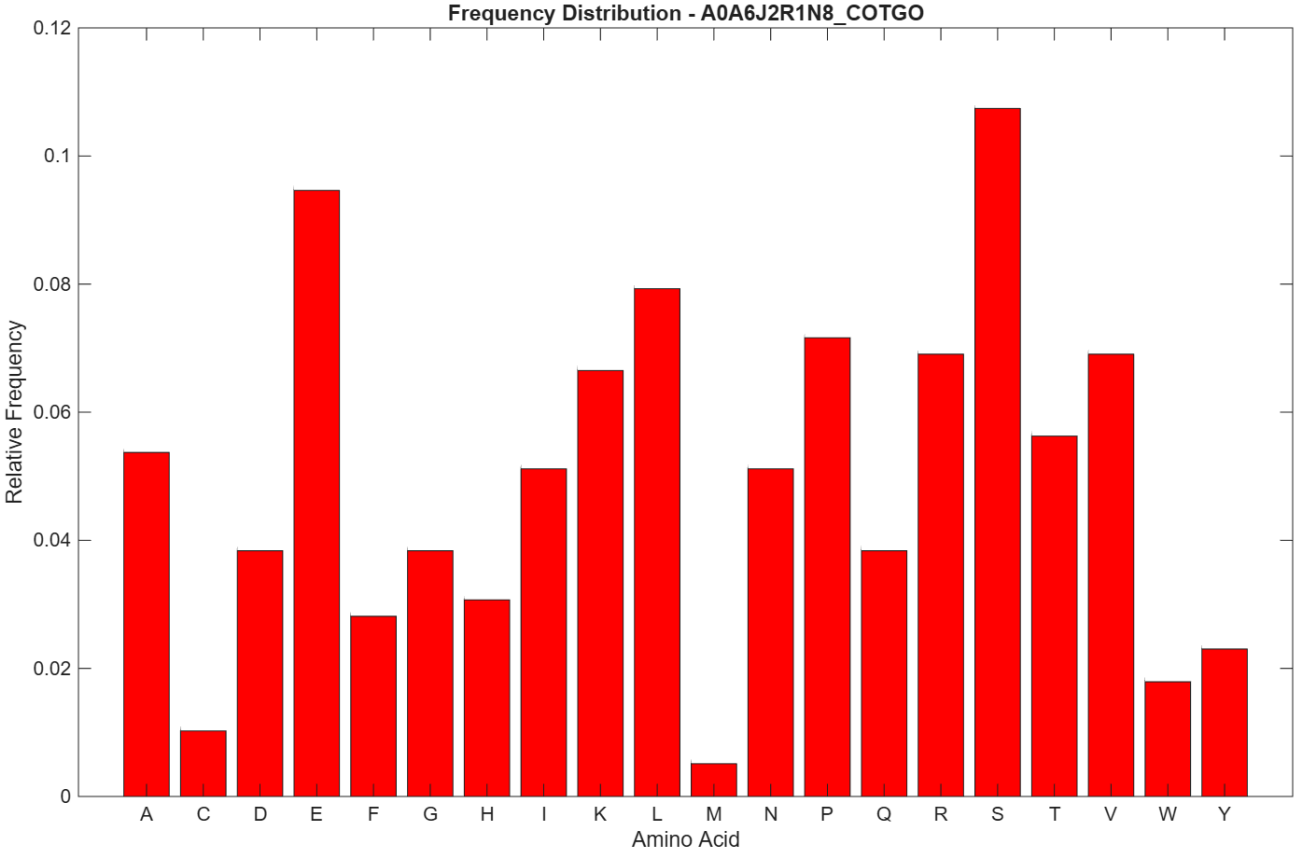
Amino acid frequency distribution in A0A6J2R1N8_COTGO.

Furthermore, it was observed that most sequences followed the same broad pattern: high R, E, K, and P, and low C. Even so, a few sequences departed from that pattern enough to merit separate comment. The finest case was A0A6J2R1N8_COTGO (Figure 3). It was the only sequence in which M, rather than C, was the lowest residue. It also had the highest S percentage in the set (10.7%), together with unusually low D, G, and Q values (all 3.8%) and low Y (2.3%), while H (3.1%) and N (5.1%) were comparatively high. A0A9Y4JI86_9TELE had the highest E percentage observed (11.8%), C at the upper observed bound (1.3%), and relatively low D (4.0%), N (2.3%), Q (3.5%), and Y (3.0%), with elevated S (8.3%); its Q value was the lowest Q value in the set. A0A6J1UCS7_9SAUR showed a co-dominant E/K profile (10.1% each), with comparatively low R (5.7%), A (3.9%), D (4.1%), and G (3.6%), and elevated I and S (7.8% each). A0A1U7S7U0_ALLSI had the lowest A value in the data set (3.6%, tied with A0A8J0Q774_XENTR), along with C at the upper observed bound (1.3%), high K (9.5%), low R (5.9%), and the lowest observed V value (4.1%). Finally, A0A6P6ED59_OCTDE contained the highest A percentage observed (8.0%), with P as its highest residue (8.2%), R and A tied at 8.0%, the lowest observed T value (3.4%), and relatively low N (2.1%).

Taken together, the NCF1 sequences did not fall into a completely divergent compositional class. They shared a clear bias toward R, E, K, and P at the high-frequency end and toward C at the low-frequency end. Within that common framework, A0A6J2R1N8_COTGO was the most distinct because of its M minimum and S maximum, while A0A9Y4JI86_9TELE, A0A6J1UCS7_9SAUR, A0A1U7S7U0_ALLSI, and A0A6P6ED59_OCTDE showed notable quantitative shifts in particular amino acid frequencies.

#### 4.1.2. sAmino acid frequency based Shannon entropy analysis of NCF1 sequences

Shannon entropy was calculated for each of the 41 NCF1 sequences from the 20 amino acid frequencies. For a 20-residue alphabet, the theoretical maximum of the Shannon entropy is 4.3219 bits. None of the NCF1 sequences reached this ideal, but all retained high entropy values, indicating that no NCF1 sequence was dominated by one or a few residues. Across the set of NCF1 sequences, Shannon entropy ranged from 4.098 to 4.198 bits, with a mean of 4.163 *±*0.018 bits and a median of 4.164 bits (Figure 4). The overall spread was narrow, and most sequences fell between about 4.15 and 4.18 bits.

**Figure 4.**
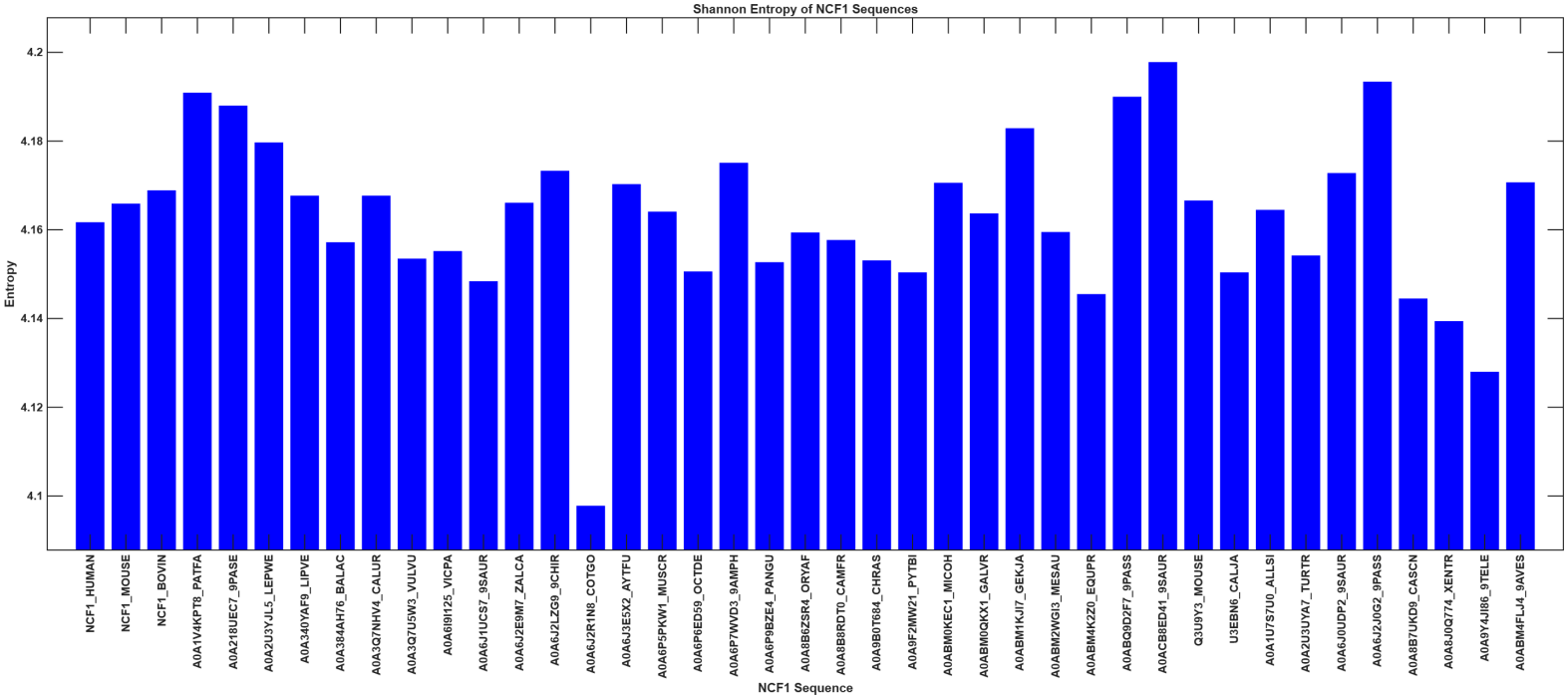
Shannon entropy profiles of NCF1 proteins across organisms.

The lowest entropy was observed in A0A6J2R1N8_COTGO (4.098 bits), the same sequence that stood out in the frequency analysis because of its M minimum (0.5%) and S maximum (10.7%) (Figure 4). The next lowest values were recorded in A0A9Y4JI86_9TELE (4.128 bits), A0A8J0Q774_XENTR (4.139 bits), A0A8B7UKD9_CASCN (4.144 bits),and A0ABM4K2Z0_EQUPR (4.146 bits). In A0A9Y4JI86_9TELE, the lower entropy was consistent with its extreme E percentage (11.8%) and comparatively low N and Q values. In A0A8B7UKD9_CASCN and A0ABM4K2Z0_EQUPR, the lower values reflected a moderate excess of R (8.2% and 8.4%, respectively) alongside several below-average frequencies. At the upper end, the highest entropy was recorded in A0ACB8ED41_9SAUR (4.198 bits), which had a comparatively even spread, with E and K tied at 8.3% and no single value exceeding that level. It was followed by A0A6J2J0G2_9PASS (4.193 bits), A0A1V4KPT8_PATFA (4.191 bits), A0ABQ9D2F7_9PASS (4.190 bits), and A0A218UEC7_9PASE (4.188 bits).

The three named NCF1 sequences, NCF1_HUMAN (4.162 bits), NCF1_MOUSE (4.166 bits), and NCF1_BOVIN (4.169 bits), all sat close to the overall mean and showed no unusual entropy signature. Overall, the Shannon entropy analysis supported the picture obtained from the frequency comparison: the NCF1 sequences share a broadly balanced amino acid composition, and the differences between them are modest and driven mainly by a small number of extreme frequencies, most notably the low M and high S values in A0A6J2R1N8_COTGO and the high E value in A0A9Y4JI86_9TELE.

#### 4.1.3. Dendrogram relationship among NCF1 sequences based on amino acid frequency distribution

A dendrogram illustrating the relationships among the 41 NCF1 sequences was constructed based on the similarity of their amino acid frequency profiles (Figure 5). Using a distance threshold cutoff of 0.4, the NCF1 sequences from various organisms were grouped into seven distinct clusters, as summarized in Table 4.

**Table 4:** List of proximal clusters based on proximity derived from amino acid frequency profile of NCF1 sequences.

| **Cluster** | **Sets of NCF1 sequences** |
|---|---|
| Cluster-1 | {NCF1_MOUSE, Q3U9YS_MOUSE, A0A6P5PKW1_MUSCR, A0ABM0KEC1_MICOH, A0ABM2WGI3_MESAU} |
| Cluster-2 | {NCF1_HUMAN, A0A6J2LZG9_9CHIR, U3EBN6_CALJA} |
| Cluster-3 | {A0A340YAF9_LIPVE, A0A384AH76_BALAC, A0ABM0QKX1_GALVR, A0A6I8I125_VICPA, A0A8B8RDT0_CAMFR, A0A2U3UYA7_TURTR, A0A8B6ZSR4_ORYAF} |
| Cluster-4 | {A0A2U3YJL5_LEPWE, A0A3Q7NHV4_CALUR, A0A6J2E9M7_ZALCA} |
| Cluster-5 | {A0A218UEC7_9PASE, A0A6J2J0G2_9PASS, A0ABQ9D2F7_9PASS} |
| Cluster-6 | {A0A6J3E5X2_AYTFU, A0ABM4FLJ4_9AVES} |
| Cluster-7 | {A0A6J1UCS7_9SAUR, A0A6P9BZE4_PANGU} |

**Figure 5.**
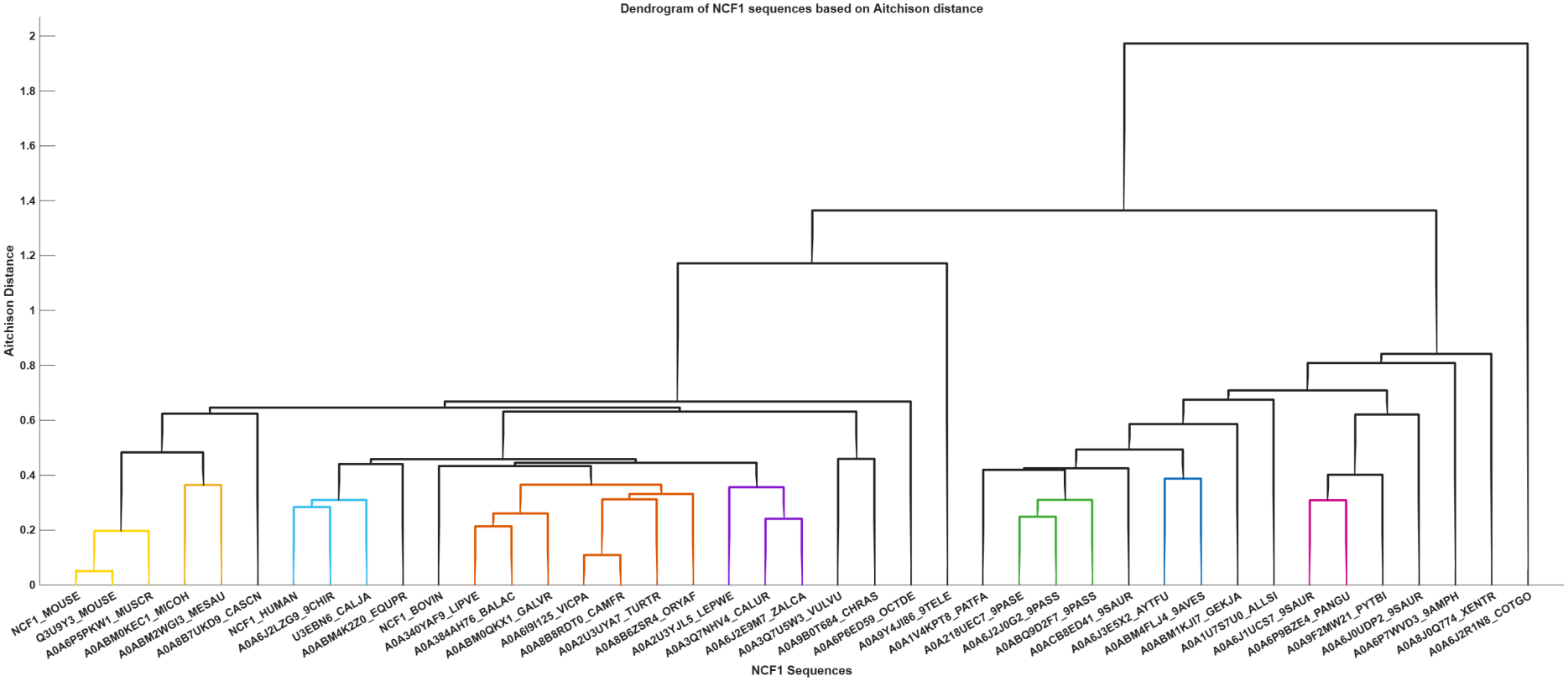
Dendrogram relationship among NCF1 sequences based on amino acid frequency distribution

### 4.2. Analysis of frequency-dominant amino acid distribution over NCF1 sequences

The 41 NCF1 sequences were scanned in consecutive 20-residue windows (window size 20 was chosen without loss of generality), and the most frequent amino acid in each window was recorded (Table 5). Window 20 (residues 381–400) was available for only 35 entries, since three entries (rows 5, 21 and 26) lacked residues beyond position 380. Each window was treated independently, and the number of entries sharing the same dominant residue was counted.

**Table 5:** Frequency dominant amino acid profiles of 41 NCF1 protein sequences based on successive 20-residue windows. NCF1 sequences sharing identical profiles were grouped together.

| **S/N** | **NCF1 Sequences** | **Window ranges** |  |  |  |  |  |  |  |  |  |  |  |  |  |  |  |  |  |  |  | **Dominant Residue** | **Windows_Dominated** | **Degree of dominance (DD)** |
|---|---|---|---|---|---|---|---|---|---|---|---|---|---|---|---|---|---|---|---|---|---|---|---|---|
|  |  | **1–20** | **21–40** | **41–60** | **61-80** | **81-100** | **101-120** | **121-140** | **141-160** | **161-180** | **181-200** | **201-220** | **221-240** | **241-260** | **261-280** | **281-300** | **301-320** | **321-340** | **341-360** | **361-380** | **381-400** |  |  |  |
| 1 | NCF1_MOUSE, A0A6P5PKW1_MUSCR, Q3U9Y3_MOUSE | F | V | E | E | G | L | D | I | A | V | E | A | E | G | Q | R | R | P | P | K | R | 2 | 10 |
| 2 | A0A6I9I125_VICPA, A0A8B8RDT0_CAMFR | F | V | E | P | A | L | D | I | A | V | E | A | E | D | Q | R | R | E | P | S | R and P | 4 | 20 |
| 3 | A0A9F2MW21_PYTBI | E | F | E | I | N | L | T | I | A | V | P | E | E | D | A | S | K | N | P | K | E | 4 | 20 |
| 4 | A0A9Y4JI86_9TELE | E | L | E | I | S | L | E | I | T | E | E | E | E | G | E | P | L | S | P | S | E | 8 | 40 |
| 5 | A0ABM1KJI7_GEKJA | E | V | E | I | S | I | P | I | A | V | E | A | E | Y | E | I | R | A | I |  | E | 5 | 26.3 |
| 6 | A0ACB8ED41_9SAUR | E | V | E | I | S | L | D | I | T | V | E | A | E | Y | E | I | R | A | L | A | E and K | 5 | 25 |
| 7 | A0A6P7WVD3_9AMPH | E | V | F | P | S | L | P | I | A | V | P | A | E | Y | K | R | R | N | P | E | K | 1 | 5 |
| 8 | A0A1V4KPT8_PATFA | F | L | E | I | Q | V | D | K | A | V | E | E | E | Y | S | R | K | N | P | R | K | 2 | 10 |
| 9 | A0A1U7S7U0_ALLSI | F | L | E | I | T | L | P | I | K | D | E | Y | E | Y | N | R | K | E | P | K | K | 3 | 15 |
| 10 | A0ABM0QKX1_GALVR | F | L | E | P | A | L | L | I | S | V | E | P | E | D | Q | R | R | P | P | S | L, R, and S | 7 | 35 |
| 11 | A0A6J3E5X2_AYTFU | F | L | E | P | Q | L | D | I | A | D | E | E | E | Y | E | R | R | E | P | R | E | 6 | 30 |
| 12 | A0ABM4FLJ4_9AVES | F | L | E | P | T | L | D | K | A | E | E | A | E | Y | S | R | R | E | P | R | E | 5 | 25 |
| 13 | A0A8B7UKD9_CASCN | F | V | E | A | T | L | D | I | A | V | E | A | E | D | Q | R | R | Q | P | T | R and S | 2 | 10 |
| 14 | A0ABM2WGI3_MESAU | F | V | E | E | T | L | D | I | A | V | E | A | E | G | A | R | R | P | P | A | R | 2 | 10 |
| 15 | A0ABM0KEC1_MICOH | F | V | E | E | T | L | D | I | A | V | E | A | E | G | Q | R | R | P | P | A | A and R | 5 | 25 |
| 16 | A0ABM4K2Z0_EQUPR | F | V | E | I | A | L | K | I | A | V | E | A | E | D | Q | R | R | P | P | A | R | 2 | 10 |
| 17 | A0A218UEC7_9PASE | F | V | E | I | A | V | D | T | A | V | E | A | E | Y | A | R | R | N | P | T | K | 0 | 0 |
| 18 | A0A2U3UYA7_TURTR | F | V | E | I | E | L | D | I | A | V | E | E | E | D | Q | R | R | E | P | A | S | 0 | 0 |
| 19 | A0A6J0UDP2_9SAUR | F | V | E | I | S | R | P | I | K | K | E | A | E | Y | N | I | R | K | P | K | E and K | 7 | 35 |
| 20 | A0A6J2LZG9_9CHIR | F | V | E | I | T | L | D | I | A | V | E | A | E | D | A | R | R | E | P | A | R | 2 | 10 |
| 21 | A0ABQ9D2F7_9PASS | F | V | E | I | T | V | D | K | A | V | E | E | E | Y | S | R | K | A | R |  | E | 4 | 21.1 |
| 22 | A0A6J2J0G2_9PASS | F | V | E | I | T | V | D | L | A | V | E | E | E | Y | P | R | Q | N | P | R | K | 0 | 0 |
| 23 | A0A6J2E9M7_ZALCA | F | V | E | P | A | L | D | I | A | K | P | E | E | D | Q | R | R | P | P | A | R | 2 | 10 |
| 24 | A0A3Q7NHV4_CALUR | F | V | E | P | A | L | D | I | A | K | P | E | E | D | Q | R | R | R | P | A | R | 3 | 15 |
| 25 | A0A6P6ED59_OCTDE | F | V | E | P | A | P | P | I | A | V | E | A | E | V | Q | R | R | P | P | A | P | 5 | 25 |
| 26 | A0A2U3YJL5_LEPWE | F | V | E | P | E | L | D | I | A | K | P | E | E | I | Q | R | R | R | G |  | R | 3 | 15.8 |
| 27 | A0A340YAF9_LIPVE | F | V | E | P | E | L | D | I | A | V | E | A | E | D | Q | R | R | E | P | S | P | 2 | 10 |
| 28 | A0A384AH76_BALAC | F | V | E | P | E | L | D | I | A | V | E | A | E | D | Q | R | R | P | P | S | P | 3 | 15 |
| 29 | NCF1_BOVIN | F | V | E | P | T | L | D | I | A | V | E | A | E | D | A | R | R | Q | P | S | R | 2 | 10 |
| 30 | U3EBN6_CALJA | F | V | E | P | T | L | D | I | A | V | E | A | E | D | Q | R | R | P | P | A | R | 2 | 10 |
| 31 | A0A9B0T684_CHRAS | F | V | E | P | T | L | D | I | A | V | E | E | E | D | A | R | R | A | P | A | R | 2 | 10 |
| 32 | A0A3Q7U5W3_VULVU | F | V | E | P | T | L | D | I | A | V | E | E | E | D | A | R | R | P | P | S | P | 3 | 15 |
| 33 | NCF1_HUMAN | F | V | E | P | T | L | D | T | A | V | E | A | E | D | Q | R | R | P | P | A | R | 2 | 10 |
| 34 | A0A8B6ZSR4_ORYAF | F | V | F | P | T | L | D | I | A | V | E | A | E | D | Q | R | R | A | P | A | R | 2 | 10 |
| 35 | A0A6J1UCS7_9SAUR | I | F | E | I | E | L | P | I | A | V | P | E | E | D | N | S | K | E | P | K | E and K | 7 | 35 |
| 36 | A0A6P9BZE4_PANGU | I | F | E | I | R | I | P | I | A | V | P | E | E | E | N | S | K | E | P | K | E | 5 | 25 |
| 37 | A0A8J0Q774_XENTR | I | V | E | P | S | L | N | I | A | E | P | D | E | G | K | K | Q | E | P | K | K | 3 | 15 |
| 38 | A0A6J2R1N8_COTGO | V | L | S | L | E | R | I | E | W | E | E | V | E | R | A | R | K | S | S | S | S | 4 | 20 |

The most striking result was window 13 (residues 241–260), where glutamic acid (E) was the dominant residue in all 38 entries (100%). This was the only completely invariant window in the set of 41 NCF1 sequences. Window 3 (41–60) came close, with E dominant in 35 of 38 entries (92.1%); the only exceptions were rows 7 and 34, where phenylalanine (F) dominated, and row 38, where serine (S) dominated. Window 19 (361–380) was dominated by proline (P) in 33 of 38 entries (86.8%), with isolated exceptions in rows 5, 6, 21, 26 and 38. Window 9 (161–180) was dominated by alanine (A) in 32 of 38 entries (84.2%), and window 8 (141–160) by isoleucine (I) in 31 of 38 entries (81.6%). Two windows shared the next rank: window 11 (201–220) was dominated by E in 30 of 38 entries (78.9%), and window 16 (301–320) by arginine (R) in 30 of 38 entries (78.9%).

A second group of windows was strongly but not overwhelmingly conserved. Window 1 (1–20) was dominated by F in 29 of 38 entries (76.3%), and window 6 (101–120) by leucine (L) in 29 of 38 entries (76.3%). Windows 2 (21–40) and 10 (181–200) were both dominated by valine (V) in 28 of 38 entries (73.7%). Window 17 (321–340) was dominated by R in 26 of 38 entries (68.4%), with E, K, L, Q and A accounting for the remaining entries. Window 7 (121–140) was dominated by aspartic acid (D) in 25 of 38 entries (65.8%), with P the next most frequent residue (7 entries).

The remaining windows were considerably more variable. Window 12 (221–240) was led by A in 20 of 38 entries (52.6%), but E was nearly as frequent (14 entries). Window 4 (61–80) was led by P in 18 of 38 entries (47.4%), with I close behind (15 entries). Window 14 (261–280) was led by D in 18 of 38 entries (47.4%), followed by tyrosine (Y) in 11 entries and glycine (G) in 5. Window 18 (341–360) was led by P in 17 of 38 entries (44.7%), with E in 8 and asparagine

(N) in 5. Window 15 (281–300) was led by glutamine (Q) in 16 of 38 entries (42.1%), with A in 8. Window 20 (381–400) was led by R in 14 of the 35 available entries (40.0%), followed by K in 7, S in 6 and P in 4. Window 5 (81–100) was the most heterogeneous of all, with threonine (T) dominant in only 14 of 38 entries (36.8%), followed by A in 7 and by S and E in 6 each.

Taken together, the window analysis revealed a clear block structure in the NCF1 sequences. The N-terminal and central regions were dominated by a relatively small set of residues—F, V, E, L, I, A and D—with windows 3, 8, 9, 11 and 13 being particularly robust. The C-terminal region was more R- and P-rich, especially in windows 16, 17 and 19, but it also contained the most variable windows in the set (windows 18 and 20). This pattern was consistent with the overall composition results: E, R and P were not only frequent across the whole protein, but also repeatedly dominated specific local stretches, whereas residues such as T, Q and P in windows 5, 15 and 18 were locally dominant only in a minority of sequences.

The degree of dominance (DD) summarised how often a sequence’s overall dominant residue also dominated its individual windows. Across the 38 entries, DD ranged from 0% to 40%, with a mean of 16.7% and a median of 15%. The highest value was in A0A9Y4JI86_9TELE, where E dominated 8 of 20 windows (40%), matching its high overall E frequency and its low Shannon entropy (Figure 6). Three entries—A0A218UEC7_9PASE, A0A2U3UYA7_TURTR, and

**Figure 6.**
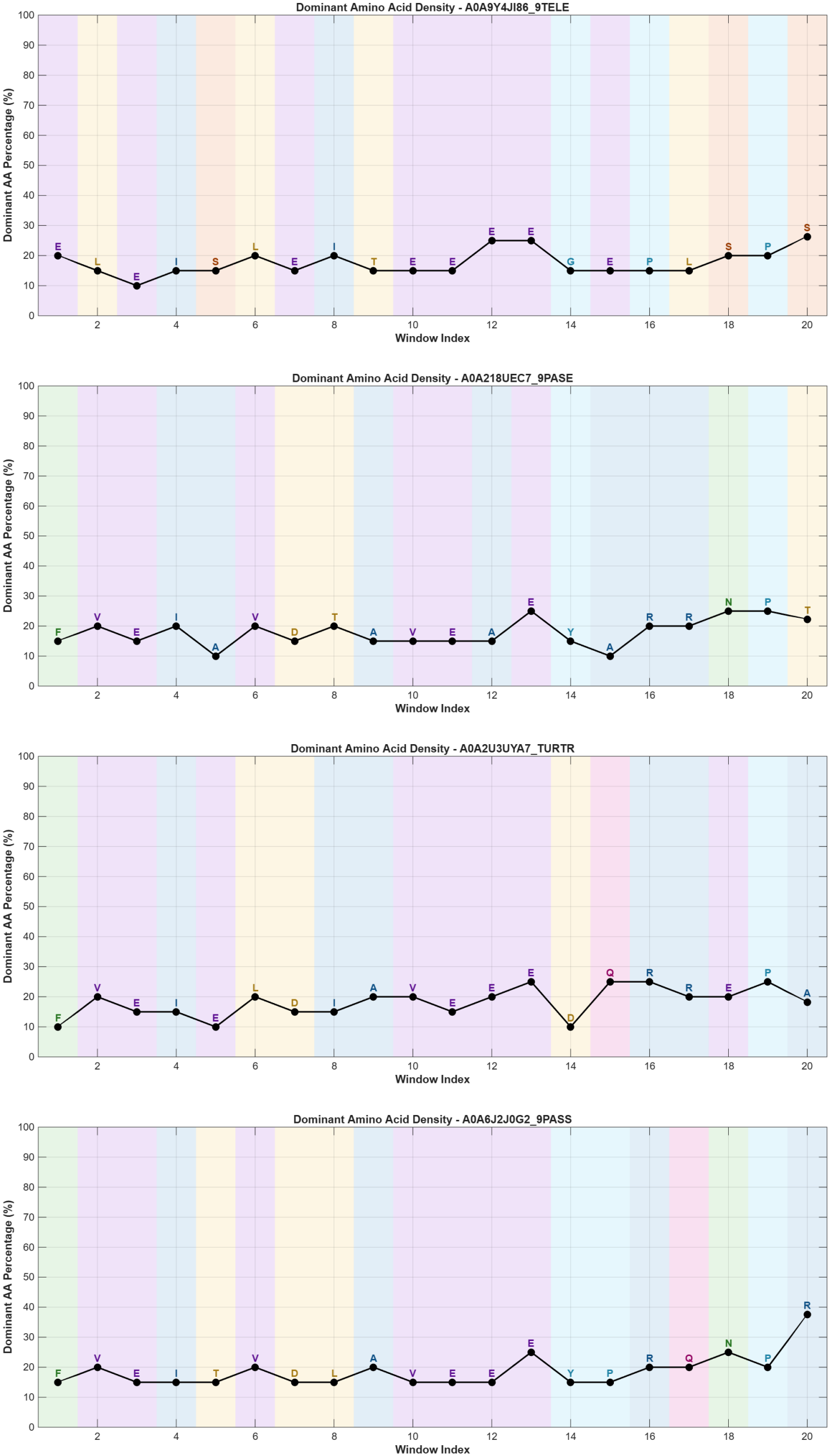
Window-wise dominance of overall frequent amino acids in four representative NCF1 sequences. Plots show the dominant amino acid percentage per 20-residue window. Purple markers indicate windows where the sequence’s overall dominant residue matched the local window dominant (contributing to the Degree of Dominance, DD), while black markers indicate alternative local dominants. The panels illustrate the spectrum of DD, from high local dominance (top) to complete absence of local dominance (bottom three, DD = 0%).

A0A6J2J0G2_9PASS—had DD = 0%, meaning that their overall dominant residue never dominated a single 20-residue window (Figure 6). This confirmed that global frequency ranking and local window dominance were related but not equivalent: a residue could be marginally the most abundant overall yet be outcompeted in every individual window by locally clustered residues. The three named NCF1 sequences (human, mouse and bovine) all showed DD = 10%, sitting at the lower end of the distribution and matching the general pattern for mammalian sequences in the set.

### 4.3. Phylogenetic relationships among NCF1 sequences based on amino acid sequence homology

#### 4.3.1. Position specific Shannon entropy profile of NCF1 protein sequences

Site-specific conservation across the NCF1 alignment was assessed using Shannon entropy (Figure 7). Rather than maintaining a uniform level of conservation, the profile showed a distinct “sawtooth” pattern, swinging between strict conservation (0 bits) and extreme variability (peaking at roughly 3.3 bits). This indicated that NCF1 contains distinct blocks of highly conserved residues interspersed with hypervariable stretches.

**Figure 7.**
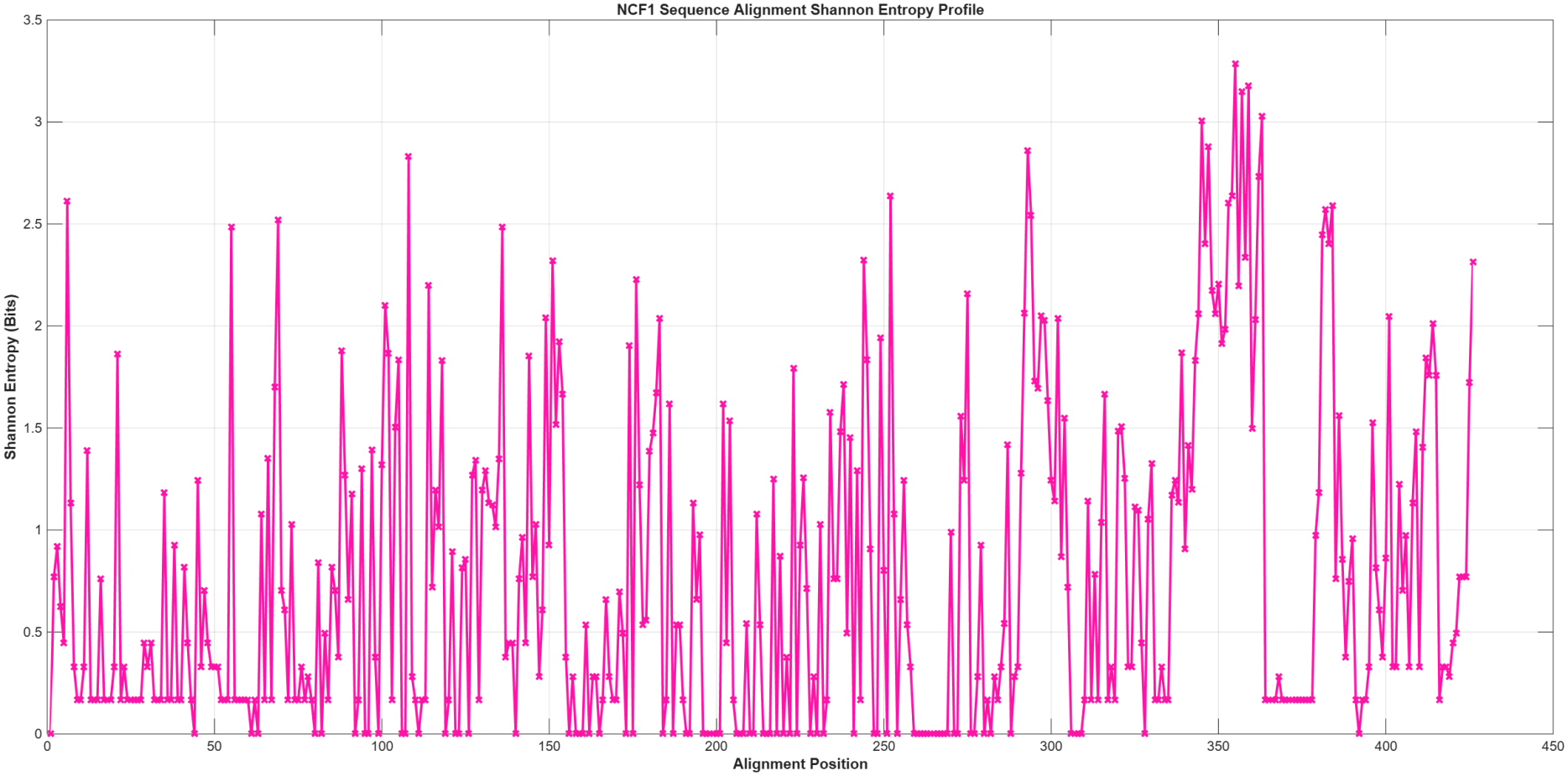
Position-specific Shannon entropy profile of the NCF1 multiple sequence alignment. The plot displayed the Shannon entropy (in bits, y-axis) calculated at each alignment position (x-axis) across the NCF1 sequences. Higher entropy values indicated greater amino acid variability and lower conservation at a given site, whereas values approaching zero reflected strict conservation.

Several regions were nearly completely conserved (approaching 0 bits), most notably the extended stretches around alignment positions 195–210, 260–285, and 365–380, as well as shorter blocks near positions 10–20, 40–50, and 70–80. These zero-entropy blocks likely represented structurally or functionally critical sites—such as the protein core or binding interfaces—maintained by strong purifying selection.

Conversely, other segments varied extensively. Regions around positions 100–110 and 240–260 showed high entropy (>2.0 bits), but the most striking feature was a massive hypervariable cluster between positions 340 and 370. Here, entropy peaked at roughly 3.3 bits, implying that nearly any of the 20 amino acids could be tolerated at these sites. This suggests a highly flexible loop, an intrinsically disordered region, or a segment under positive selection.

Overall, the entropy profile confirmed a block-like architecture for NCF1—a conserved framework punctuated by highly variable regions. This aligned well with the earlier window-dominance analysis, where certain 20-residue blocks were strictly dominated by a single residue, while others showed mixed local dominants.

#### 4.3.2. Conserved and variable residues derived from sequence logo

Overall conservation and variability of the NCF1 proteins were depicted by a sequence logo, that was generated from the multiple sequence alignment through [WebLogo](https://weblogo.threeplusone.com/create.cgi) (Figure 8). The logo confirmed a highly structured architecture, where extensive regions of strong purifying selection (approaching 4.0 bits of information) were punctuated by short, highly variable stretches (dropping below 1.0 bit). The global sequence was notably rich in basic residues, specifically lysine (K) and arginine (R), which frequently occupied highly conserved positions, consistent with the earlier global composition analysis.

**Figure 8.**
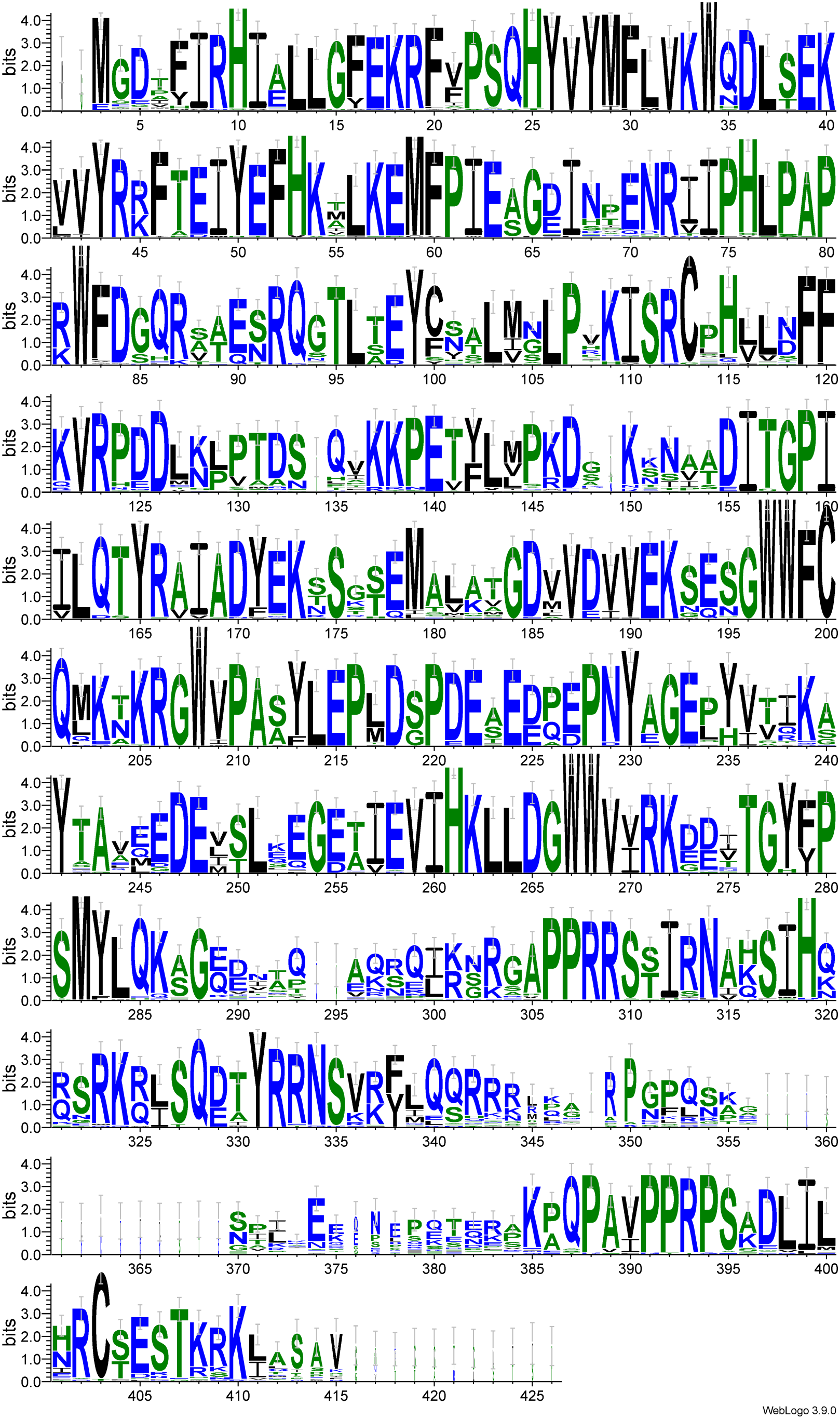
Sequence logo of the NCF1 multiple sequence alignment. The y-axis displays the information content (bits) at each alignment position, with letter height indicating the degree of sequence conservation and letter width indicating amino acid frequency. Residues are color-coded by physicochemical property (blue, basic; red, acidic; green, polar; black, hydrophobic). The logo highlights highly conserved blocks interspersed with variable regions, matching the Shannon entropy profile.

Several distinct blocks of high conservation were evident across the alignment. In the N-terminal region, strong conservation was observed at positions 16–20 (G-E-K-R-E). The central region featured highly conserved motifs, including an acidic stretch at positions 95–105 (E-G-S-A-L-M-N-L-P) and a distinct block at positions 135–140 (Q-K-K-P-E). The latter half of the protein contained particularly dense conserved clusters, such as the motif at positions 251–260 (S-L-E-G-E-T-I-E-V-I) and a basic stretch at positions 329–340 (R-S-R-K-L-S-D-T-V-R-R).

Proline (P) also emerged as a major structural determinant in the C-terminal region. A prominent proline-rich block was located at positions 385–400 (E-R-P-Q-P-A-V-P-P-R-P-S), immediately followed by a highly conserved basic and polar motif at positions 405–412 (H-R-E-S-I-K-K). These strictly conserved regions likely represented structurally or functionally critical sites, such as the protein core, binding interfaces, or interaction domains.

Despite this strong overall conservation, the logo revealed several distinct gaps where information content dropped sharply. The most prominent variable stretches occurred roughly at positions 360–375 and near the immediate C-terminus (positions 415–430). These variable regions corresponded precisely to the high-entropy peaks observed in the Shannon entropy profile, suggesting that these segments were either tolerant to substitution, intrinsically disordered, or subject to positive selection.

#### 4.3.3. Clinically significant NCF1 variants

It was know from the previous studies that 11 distinct clinical variant mutation sites across the NCF1_HUMAN protein, mostly associated with autosomal recessive Chronic Granulomatous Disease (CGD1) were identified [15, 39].

These included three nonsense mutations (Q91*, C111*, W193*), three missense substitutions (two at position 42 [R>Q and R>W] and one at G192S), and four deletions (loss of Y26, Y97, E168, and V271). While nearly all were classified as pathogenic, the R90H variant was deemed likely pathogenic and has also been linked to systemic lupus erythematosus susceptibility.

Strikingly, mapping these clinical variants onto the conservation profiles revealed a clear pattern. The vast majority of these disease-causing mutations—including the nonsense mutations at positions 91, 111, and 193, and the deletions at 97, 168, and 271—fell directly within the highly conserved, zero-entropy blocks identified in our sequence logo and Shannon entropy analyses.

The strict conservation of these specific regions implies that they were structurally or functionally indispensable. Consequently, disruptive mutations at these sites were highly intolerant and resulted in severe loss-of-function phenotypes like CGD. The clustering of pathogenic variants within these conserved blocks strongly corroborated the functional importance of the NCF1 structural core identified in this study.

#### 4.3.4. Phylogenetic relationships among NCF1 protein sequences based on amino acid sequence homology

A total of 41 NCF1 sequences were subjected to hierarchical clustering based on sequence homology, and the resulting dendrogram was cut at a distance threshold of 10 to define clusters (Figure 9). This yielded eight color-coded subclusters (Cluster-1 to Cluster-8), which largely mirrored taxonomic and phylogenetic boundaries (Table 6). The largest cluster, Cluster-1, consisted entirely of five mammalian sequences (including mouse and other small mammals), while Cluster-3 grouped six sequences from larger mammals (such as bovine and other ungulates). Primate sequences formed their own tight clade in Cluster-2, reflecting their close evolutionary proximity. The remaining mammalian lineages were distributed across Clusters 4, 5, and 6. In contrast, the non-mammalian sequences separated distinctly from the mammalian clades. Avian and reptilian species clustered together in the peripheral Clusters 7 and 8.

**Table 6:** List of cluster containing proximal NCF1 sequences from various organisms derived from amino acid sequence homology by setting a distance cutoff of 10.

| **Cluster** | **Sets of proximal NCF1 protein sequences** |
|---|---|
| Cluster-1 | {NCF1_MOUSE, Q3U9Y3_MOUSE, A0A6P5PKW1_MUSCR, A0ABM0KEC1_MICOH, A0ABM2WGI3_MESAU} |
| Cluster-2 | {NCF1_HUMAN, U3EBN6_CALJA} |
| Cluster-3 | {NCF1_BOVIN, A0A340YAF9_LIPVE, A0A384AH76_BALAC, A0A2U3UYA7_TURTR, A0A6I9I125_VICPA, A0A8B8RDT0_CAMFR} |
| Cluster-4 | {A0A8B6ZSR4_ORYAF, A0A9B0T684_CHRAS} |
| Cluster-5 | {A0A3Q7NHV4_CALUR, A0A6J2E9M7_ZALCA, A0A3Q7U5W3_VULVU} |
| Cluster-6 | {A0A1V4KPT8_PATFA, A0A6J3E5X2_AYTFU} |
| Cluster-7 | {A0A218UEC7_9PASE, A0A6J2J0G2_9PASS, A0ABQ9D2F7_9PASS} |
| Cluster-8 | {A0ACB8ED41_9SAUR, A0A6P9BZE4_PANGU, A0A9F2MW21_PYTBI} |

**Figure 9.**
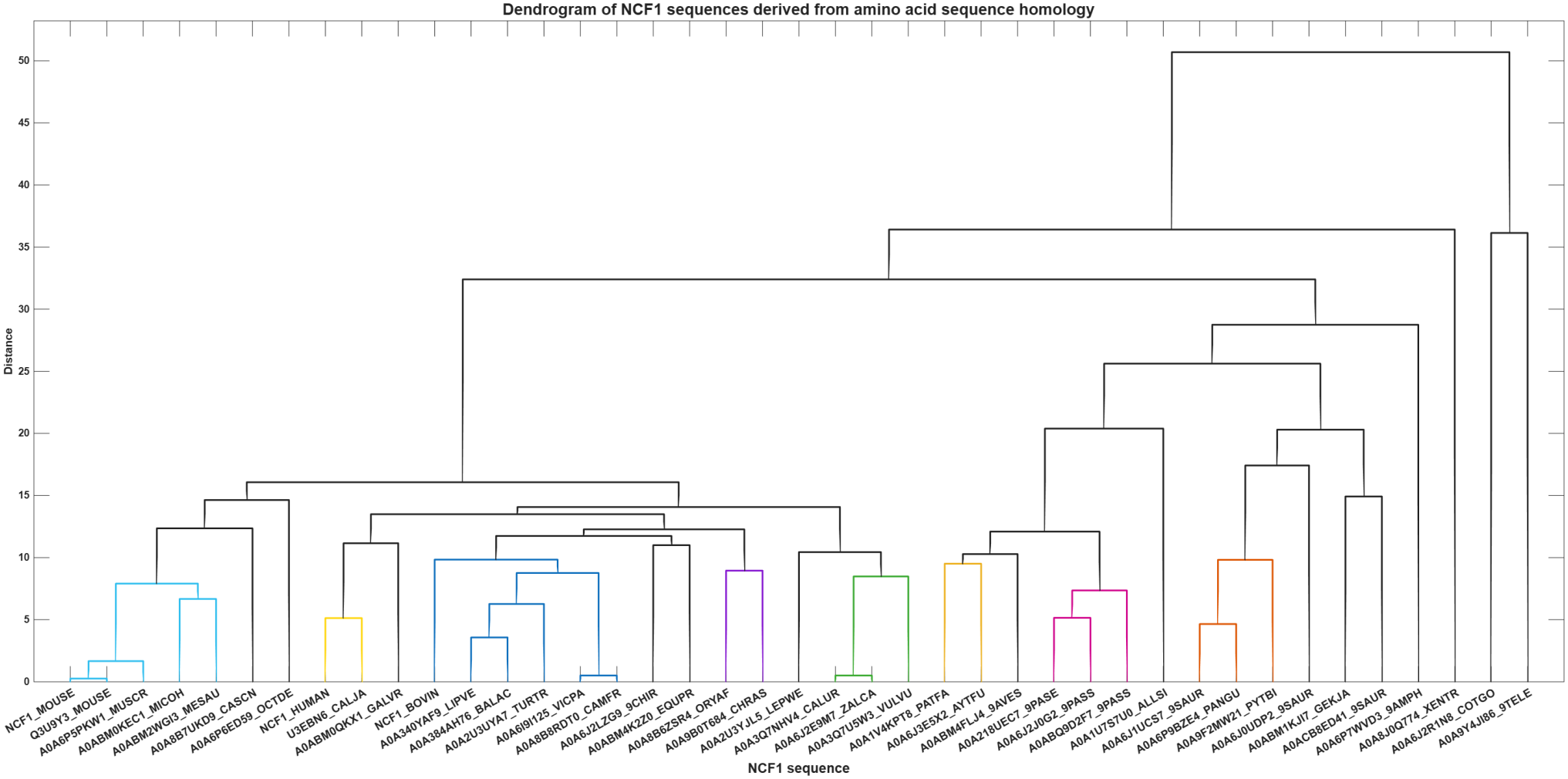
Dendrogram showing homology relationships among 41 NCF1 sequences based on amino acid sequence similarity.

### 4.4. Dendrogram relationships among NCF1 sequences based on dipeptide composition analysis

The 400 possible dipeptides were classified into five prevalence categories based on their occurrence across NCF1 protein sequences (Table 7). The results revealed a strong compositional bias that directly reflected the overall amino acid abundance. Highly prevalent dipeptides (>75%) were overwhelmingly enriched in aliphatic, acidic, and basic residues. Proline-, leucine-, and valine-rich combinations (e.g., PP, LP, LL, LV) and charged clusters (e.g., DE, EK, EP, KK, RR) dominated this category. Interestingly, specific aromatic pairs such as WW and WF were also highly prevalent, indicating localized hydrophobic packing.

**Table 7:** Prevalence categories of the 400 dipeptides across NCF1 protein sequences. Categories: highly prevalent (>75%), common (51–75%), moderately prevalent (26–50%), rare (*≤* 25%), and absent (0%).

| **Highly prevalent** | **Highly prevalent** | **Highly prevalent** | **Highly prevalent** | **Moderately prevalent** | **Common** | **Rare** | **Rare** | **Absent** |
|---|---|---|---|---|---|---|---|---|
| AP | SG | EF | NP | GS | IN | MV | IF | AC |
| CQ | SQ | GY | QG | HS | KN | NG | IG | AW |
| DE | SR | HL | TA | NF | PT | VN | KH | CC |
| DG | TG | IT | TK | NI | RW | HT | MH | CD |
| DL | TL | IY | AV | VL | AS | MN | MR | CF |
| EK | TY | KI | EA | AH | DP | RE | WI | CI |
| EP | VE | KP | GQ | DA | GA | SF | CA | CW |
| ET | VI | LI | GT | MP | LM | TN | CE | DW |
| EV | VR | LS | NA | NV | PL | TV | CG | EC |
| FC | VV | MY | SV | QL | PQ | CN | DC | FW |
| FL | WF | NY | YT | TS | PV | FQ | FR | GC |
| FP | WV | VP | FI | YI | YF | MT | FY | GM |
| GD | WW | YM | LH | ER | LN | VF | HG | HA |
| GE | YE | AQ | RI | NL | QA | AF | HP | HC |
| GP | YL | DV | SY | NT | QM | AM | LF | HH |
| GW | YR | EN | LA | QN | QQ | CY | NH | HM |
| HI | AD | EY | MA | QV | YC | DK | PC | HW |
| HK | EI | FD | MK | DF | AK | DM | WH | IC |
| IH | EM | KA | VT | GR | CS | MQ | WY | IW |
| II | ES | LT | WQ | HV | EQ | SH | YD | KC |
| IL | FF | VD | YA | IV | HQ | VG |  | LC |
| IP | FH | AE |  | KQ | RL | YS |  | LW |
| KE | HY | DD |  | NK | PY | CL |  | MC |
| KL | IA | IK |  | GL | RV | FA |  | MW |
| KR | IE | MG |  | KF | TF | HN |  | NC |
| KS | IR | QK |  | KY | TQ | PF |  | NW |
| KV | IS | QP |  | LR | VA | RD |  | RM |
| LD | KK | QR |  | NE | CP | WN |  | SC |
| LE | KW | QT |  | NN | HR | FN |  | SW |
| LG | NR | RA |  | YY | AT | FS |  | TC |
| LK | NS | SS |  | AR | SK | GG |  | TM |
| LL | PK | AI |  | CT | EL | GN |  | VW |
| LP | PN | DY |  | YP | GK | HD |  | WA |
| LQ | QH | EE |  | KM | KG | HF |  | WC |
| LV | RC | FT |  | MS | PG | NM |  | WD |
| MF | RF | KT |  | SD | QE | VH |  | WG |
| PA | RH | QI |  | TT | VS | CV |  | WK |
| PD | RQ | SA |  | ML | FV | DQ |  | WP |
| PE | SI | TD |  | RT |  | EH |  | WT |
| PH | SM | EG |  | AN |  | GH |  | YH |
| PI | ST | FE |  | ND |  | IM |  | YW |
| PP | TE | KD |  | NQ |  | LY |  |  |
| PR | VK | QD |  | PM |  | QF |  |  |
| PS | VY | QS |  | DN |  | RY |  |  |
| RG | YV | SL |  | IQ |  | CH |  |  |
| RK | AG | SP |  | MD |  | CK |  |  |
| RN | AL | TI |  | ME |  | DR |  |  |
| RP | AY | AA |  | SN |  | FM |  |  |
| RR | DI | DT |  | TP |  | GV |  |  |
| RS | DS | FK |  | TR |  | HE |  |  |
| SE | ED | GF |  | VQ |  | ID |  |  |

On the other side, the “absent” (0%) and “rare” (*≤* 25%) categories were heavily dictated by two residues: cysteine (C) and tryptophan (W). Almost all dipeptides containing cysteine in combination with other residues (e.g., AC, CC, CD, CW, IC, LC, WC) were completely absent from the dataset. Similarly, tryptophan paired with non-aromatic or non-hydrophobic residues (e.g., AW, DW, IW, LW, YW) was largely missing. This near-total absence aligned with earlier findings that C and W were the rarest residues in NCF1, which imposed strict limitations on their local sequence context.

Furthermore, eight clusters of NCF1 sequences were derived from the dendrogram relationship derived from dipeptide composition of amino acids across 41 NCF1 protein sequences (Figure 10). The resulting dendrogram revealed that the sequences largely grouped according to their biological and taxonomic lineages, indicating that dipeptide compositional constraints are strongly lineage-specific. The rodent NCF1 sequences segregated into two distinct groups: Cluster-1 contained the primary mouse sequences, while Cluster-2 grouped two other small mammals (A0ABM0KEC1_MICOH,A0ABM2WGI3_MESAU) (Table 8). Primate sequences formed their own exclusive clade in Cluster-3 (*NCF1_HUMAN, U3EBN6_CALJA*) (Table 8).

**Table 8:** List of proximal NCF1 sequences based on dipeptide frequency composition proximity.

| **Cluster** | **Sets of NCF1 sequences** |
|---|---|
| Cluster-1 | {NCF1_MOUSE, Q3U9Y3_MOUSE, A0A6P5PKW1_MUSCR} |
| Cluster-2 | {A0ABM0KEC1_MICOH, A0ABM2WGI3_MESAU} |
| Cluster-3 | {NCF1_HUMAN, U3EBN6_CALJA} |
| Cluster-4 | {A0A340YAF9_LIPVE, A0A384AH76_BALAC, A0A2U3UYA7_TURTR} |
| Cluster-5 | {A0A6I9I125_VICPA, A0A8B8RDT0_CAMFR} |
| Cluster-6 | {A0A6J2E9M7_ZALCA, A0A3Q7NGV4_CALUR} |
| Cluster-7 | {A0A6J2J0G2_9PASS, A0ABQ9D2F7_9PASS} |
| Cluster-8 | {A0A6P9BZE4_PANGU, A0A6J1UCS7_9SAUR} |

**Figure 10.**
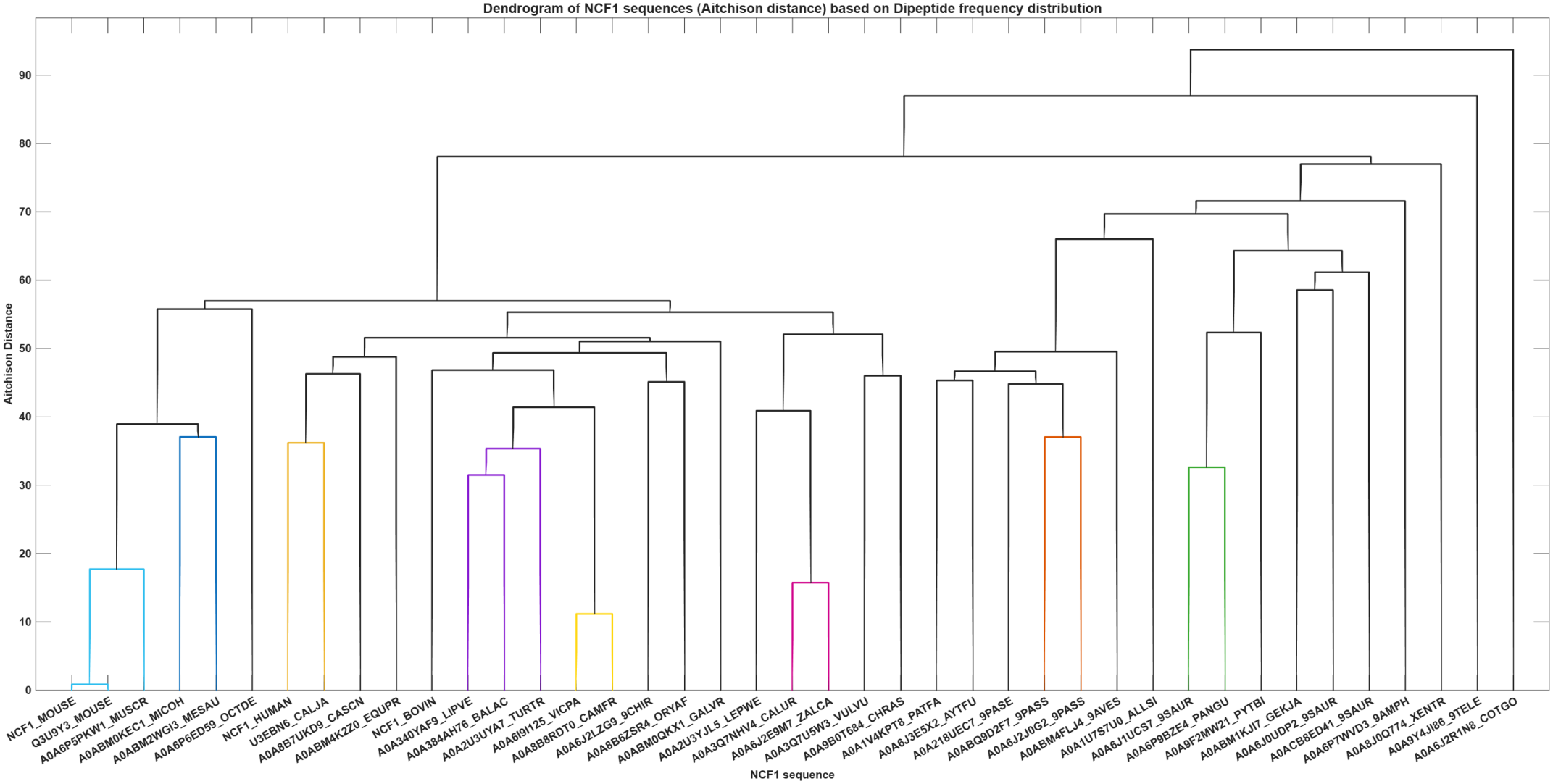
Hierarchical clustering of NCF1 sequences based on dipeptide frequencies. The dendrogram was constructed using the Aitchison distance (y-axis). Colored branches indicate distinct subclusters sharing similar dipeptide profiles, reflecting shared compositional constraints and local sequence contexts across the NCF1 family.

The remaining mammalian sequences partitioned into several clusters (Clusters 4, 5, and 6), reflecting moderate divergence in their local dipeptide usage, while the non-mammalian sequences diverged more sharply, occupying the terminal branches of the tree. Birds clustered together in Cluster-7 (A0A6J2J0G2_9PASS, A0ABQ9D2F7_9PASS),while Cluster-8 grouped a pangolin and a reptile (A0A6P9BZE4_PANGU, A0A6J1UCS7_9SAUR) (Table 8).

### 4.5. Dendrogram relationships across NCF1 sequences derived from the longest polystring analysis

The longest continuous homopolymer stretch was determined for each of the 20 standard amino acids across the 41 NCF1 sequences. Overall, the data revealed a stringent global restriction against extended repetitive tracts, with most residues rarely forming consecutive runs longer than two residues. Cysteine (C) and histidine (H) were the most tightly constrained, never forming consecutive runs in any sequence, while methionine (M) occurred consecutively only once (a run of two in A0A6J1UCS7_9SAUR). Most other residues—including A, D, F, G, N, T, Y, V, I, and W—were also capped at a maximum run length of two across the entire dataset.

Despite this strong global constraint, a few sequences exhibited distinct, localized extensions. The most striking outlier was A0A6J2R1N8_COTGO, which harbored the longest uninterrupted stretch in the dataset: a five-residue serine tract. This sequence also uniquely paired a three-residue proline tract with a phenylalanine maximum of just one residue. Another notable outlier, A0A340YAF9_LIPVE, contained the dataset’s only four-residue arginine tract. Additionally, A0A6J1UCS7_9SAUR stood out as the sole sequence capable of forming a two-residue methionine run while also containing three-residue tracts of both lysine and leucine. On the other side, several sequences displayed unusually compact homopolymer profiles. For instance, A0A2U3YJL5_LEPWE completely avoided repeats for both alanine and glutamic acid (maximum run length of one). Similarly, A0ABM1KJI7_GEKJA stood out as one of the rare sequences where serine never occurred consecutively.

These constraints became even clearer when the maximum run lengths were normalized to sequence length and plotted across all 41 sequences (Figure 11). Most amino acids clustered tightly at normalized lengths of 0.25% or 0.50%, which corresponded directly to the one- and two-residue limits. The previously noted extremes were visible as distinct upper outliers: serine and arginine reached approximately 1.27% and 1.01% (which reflected the five- and four-residue tracts), respectively, while glutamic acid, lysine, leucine, and proline peaked near 0.76% (three-residue runs). Other residues, such as asparagine, glutamine, threonine, and tyrosine, showed wider interquartile ranges, which reflected sequence-specific variability between single and double runs.

**Figure 11.**
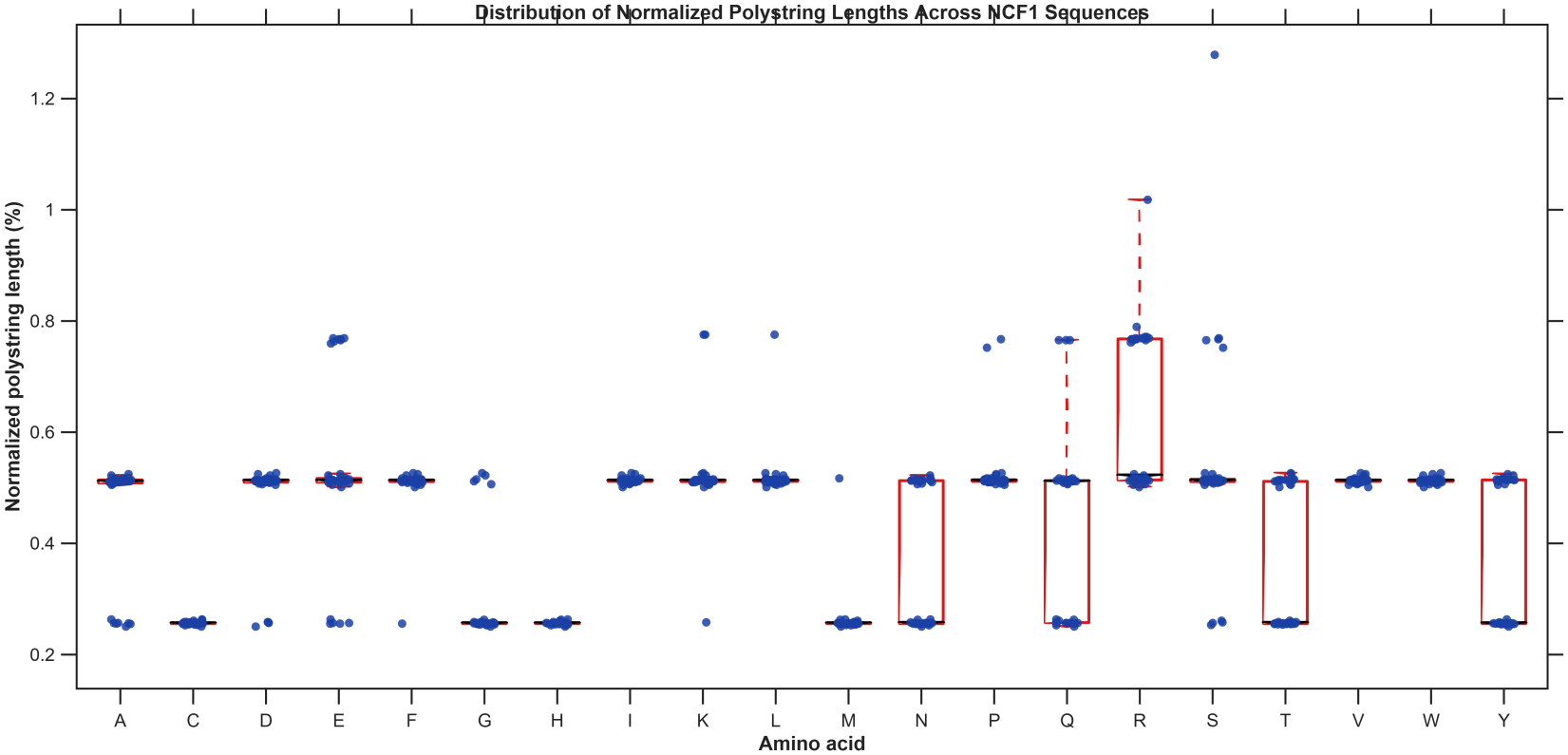
Distribution of normalized maximum homopolymer run lengths across NCF1 sequences. Boxplots display the longest consecutive run of each amino acid as a percentage of total sequence length. Most residues cluster tightly at 1- and 2-residue runs (0.25–0.50%), reflecting strong constraints against repeats. Notable outliers include serine and arginine, which form tracts up to 5 and 4 residues, respectively.

Consequently, this tight clustering reinforced the presence of strong purifying selection across the NCF1 family. Because extended homopolymers could promote protein aggregation, misfolding, or replication slippage, keeping these tracts exceptionally short likely helped maintain the compact, functional structural core of the protein. The rare, extended tracts observed in sequences like A0A6J2R1N8_COTGO may therefore have represented localized structural adaptations or intrinsically disordered regions.

Hierarchical clustering was performed using Euclidean distance based on the normalized polystring lengths to evaluate how constraints on homopolymer tracts varied across the NCF1 family (Figure 12). The resulting dendrogram partitioned the proximal NCF1 sequences into seven distinct, color-coded clusters based on a threshold 0.3 (Table 9). The clustering pattern revealed distinct groupings based on how sequences handled localized repetitive stretches. Notably, Cluster-1 grouped the human and primate sequences with two other mammalian lineages, while Cluster-2 tightly grouped several rodent-associated sequences.

**Table 9:** Clusters of NCF1 protein sequences derived from normalized polystring lengths. The seven clusters were generated via hierarchical clustering (Euclidean distance), grouping sequences based on their maximum homopolymer run length profiles.

| **Cluster** | **Sets of NCF1 sequences** |
|---|---|
| Cluster-1 | {NCF1_HUMAN, U3EBN6_CALJA, A0A384AH76_BALAC, A0A340YAF9_LIPVE} |
| Cluster-2 | {NCF1_MOUSE, A0A6P5PKW1_MUSCR, A0ABM2WGI3_MESAU, A0ABM4K2Z0_EQUPR, Q3U9Y3_MOUSE} |
| Cluster-3 | {A0A6I9I125_VICPA, A0A8B6ZSR4_ORYAF, A0ABM0QKX1_GALVR} |
| Cluster-4 | {A0A8B7UKD9_CASCN, A0A6J2LZG9_9CHIR} |
| Cluster-5 | {A0A1V4KPT8_PATFA, A0A2A8UEC7_9PASE, A0ABM4FLJ4_9AVES, A0A6J2J0G2_9PASS, A0ACB8ED41_9SAUR, A0ABQ9D2F7_9PASS} |
| Cluster-6 | {A0A6P9BZE4_PANGU, A0A6J0UDP2_9SAUR} |
| Cluster-7 | {A0A3Q7NHV4_CALUR, A0A62E9M7_ZALCA} |

**Figure 12.**
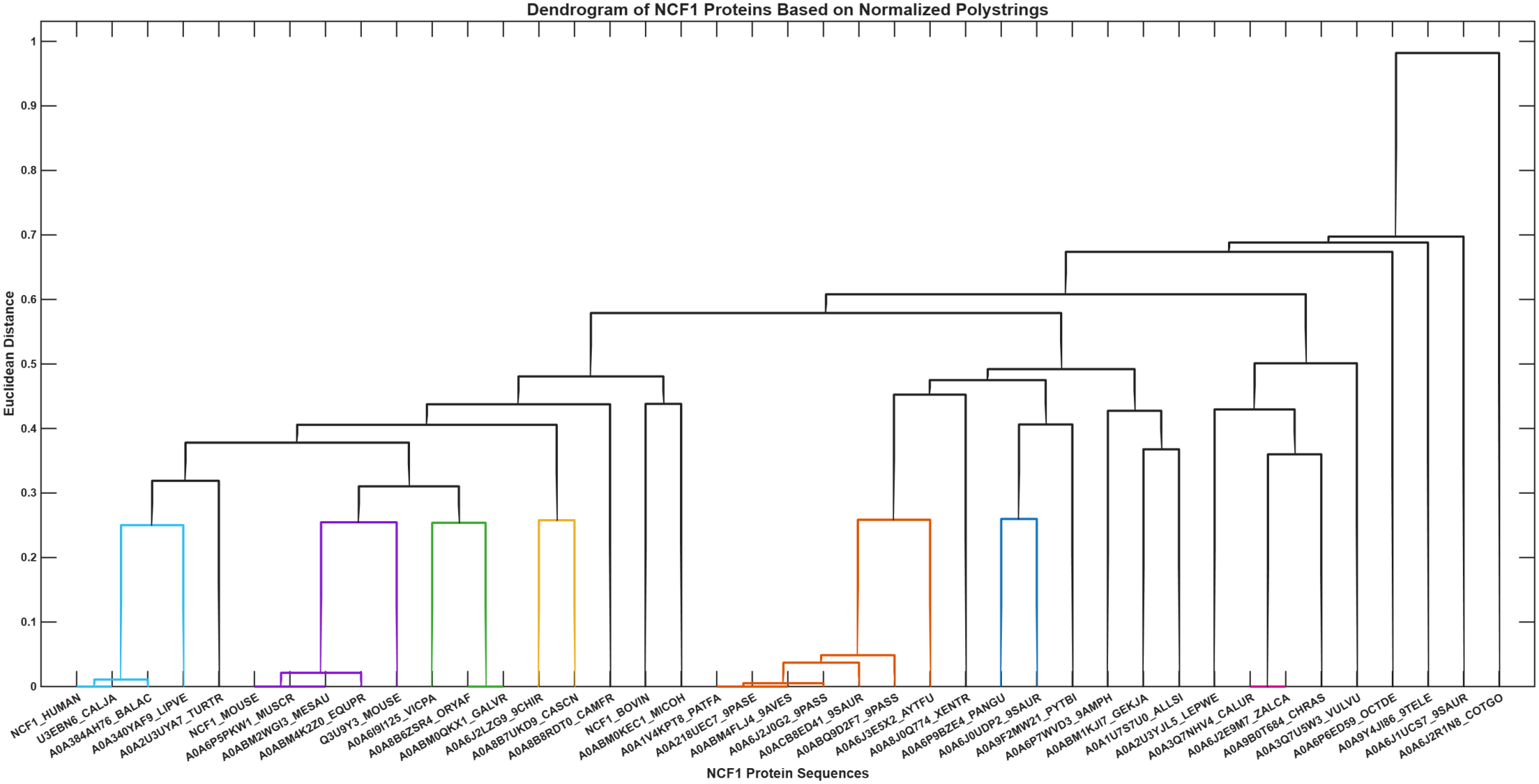
Hierarchical clustering dendrogram of NCF1 proteins based on normalized polystring lengths. The tree was constructed using Euclidean distance (y-axis) to group the 41 NCF1 sequences (x-axis) according to their maximum homopolymer run length profiles. Distinctively colored branches indicate subclusters of sequences that share highly similar constraints on repetitive tracts. This clustering highlights how sequence-specific variations in localized extensions (such as extended serine or arginine runs) group specific NCF1 proteins together.

### 4.6. Analysis of hydropathy-based n-gram frequency across NCF1 sequences

#### 4.6.1. Relative frequency of 2-grams and associated phylogenetic relationships of NCF1 sequences

Local organization of hydrophobic and hydrophilic residues was captured by translating the 41 NCF1 sequences into binary strings (Polar = 1; Non-polar = 0) and calculating the frequencies of the four possible 2-grams. The resulting distributions, visualized as violin plots, revealed a clear overarching bias across the family (Figure 13). The polar-polar 2-gram (11) dominated the local sequence contexts, ranging from 27.4% to 37.9%. Mixed pairs (01 and 10) consistently formed the second most frequent category (21.6% to 25.7%), while the hydrophobic-hydrophobic 2-gram (00) was universally suppressed (17.0% to 24.6%). This consistent hierarchy indicated that NCF1 sequences strongly favored adjacent polar residues, while strictly avoiding extended non-polar stretches—a hallmark of soluble, well-folded proteins.

**Figure 13.**
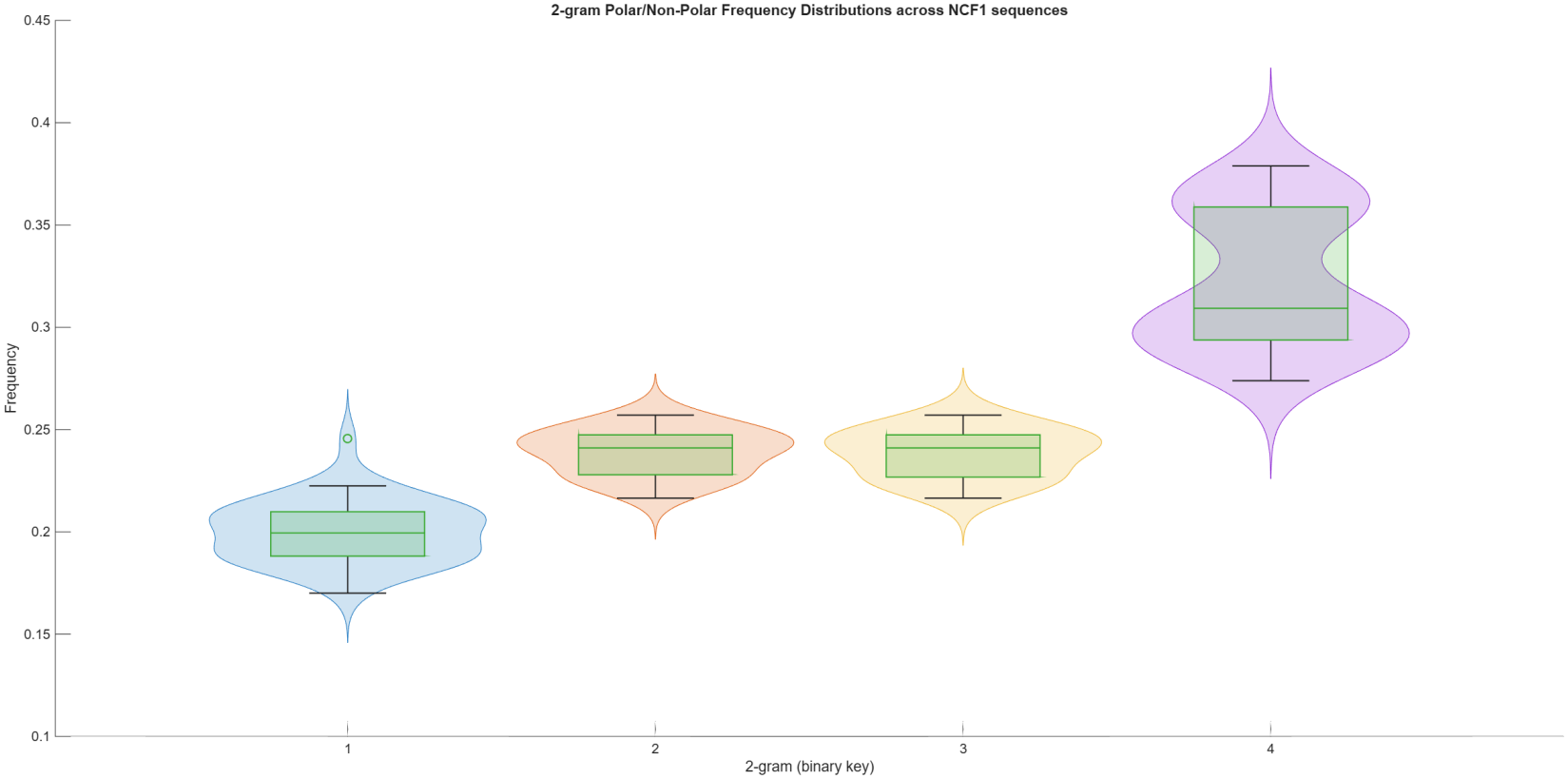
Violin plots overlaid with boxplots displaying the distribution of the four possible polar/non-polar 2-gram frequencies (binary keys 1–4) across the 41 NCF1 sequences.

Despite this shared global trend, a few NCF1 sequences displayed distinctly skewed profiles. The most polar-biased sequence was A0ABM4FLJ4_9AVES, which possessed the highest proportion of 11-grams (37.9%) and the lowest proportion of mixed pairs (21.6%), while A0A6P6ED59_OCTDE emerged as a unique outlier. It harbored the highest frequency of hydrophobic 00-grams (24.6%) and was the only sequence in the dataset, where the polar-polar 11-gram dropped below 28% (27.4%), suggesting a notably more hydrophobic local architecture. Meanwhile, A0A6P7WVD3_9AMPH exhibited the strongest avoidance of hydrophobic pairs, with the dataset’s lowest 00-gram frequency (17.0%).

Most notably, A0ABM0KEC1_MICOH and A0ABM2WGI3_MESAU shared perfectly identical 2-gram distributions. The same was true for NCF1_MOUSE and Q3U9Y3_MOUSE, which grouped tightly together. This indicated that while the entire NCF1 family maintained a strict polar bias, the exact balance of hydropathic patterns was highly conserved among closely related species.

Hierarchical clustering of the 41 NCF1 sequences based on their 2-gram frequency profiles revealed a distinct phylogenetic signature, partitioning the proteins into nine subclusters at a defined distance threshold 0.010 (Figure 14). This grouping demonstrated that the precise balance of polar and non-polar local stretches remained highly conserved within specific evolutionary lineages.

**Figure 14.**
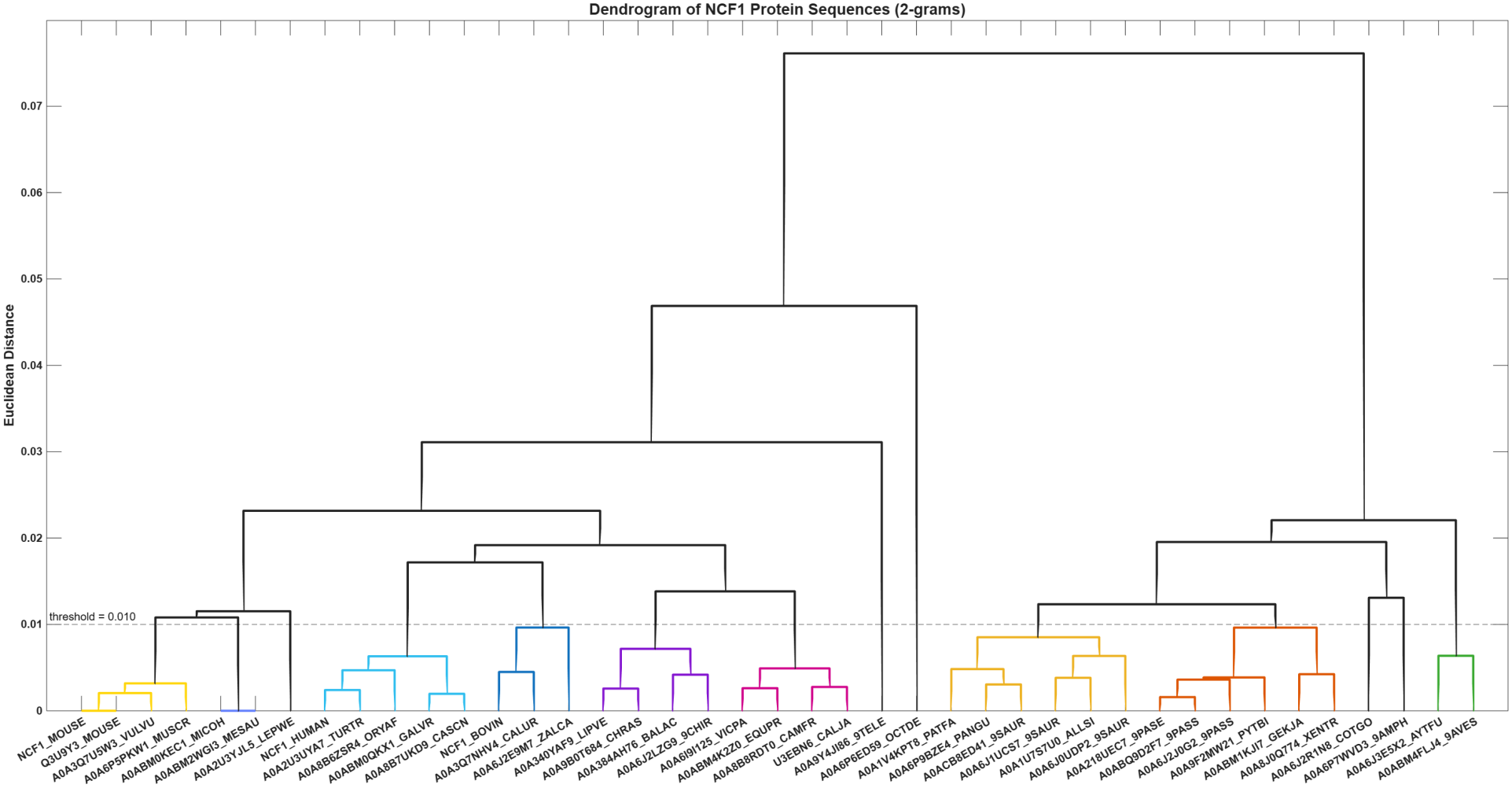
Hierarchical clustering dendrogram constructed using Euclidean distance based on the 2-gram frequency profiles. The horizontal dashed line indicates the distance threshold (0.010) used to delineate the distinct sequence subclusters, which are highlighted by different colored branches.

Clusters 1 and 2 were largely dominated by small mammals and rodents. The tight grouping in Cluster 2 was particularly notable, as it contained A0ABM0KEC1_MICOH and A0ABM2WGI3_MESAU—two sequences that shared nearly identical 2-gram distributions. Similarly, Cluster 1 grouped the primary mouse sequences (NCF1_MOUSE and Q3U9Y3_MOUSE) alongside other closely related rodent lineages (Table 10). Clusters 3 through 6 comprised a diverse array of larger mammals, including primates, ungulates, and marine mammals. Despite their broader taxonomic spread, these sequences clustered together because they maintained a highly similar proportion of polar-polar (11) and hydrophobic-hydrophobic (00) motifs.

**Table 10:** Sequence clusters derived from hydropathy 2-gram frequencies. The table lists the nine subclusters of NCF1 sequences generated via hierarchical clustering, grouping proteins that share highly similar polar/non-polar local sequence patterns.

| **Cluster** | **Sets of Proximal NCF1 sequences** |
|---|---|
| Cluster-1 | {NCF1_MOUSE, Q3U9Y3_MOUSE, A0A3Q7U5W3_VULVU, A0A6P5PKW1_MUSCR} |
| Cluster-2 | {A0ABM0KEC1_MICOH, A0ABM2WGI3_MESAU} |
| Cluster-3 | {NCF1_HUMAN, A0A2U3UYA7_TURTR, A0A8B6ZSR4_ORYAF, A0ABM0QKX1_GALVR, A0A8B7UKD9_CASCN} |
| Cluster-4 | {NCF1_BOVIN, A0A3Q7NHV4_CALUR, A0A6J2E9M7_ZALCA} |
| Cluster-5 | {A0A340YAF9_LIPVE, A0A384AH76_BALAC, A0A9B0T684_CHRAS, A0A6J2LZG9_9CHIR} |
| Cluster-6 | {A0A6I9I125_VICPA, A0ABM4K2Z0_EQUPR, A0A8B8RDT0_CAMFR, U3EBN6_CALJA} |
| Cluster-7 | {A0A1V4KPT8_PATFA, A0A6P9BZE4_PANGU, A0ACB8ED41_9SAUR, A0A6J1UCS7_9SAUR, A0A1U7S7U0_ALLSI, A0A6J0UDP2_9SAUR} |
| Cluster-8 | {A0A218UEC7_9PASE, A0ABQ9D2F7_9PASS, A0A6J2J0G2_9PASS, A0A9F2MW21_PYTBI, A0ABM1KJI7_GEKJA, A0A8J0Q774_XENTR} |
| Cluster-9 | {A0A6J3E5X2_AYTFU, A0ABM4FLJ4_9AVES} |

The most striking separation occurred in Clusters 7, 8, and 9, which distinctly grouped the non-mammalian sequences (reptiles, birds, and amphibians) away from the mammalian clades. These clusters captured the extreme hydropathy profiles identified earlier. For instance, Cluster 9 uniquely grouped A0A6J3E5X2_AYTFU with A0ABM4FLJ4_9AVES, the latter being the most highly polar-biased NCF1 sequence, while, Cluster 7 contained A0A6J1UCS7_9SAUR, which exhibited the strongest avoidance of hydrophobic pairs (Table 10).

Consequently, the clustering reinforced that while all NCF1 sequences maintained a strict polar-biased structural framework, the specific distribution of hydropathic n-grams served as a reliable fingerprint for evolutionary divergence.

#### 4.6.2. Relative frequency of 3-grams and associated phylogenetic relationships of NCF1 sequences

Extending the same binary encoding to the eight possible 3-grams revealed a clear overarching bias across the family (Figure 15). The all-polar 3-gram (111) dominated the local sequence contexts, ranging from 14.7% to 25.3% and ranking first in 40 of the 41 NCF1 sequences. The mixed 3-grams carrying two polar and one non-polar residue (011, 110 and 101) consistently formed the second most frequent category, with near-identical means (13.55%, 13.53% and 13.51%) and a combined range of 11.6% to 15.1%. The mixed 3-grams carrying one polar and two non-polar residues (001, 010 and 100) occupied the next tier (means 10.42%, 10.35% and 10.34%; range 7.8% to 12.4%), while the all-hydrophobic 3-gram (000) was universally suppressed, ranging from 7.6% to 12.7% and ranking last, alone or tied, in 31 NCF1 sequences. This consistent hierarchy indicated that NCF1 sequences strongly favoured runs of adjacent polar residues, while avoiding extended non-polar stretches—a hallmark of soluble, well-folded proteins. The violin plots further showed that the intermediate 3-grams were tightly and unimodally distributed, whereas 111 and 010 were each bimodal, splitting the family into two groups.

**Figure 15.**
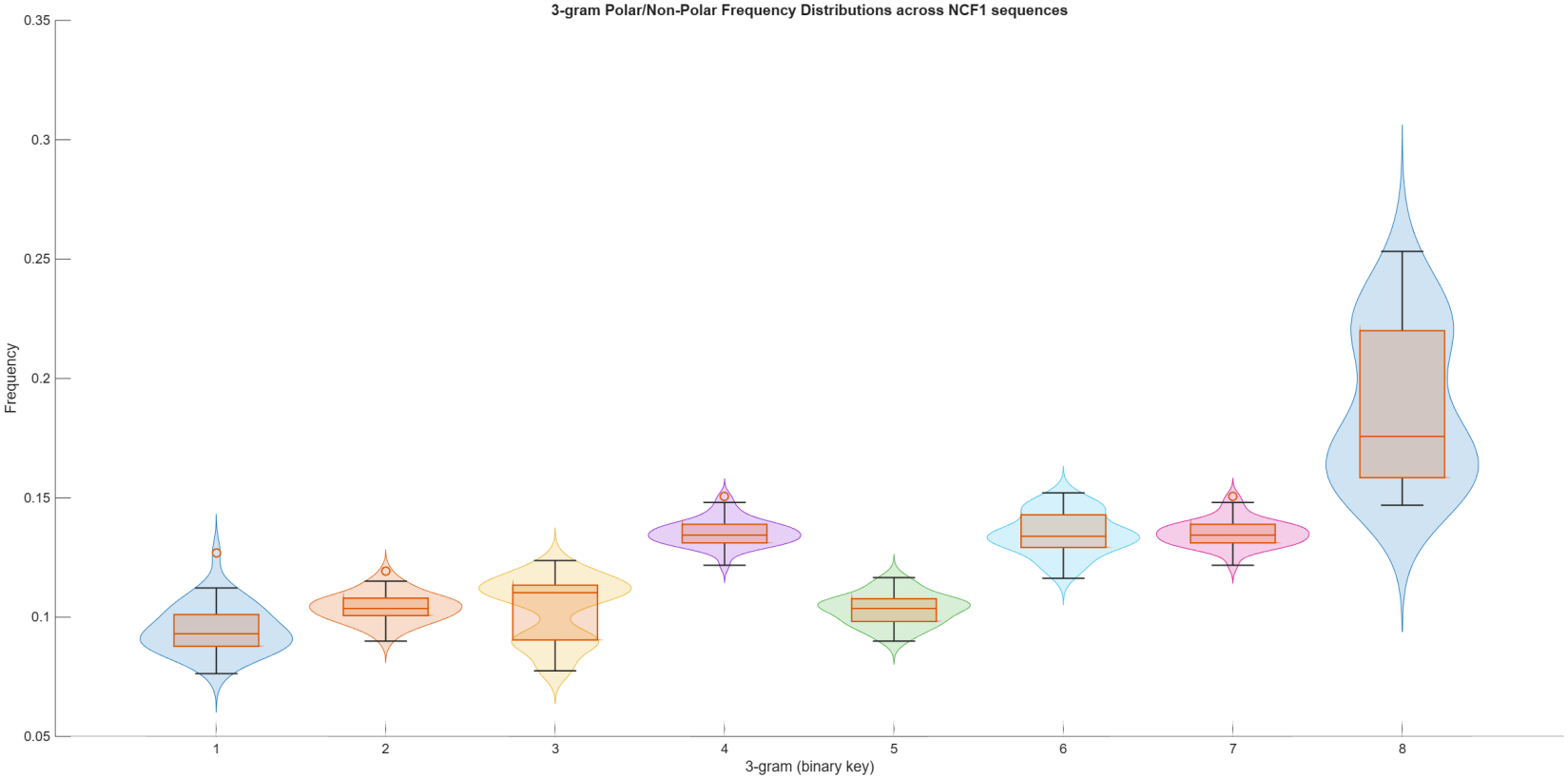
Violin plots overlaid with boxplots displaying the distribution of the four possible polar/non-polar 3-gram frequencies (binary keys 1–8) across the 41 NCF1 sequences.

Even so, a few NCF1 sequences departed noticeably from this shared trend and showed distinctly skewed 3-gram profiles. The most polar-biased sequence was A0ABM4FLJ4_9AVES, which possessed the highest proportion of all-polar 111-grams (25.3%) and the lowest proportion of 101-grams (11.6%). A0A6P6ED59_OCTDE emerged as a unique outlier: it harboured the highest frequencies of all-hydrophobic 000-grams (12.7%) and of the 001 and 100 mixed 3-grams (11.9% and 11.7%), and it showed the narrowest spread between its most and least frequent 3-grams, suggesting a notably less polar local architecture. Meanwhile, A0A6P7WVD3_9AMPH exhibited the strongest avoidance of hydrophobic runs, with the dataset’s lowest 000-gram frequency (7.6%). A0ABM0KEC1_MICOH was the only sequence in which the all-polar 111-gram was not the most frequent, being narrowly outranked by the 101-gram (15.2% vs 15.0%), and it also carried the highest proportion of 010-grams (12.4%). Most notably, only NCF1_MOUSE and Q3U9Y3_MOUSE retained perfectly identical 3-gram distributions. The pairs that had been identical at the 2-gram level (A0ABM0KEC1_MICOH with A0ABM2WGI3_MESAU, and A0A6P5PKW1_MUSCR with NCF1_MOUSE) diverged once triplet context was resolved, indicating that the finer 3-gram resolution distinguished sequences that 2-grams could not. This indicated that while the entire NCF1 family maintained a strict polar bias, the exact balance of hydropathic patterns was highly conserved among closely related species.

The relationships between the local hydropathic patterns were further explored by hierarchically clustering the 41 NCF1 sequences based on the Euclidean distance of their normalized 3-gram frequencies (Figure 16). The resulting dendrogram separated the sequences into two major branches, which resolved into nine distinct clusters below a distance threshold of 0.015 (Table 11). The largest cluster grouped seven sequences, predominantly from rodents and ungulates, while Cluster-4 strictly paired the human and bovine orthologs (NCF1_HUMAN and NCF1_BOVIN). A distinct sub-branch on the right side of the dendrogram contained the sequences with the most pronounced polar bias (Clusters 7, 8, and 9). Notably, A0A6P6ED59_OCTDE emerged as a distinct singleton, reflecting its unique, less polar 3-gram profile. Overall, the clustering demonstrated that the exact balance of local hydropathic patterns was highly conserved among closely related species and clearly segregated sequences with skewed profiles from the rest of the family.

**Table 11:** Sequence clusters derived from hydropathy 3-gram frequencies. The table lists the nine subclusters of NCF1 sequences generated via hierarchical clustering, grouping proteins that share highly similar polar/non-polar local sequence patterns.

| **Cluster** | **Sets of Proximal NCF1 sequences** |
|---|---|
| Cluster-1 | {NCF1_MOUSE, Q3U9Y3_MOUSE, A0A3Q7U5W3_VULVU, A0A6P5PKW1_MUSCR, A0A2U3YJL5_LEPWE, A0A9B0T684_CHRAS, A0ABM0KEC1_MICOH} |
| Cluster-2 | {A0A384AH76_BALAC, A0ABM4K2Z0_EQUPR} |
| Cluster-3 | {A0A6I9I125_VICPA, A0A8B8RDT0_CAMFR, A0A6J2LZG9_9CHIR, U3EBN6_CALJA} |
| Cluster-4 | {NCF1_HUMAN, NCF1_BOVIN} |
| Cluster-5 | {A0A8B6ZSR4_ORYAF, A0A2U3UYA7_TURTR, A0ABM0QKX1_GALVR, A0A8B7UKD9_CASCN} |
| Cluster-6 | {A0A3Q7NHV4_CALUR, A0A6J2E9M7_ZALCA} |
| Cluster-7 | {A0A1V4KPT8_PATFA, A0A1U7S7U0_ALLSI, A0A6J0UDP2_9SAUR, A0ACB8ED41_9SAUR} |
| Cluster-8 | {A0A9F2MW21_PYTBI, A0ABM1KJI7_GEKJA, A0A218UEC7_9PASE, A0ABQ9D2F7_9PASS, A0A6J2J0G2_9PASS} |
| Cluster-9 | {A0A6J1UCS7_9SAUR, A0A6P9BZE4_PANGU, A0A6J3E5X2_AYTFU, A0ABM4FLJ4_9AVES} |

**Figure 16.**
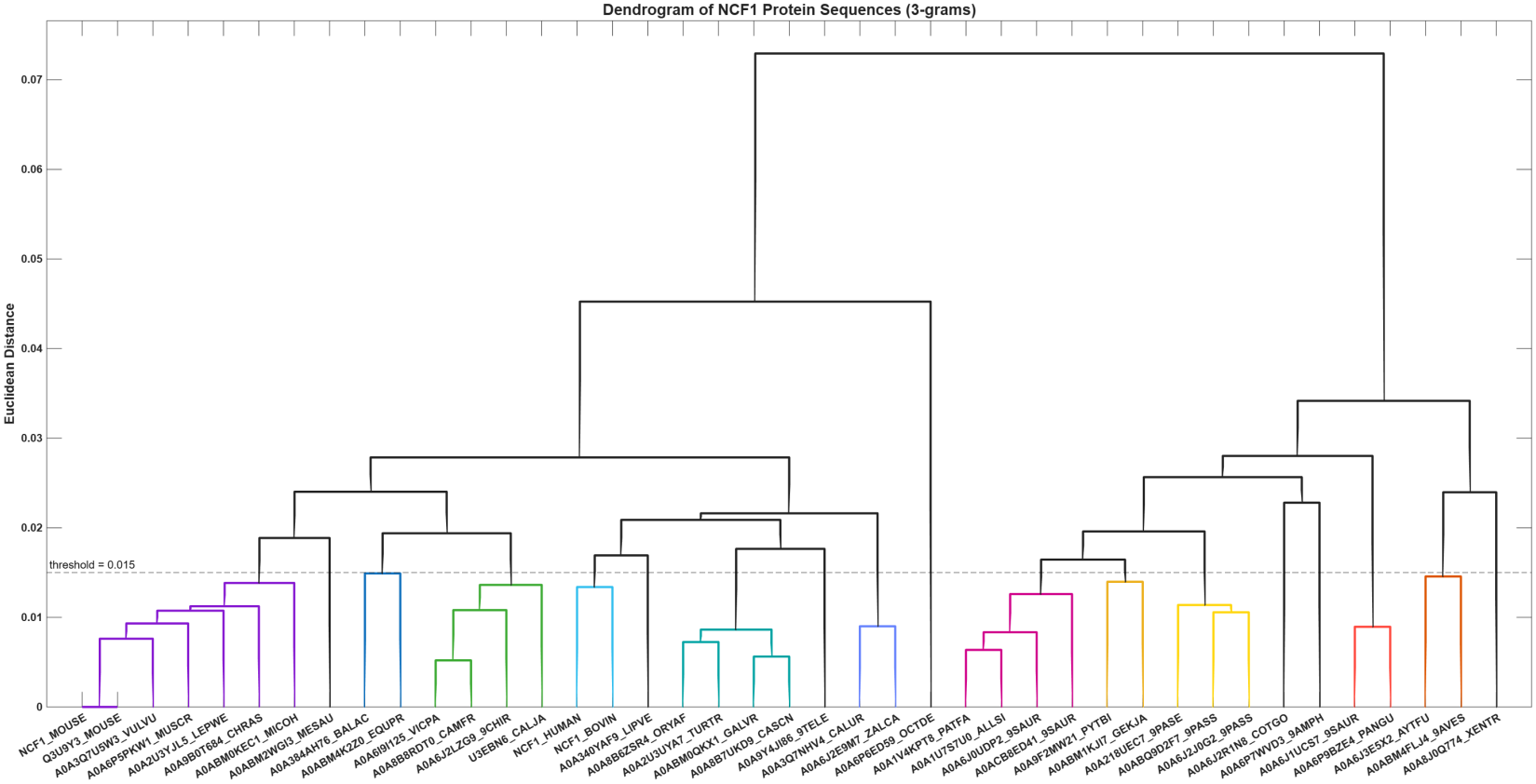
Hierarchical clustering dendrogram constructed using Euclidean distance based on the 3-gram frequency profiles. The horizontal dashed line indicates the distance threshold (0.015) used to delineate the distinct sequence subclusters, which are highlighted by different colored branches.

#### 4.6.3. Relative frequency of 4-grams and associated phylogenetic relationships of NCF1 sequences

The analysis to the sixteen possible 4-grams solidified the strong polar preference observed at lower orders. The all-polar 4-gram (1111) remained the most frequent local context in 40 of the 41 NCF1 sequences, peaking at 17.36% in A0ABM4FLJ4_9AVES (Figure 17). Conversely, alternating 4-grams and the all-hydrophobic 4-gram (0000) were heavily suppressed across the family, typically falling below 6.8%. This distinct hierarchy highlights an evolutionary preference for extended hydrophilic stretches and a strict avoidance of long non-polar runs or highly alternating patterns.

**Figure 17.**
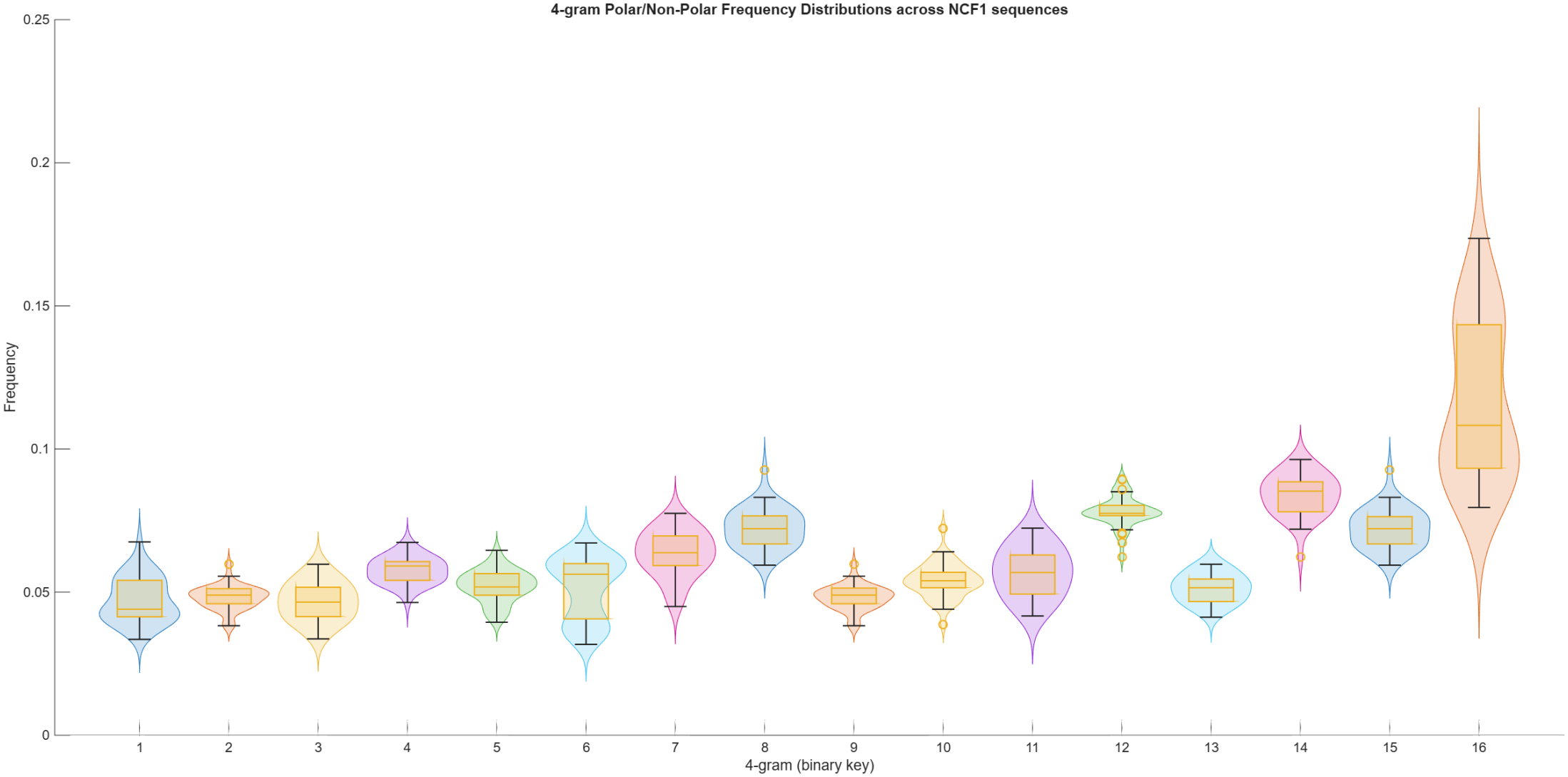
Violin plots of the sixteen binary 4-gram frequencies across the 41 NCF1 sequences. The all-polar 1111-gram was the most frequent 4-mer in 40 of 41 sequences, while the all-hydrophobic 0000-gram and highly mixed 4-grams were consistently suppressed.

A few NCF1 sequences deviated from this consensus. A0A6P6ED59_OCTDE stood out with the flattest, least polar profile in the dataset; it harboured the highest 0000-gram frequency (6.75%) and its top 4-gram barely reached 9.35%, resulting in an unusually compressed distribution. In contrast, A0A6J2R1N8_COTGO almost completely avoided hydrophobic runs, recording the lowest 0000-gram frequency (3.35%). Interestingly, A0A2U3YJL5_LEPWE was the sole sequence where 1111 was not the top 4-gram, being narrowly outranked by 1101 (8.75% vs 7.96%).

At the sequence level, only NCF1_MOUSE and Q3U9Y3_MOUSE retained perfectly identical 4-gram profiles. Pairs that had mirrored each other at the 2-gram level (such as A0A6P5PKW1_MUSCR with NCF1_MOUSE, and A0ABM0KEC1_MICOH with A0ABM2WGI3_MESAU) diverged once the 4-gram context was resolved. This confirms that while a strict polar bias is universally maintained, the fine-scale hydropathic architecture is highly conserved only among the most closely related sequences.

Hierarchical clustering of the normalized 4-gram frequencies (Euclidean distance, threshold = 0.019) resolved the 41 NCF1 sequences into two primary lineages, comprising 12 distinct clusters and 10 singletons (Figure 18; Table 12). The left and central branches (Clusters 1–6) largely grouped the mammalian orthologs, while NCF1_HUMAN branched independently nearby. The rightward branches (Clusters 7–12) consisted of the highly polar sequences, which tightly paired into closely related subsets (e.g., A0ABM4FLJ4_9AVES with A0A6J3E5X2_AYTFU in Cluster-9). Notably, two NCF1 sequences remained unclustered outliers: A0A6P6ED59_OCTDE formed a deep, early-diverging lineage matching its flattened, less polar profile, while A0A6J2R1N8_COTGO branched independently on the far right.

**Table 12:** Sequence clusters derived from hydropathy 4-gram frequencies. The table lists the twelve subclusters of NCF1 sequences generated via hierarchical clustering, grouping proteins that share highly similar polar/non-polar local sequence patterns.

| **Cluster** | **Sets of Proximal NCF1 sequences** |
|---|---|
| Cluster-1 | {NCF1_MOUSE, Q3U9Y3_MOUSE, A0A6P5PKW1_MUSCR, A0ABM0KEC1_MICOH} |
| Cluster-2 | {A0A3Q7U5W3_VULVU, A0A9B0T684_CHRAS, A0ABM4K2Z0_EQUPR} |
| Cluster-3 | {NCF1_BOVIN, A0A340YAF9_LIPVE, A0A384AH76_BALAC} |
| Cluster-4 | {A0A6I9I125_VICPA, A0A8B8RDT0_CAMFR, A0A8B6ZSR4_ORYAF, A0A6J2LZG9_9CHIR} |
| Cluster-5 | {A0A3Q7NHV4_CALUR, A0A6J2E9M7_ZALCA} |
| Cluster-6 | {A0ABM0QKX1_GALVR, A0A2U3UYA7_TURTR, A0A8B7UKD9_CASCN} |
| Cluster-7 | {A0A1V4KPT8_PATFA, A0ACB8ED41_9SAUR} |
| Cluster-8 | {A0A218UEC7_9PASE, A0A6J2J0G2_9PASS} |
| Cluster-9 | {A0A6J3E5X2_AYTFU, A0ABM4FLJ4_9AVES} |
| Cluster-10 | {A0A6P9BZE4_PANGU, A0A6J1UCS7_9SAUR} |
| Cluster-11 | {A0A6P7WVD3_9AMPH, A0A1U7S7U0_ALLSI} |
| Cluster-12 | {A0A9F2MW21_PYTBI, A0A6J0UDP2_9SAUR} |

**Figure 18.**
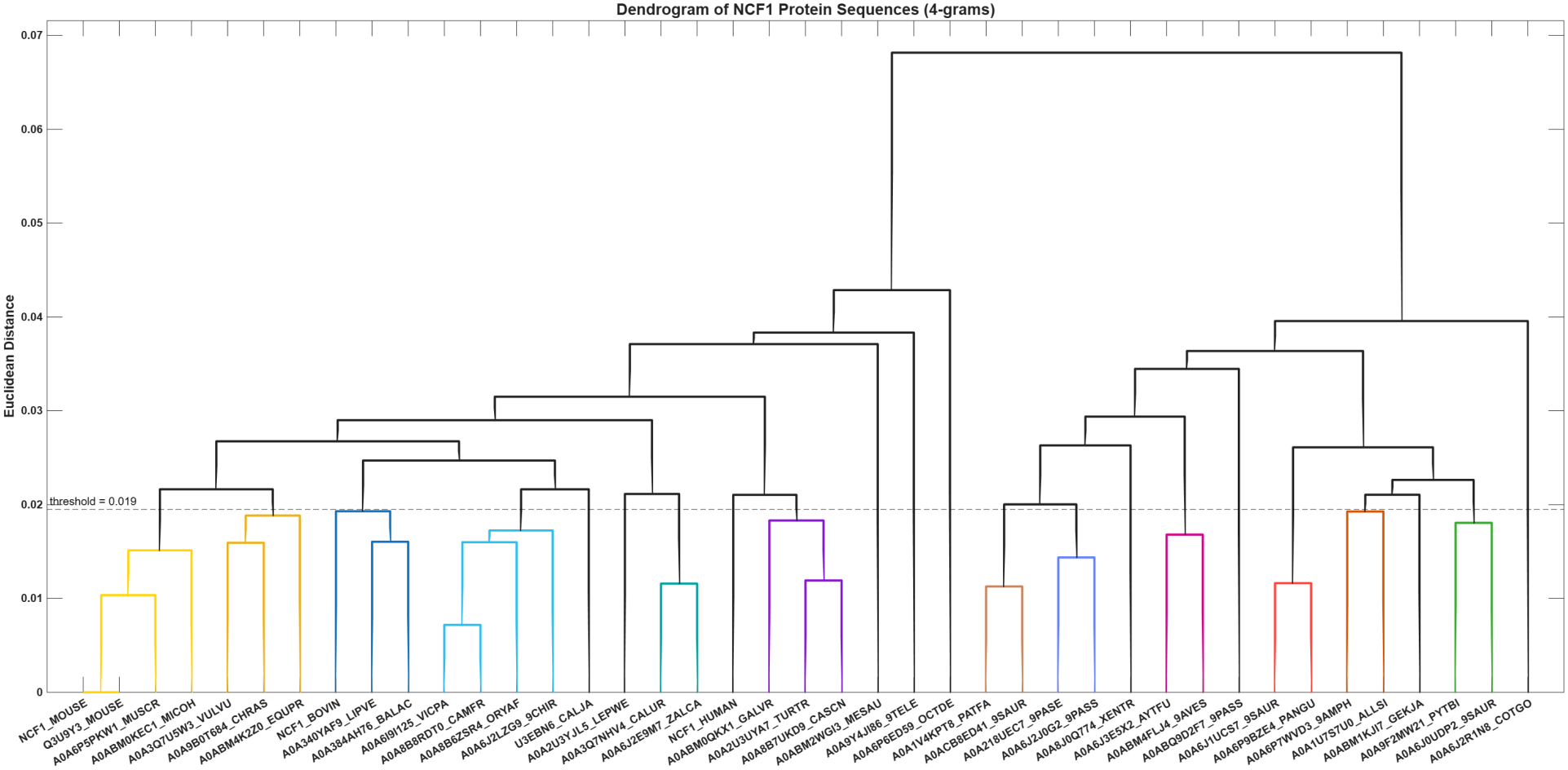
Hierarchical clustering dendrogram constructed using Euclidean distance based on the 4-gram frequency profiles. The horizontal dashed line indicates the distance threshold (0.019) used to delineate the distinct sequence subclusters, which are highlighted by different colored branches.

### 4.7. Proximal relationships of NCF sequences based on the three-dimensional CGR and fractal dimension

Fractal, derived by iterations of the three-dimensional Chaos Game Representation (CGR) (fractal generator) attractors for one NCF1 sequence A0ABM4K2Z0_EQUPR illustrated how the trajectory point clouds evolved as the contraction factor *u* increased. As *u* increased, the attractors transitioned from compact, highly localized clusters into denser clouds that filled the 3D space more thoroughly (Figure 19).

**Figure 19.**
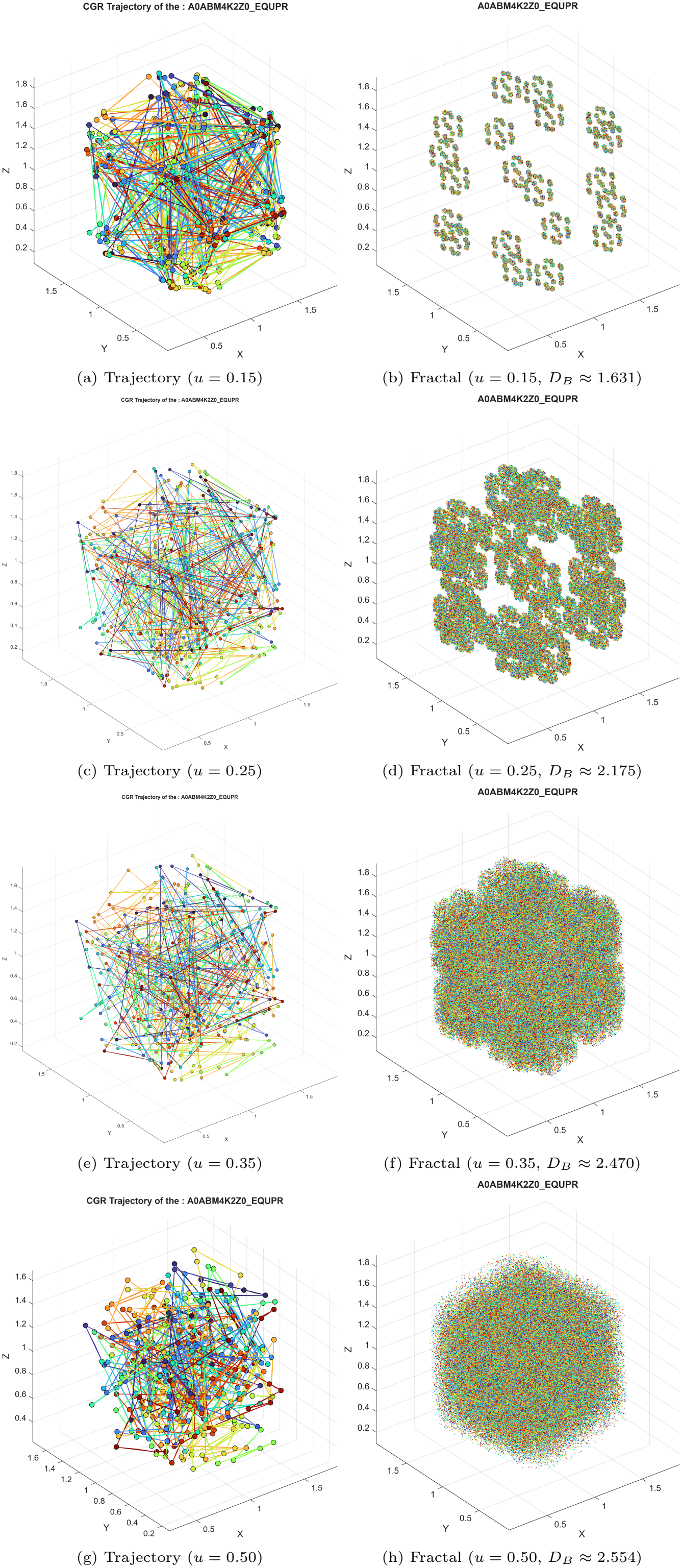
Three-dimensional CGR trajectories and corresponding fractal geometries for A0ABM4K2Z0_EQUPR.

This visual progression was mirrored quantitatively by a predictable, monotonic increase in fractal dimension (*D*_*B*_) across the family. At *u* = 0.15, the attractors were tightly constrained, yielding a median *D*_*B*_ of 1.6310. By the time *u* reached 0.50, the median *D*_*B*_ climbed to 2.5510 (Figure 20). Despite this nearly 1-unit shift in overall scale, the NCF1 family exhibited remarkably rigid conservation. At *u* = 0.50, where the distribution was widest, the entire dataset spanned a mere 0.009 units (ranging from 2.545 in A0A6I9I125_VICPA to 2.554 in A0ABM4K2Z0_EQUPR). At the most sensitive scaling (*u* = 0.15), the full range compressed to just 0.002 units, with A0A6P6ED59_OCTDE peaking marginally at 1.632, while the vast majority clustered tightly around 1.630 to 1.631.

**Figure 20.**
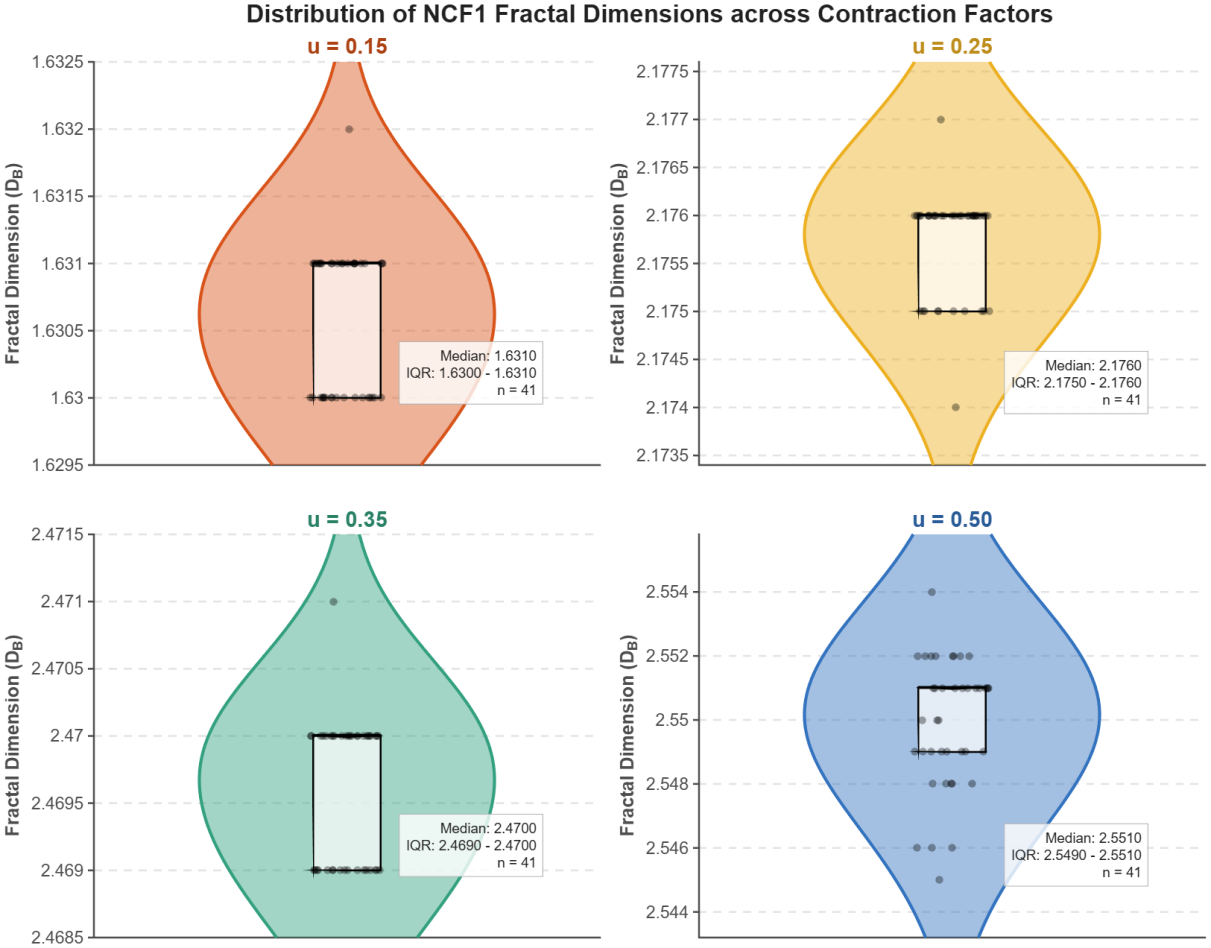
Distribution of fractal dimensions (FD) for the 41 NCF1 sequences across four contraction factors (*u* = 0.15, 0.25, 0.35, and 0.50). Each panel displays a violin plot representing the probability density, overlaid with a central boxplot (median and interquartile range) and individual jittered data points for all 41 sequences. The remarkably narrow Y-axis ranges across all panels highlight the highly conserved spatial scaling of the NCF1 family.

This striking uniformity indicated that the long-range spatial correlations and structural heterogeneity encoded within the NCF1 sequences were highly conserved. Unlike the localized variations captured by the n-grams profiles, the global three-dimensional geometric scaling of the amino acid distributions remained virtually identical across the family, reinforcing the consensus of a monolithic, well-folded protein architecture.

Hierarchical clustering of the four-dimensional fractal dimension (FD) profiles (derived from the 3D CGR) resolved the 41 NCF1 sequences into six distinct clusters and one unclustered outlier at a distance threshold of 0.003 (Figure 21; Table 13). The topology split into two primary lineages. The left branch (Clusters 1–4) grouped 27 sequences characterized by slightly lower FD values at the highest contraction factor, with the human and bovine orthologs tightly pairing within Cluster-2. The right branch (Clusters 5–6) grouped 13 sequences exhibiting marginally higher FD values at *u* = 0.50, notably including the highly polar sequence A0ABM4FLJ4_9AVES within Cluster-5.

**Table 13:** Proximal clusters of NCF1 sequences derived from their four-dimensional fractal dimension (FD) profiles. The clusters were defined by hierarchical clustering of the FD values generated from the 3D CGR across four contraction factors using a Euclidean distance threshold of 0.003.

| **Cluster** | **Sets of Proximal NCF1 sequences** |
|---|---|
| Cluster-1 | {U3EBN6_CALJA, A0A2U3UYA7_TURTR, NCF1_MOUSE, A0ABM0KEC1_MICOH, A0ACB8ED41_9SAUR, A0A384AH76_BALAC, A0ABM2WGI3_MESAU, A0A8J0Q774_XENTR} |
| Cluster-2 | {NCF1_HUMAN, NCF1_BOVIN, A0A8B6ZSR4_ORYAF, A0A1U7S7U0_ALLSI, A0A2U3YJL5_LEPWE, A0A6J2E9M7_ZALCA, A0A3Q7U5W3_VULVU, A0A6J3E5X2_AYTFU} |
| Cluster-3 | {A0A6I9I125_VICPA, A0A6J0UDP2_9SAUR, A0A9B0T684_CHRAS, A0A6P9BZE4_PANGU} |
| Cluster-4 | {A0A1V4KPT8_PATFA, A0A6J1UCS7_9SAUR, A0A8B7UKD9_CASCN, A0A9Y4JI86_9TELE, Q3U9Y3_MOUSE, A0A340YAF9_LIPVE, A0A6P7WVD3_9AMPH} |
| Cluster-5 | {A0A218UEC7_9PASE, A0A6P6ED59_OCTDE, A0ABM4FLJ4_9AVES, A0A3Q7NHV4_CALUR, A0A6J2J0G2_9PASS, A0A8B8RDT0_CAMFR} |
| Cluster-6 | {A0A6J2LZG9_9CHIR, A0ABQ9D2F7_9PASS, A0A9F2MW21_PYTBI, A0ABM0QKX1_GALVR, A0A6J2R1N8_COTGO, A0A6P5PKW1_MUSCR, A0ABM1KJI7_GEKJA} |

**Figure 21.**
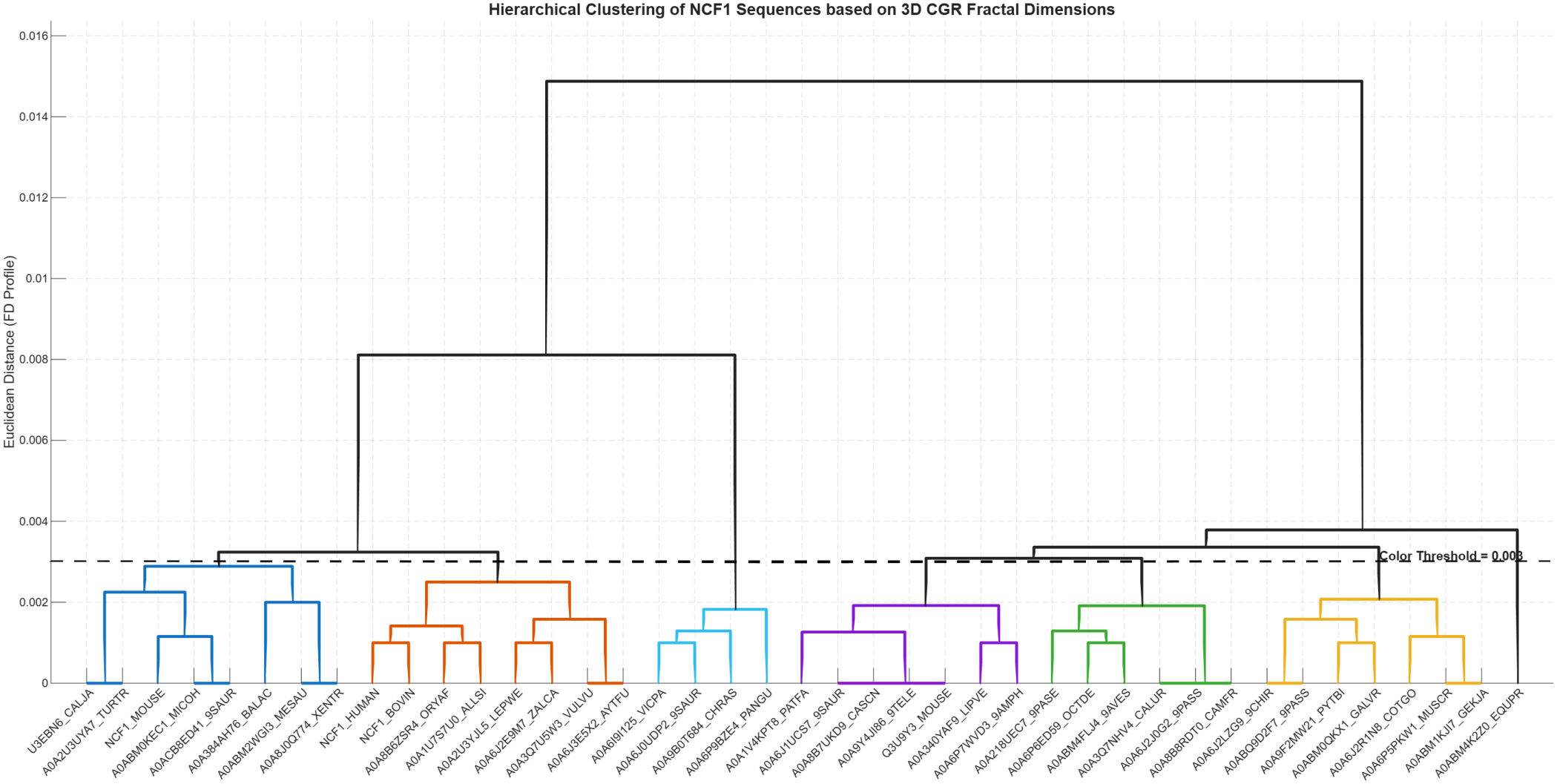
Hierarchical clustering dendrogram of the 41 NCF1 sequences based on the Euclidean distance of their four dimensional fractal dimension profiles. The dashed line marks the color threshold (0.003) used to define distinct sub-clusters.

Notably, A0ABM4K2Z0_EQUPR branched independently as a clear singleton on the far right. This topological placement perfectly mirrored its quantitative data, as it possessed the highest observed fractal dimension in the entire family at *u* = 0.50 (2.554). Despite the exceptionally short Euclidean distances across the tree—reflecting the highly conserved spatial scaling of the family—the clustering successfully segregated this structural outlier from the main NCF1 consensus.

### 4.8. Proximity of NCF1 sequences derived from hydropathy-based dance

Hydropathy-based dance for all 41 NCF1 sequences were obtained. One such dance was presented for the sequence A0ABM4K2Z0_EQUPR in Figure 22. The dance trace of A0ABM4K2Z0_EQUPR exhibited smooth, persistent arcs punctuated by localized high-curvature loops, reflecting a structured arrangement of polar and nonpolar domains. This distinct geometry yields a box-counting dimension of 1.397 and a highly persistent dynamical exponent (*v* = 0.577), confirming a directed, non-random spatial trajectory.

**Figure 22.**
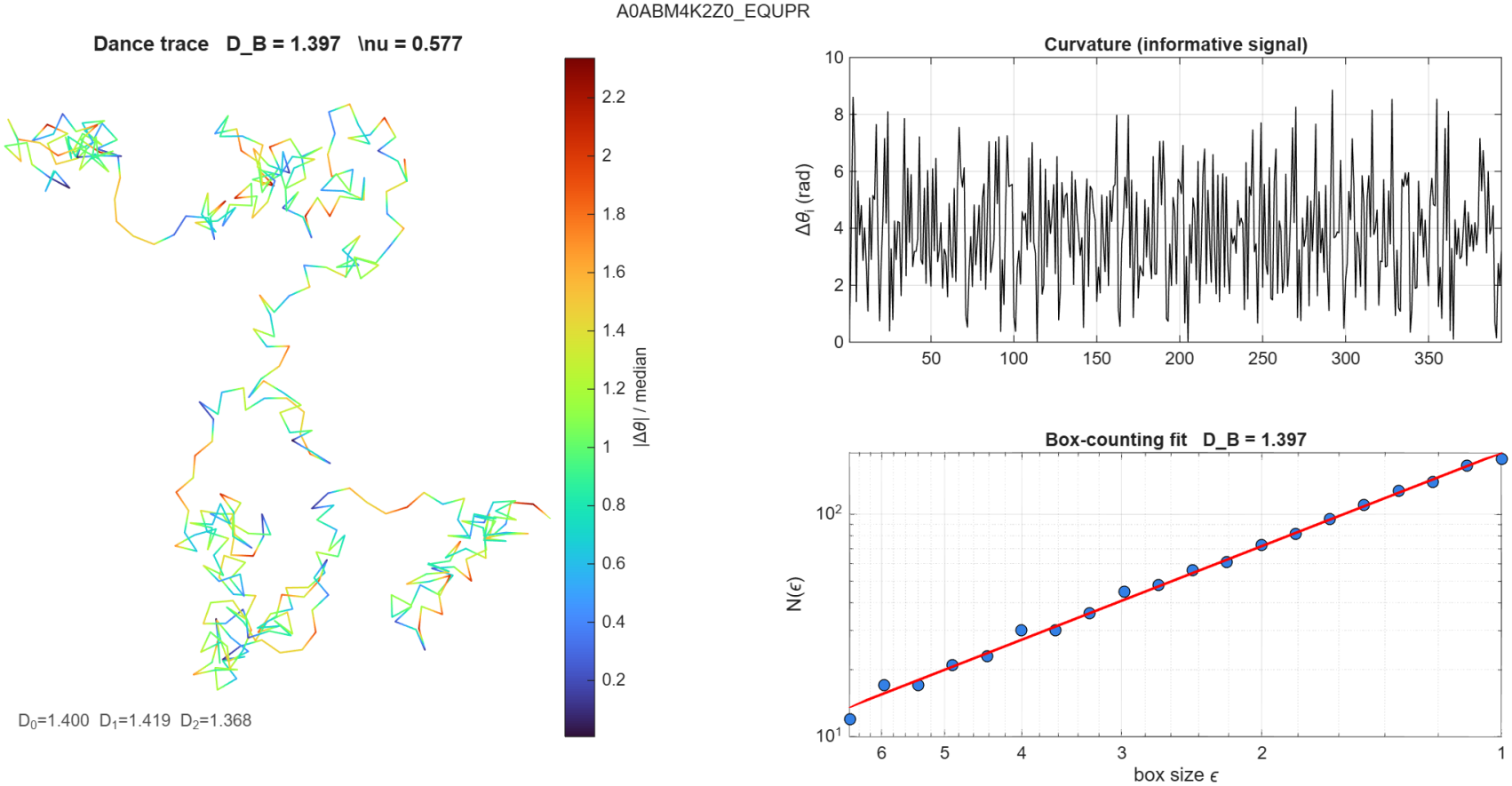
Hydropathy dance analysis of the NCF1 sequence A0ABM4K2Z0_EQUPR. (Left) The 2D cumulative trace generated by the dance encoding, colored by normalized local curvature (| Δ*θ*_*i*_ | /median). The path exhibits smooth, persistent arcs punctuated by localized, high-curvature loops, reflecting structured compositional domains. The resulting geometric and dynamical exponents are annotated below the trace. (Top right) The curvature signal (Δ*θ*_*i*_) along the sequence length, displaying the high-frequency variation in heading driven by local composition. (Bottom right) The box-counting scaling fit (log-log regression of occupied boxes *N* (*ε*) versus box size *ε*), yielding a box-counting dimension of *D*_B_ = 1.397.

**Figure 23.**
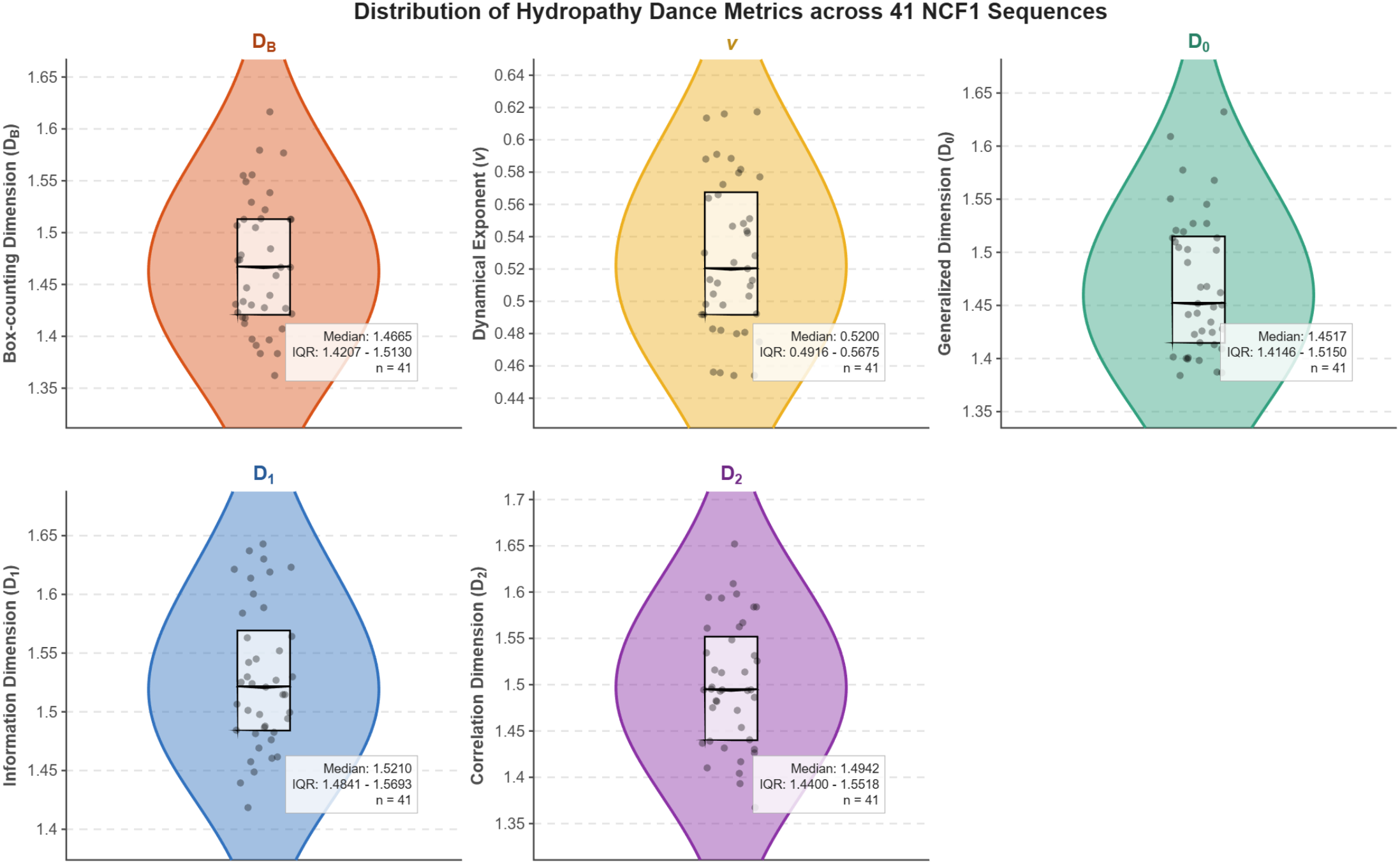
Distribution of hydropathy dance metrics across the 41 NCF1 sequences. Each violin plot displays the probability density for a specific geometric or dynamical exponent derived from the dance encoding: (*D*_B_) box-counting dimension, (*v*) dynamical exponent, (*D*_0_) generalized dimension, (*D*_1_) information dimension, and (*D*_2_) correlation dimension. Individual sequences are represented as jittered black dots, overlaid with a central boxplot indicating the median and interquartile range (IQR). Independent y-axis scaling is used across panels to highlight the specific variance within each metric. Summary statistics (median, IQR, and *n* = 41) are provided in the inset text boxes.

The hydropathy dance translated the spatial arrangement of polar and nonpolar residues into distinct geometric fingerprints across the 41 NCF1 sequences. As visualized in the distribution plots, the box-counting dimension (*D*_B_) remained consistently between 1.36 and 1.62 (median = 1.466), confirming that the NCF1 sequences generally form blocky, finite polar/nonpolar domains rather than perfectly smooth arcs or purely random area-filling walks (Figure 24. The dynamical exponent (*v*) provided further resolution: with a median of 0.520, the majority of sequences exhibited *v >* 0.50, indicating persistent, directed trajectories. This reflects structured, contiguous compositional domains within the proteins rather than purely random hydropathic mixing. The few sequences with *v <* 0.50, such as A0A6J2E9M7_ZALCA (0.454), reflected more bounded, anti-persistent oscillatory behavior.

**Figure 24.**
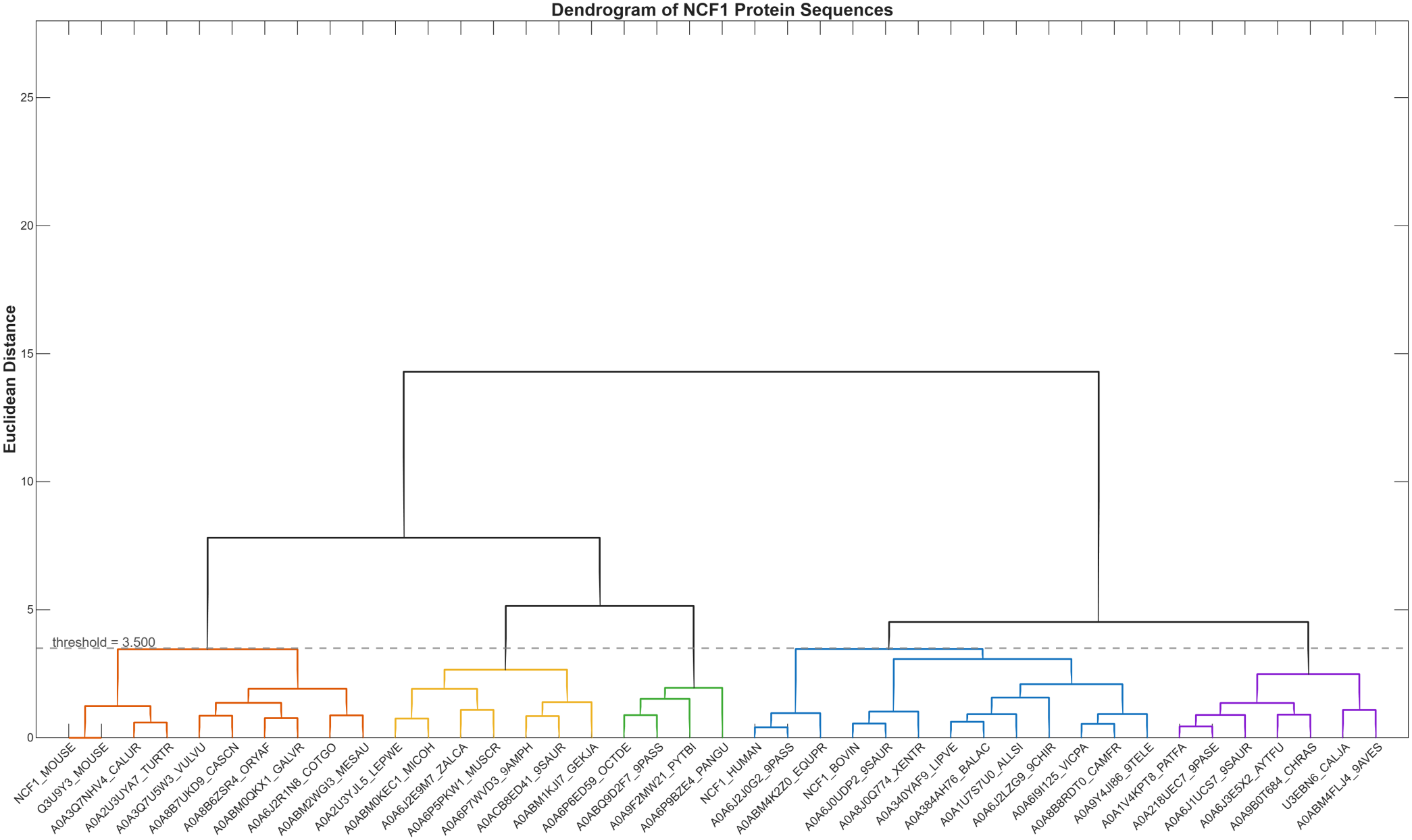
Hierarchical clustering dendrogram of the 41 NCF1 protein sequences based on their hydropathy dance metrics. Clustering was performed using the Euclidean distance between the standardized feature vectors ***ϕ*** = (*D*_B_, *v, D*_0_, *D*_1_, *D*_2_). The dashed horizontal line marks the distance threshold of 3.500, below which branches are colored to denote distinct proximal clusters. The topology groups sequences with similar spatial composition and local hydropathic arrangements.

The generalized dimensions (*D*_0_, *D*_1_, *D*_2_) closely mirrored the *D*_B_ values, forming a relatively narrow multifractal spectrum. This weak multifractality suggests a broadly uniform spatial distribution of compositional domains across the family. Specific sequences stood out at the extremes of geometric complexity. A0ACB8ED41_9SAUR consistently exhibited the highest values across *D*_B_ (1.616), *D*_0_ (1.632), and *D*_1_ (1.643), representing the most spatially complex and dense trace in the dataset. On the other side, A0ABM4K2Z0_EQUPR produced the most geometrically localized trace, recording the lowest *D*_1_ (1.419) and *D*_2_ (1.368). Notably, the correlation dimension (*D*_2_) peaked in A0A9F2MW21_PYTBI (1.652), highlighting a unique clustering of visit densities. Finally, as a strict internal control, the two murine sequences (NCF1_MOUSE and Q3U9Y3_MOUSE) produced identical fingerprints across all five metrics, confirming the precision and reproducibility of the dance encoding.

The proximal relationships among the NCF1 sequences was captured by hierarchical clustering which was performed using the standardized five-dimensional feature vectors ***ϕ*** = (*D*_B_, *v, D*_0_, *D*_1_, *D*_2_) derived from the hydropathy dance. At a Euclidean distance threshold of 3.500, the 41 sequences were partitioned into five distinct clusters, effectively separating sequences with highly persistent, smooth trajectories from those with more complex, area-filling dynamics (Figure 24; Table 14).

**Table 14:** Hierarchical clustering of the 41 NCF1 sequences based on their hydropathy dance metrics.

| **Cluster** | **Sets of Proximal NCF1 sequences** |
|---|---|
| Cluster-1 | {NCF1_MOUSE, Q3U9Y3_MOUSE, A0A3Q7NHV4_CALUR, A0A2U3UYA7_TURTR, A0A3Q7U5W3_VULVU, A0A8B7UKD9_CASCN, A0A8B6ZSR4_ORYAF, A0ABM0QKX1_GALVR, A0A6J2R1N8_COTGO, A0ABM2WGI3_MESAU} |
| Cluster-2 | {A0A2U3YJL5_LEPWE, A0ABM0KEC1_MICOH, A0A6J2E9M7_ZALCA, A0A6P5PKW1_MUSCR, A0A6P7WVD3_9AMPH, A0ACB8ED41_9SAUR, A0ABM1KJI7_GEKJA} |
| Cluster-3 | {A0A6P6ED59_OCTDE, A0ABQ9D2F7_9PASS, A0A9F2MW21_PYTBI, A0A6P9BZE4_PANGU} |
| Cluster-4 | {NCF1_HUMAN, A0A6J2J0G2_9PASS, A0ABM4K2Z0_EQUPR, NCF1_BOVIN, A0A6J0UDP2_9SAUR, A0A8J0Q774_XENTR, A0A340YAF9_LIPVE, A0A384AH76_BALAC, A0A1U7S7U0_ALLSI, A0A6J2LZG9_9CHIR, A0A6I9I125_VICPA, A0A8B8RDT0_CAMFR, A0A9Y4JI86_9TELE} |
| Cluster-5 | {A0A1V4KPT8_PATFA, A0A218UEC7_9PASE, A0A6J3E5X2_AYTFU, A0A6J1UCS7_9SAUR, A0A9B0T684_CHRAS, U3EBN6_CALJA, A0ABM4FLJ4_9AVES} |

The largest group, Cluster-4, contained 13 sequences and notably included NCF1_HUMAN and NCF1_BOVIN along-side a diverse range of vertebrates, from amphibians (A0A8J0Q774_XENTR) to teleost fish (A0A9Y4JI86_9TELE). This grouping highlights a highly conserved baseline geometric fingerprint across widely divergent species. Cluster-1 grouped 10 sequences, predominantly from rodents and ungulates (e.g., NCF1_MOUSE, A0A3Q7U5W3_VULVU), reflecting a shared compositional architecture within these mammalian lineages. The remaining clusters captured sequences with divergent spatial behaviors. Cluster-2 (7 sequences) and Cluster-5 (7 sequences) grouped the structural outliers, including sequences with the highest box-counting dimensions (e.g., A0ACB8ED41_9SAUR at *D*_B_ = 1.616 in Cluster-2) and the highly polar avian sequence A0ABM4FLJ4_9AVES in Cluster-5. Cluster-3 formed a tight group of four sequences (A0A6P6ED59_OCTDE, A0ABQ9D2F7_9PASS, A0A9F2MW21_PYTBI, A0A6P9BZE4_PANGU) characterized by elevated multifractal dimensions.

### 4.9. Evaluation of functional intrinsic disorder in human NCF1

Results of the functional intrinsic disorder analysis of the human NCF1 protein are presented in Figure 25. NCF1 (also known as p47phox), a 390-amino-acid cytosolic organizer of the phagocyte NADPH oxidase complex, contains three folded globular modules: an N-terminal Phox homology (PX) domain (residues 4–121) that binds membrane phosphoinositides (such as PtdIns(3, 4)P_2_), and two tandem Src homology 3 domains, SH3A (residues 159–214) and SH3B (residues 229–284), which cooperate to form a super-SH3 groove that recognizes the proline-rich region of transmembrane CYBA (p22phox). These ordered domains are interspersed with functionally critical intrinsically disordered regions (IDRs), including the PX–SH3A linker (residues 122–158), a short inter-SH3 linker (residues 215–228), and an extended C-terminal tail (residues 285–390). The disordered C-tail encompasses a polybasic autoinhibitory region (AIR, residues 292–340) and a C-terminal serine-, arginine-, and proline-rich module (residues 341–390) containing the proline-rich region (PRR, residues 363–368) that constitutively tethers NCF2 (p67phox) via its C-terminal SH3 domain even in the resting state [40, 16].

**Figure 25.**
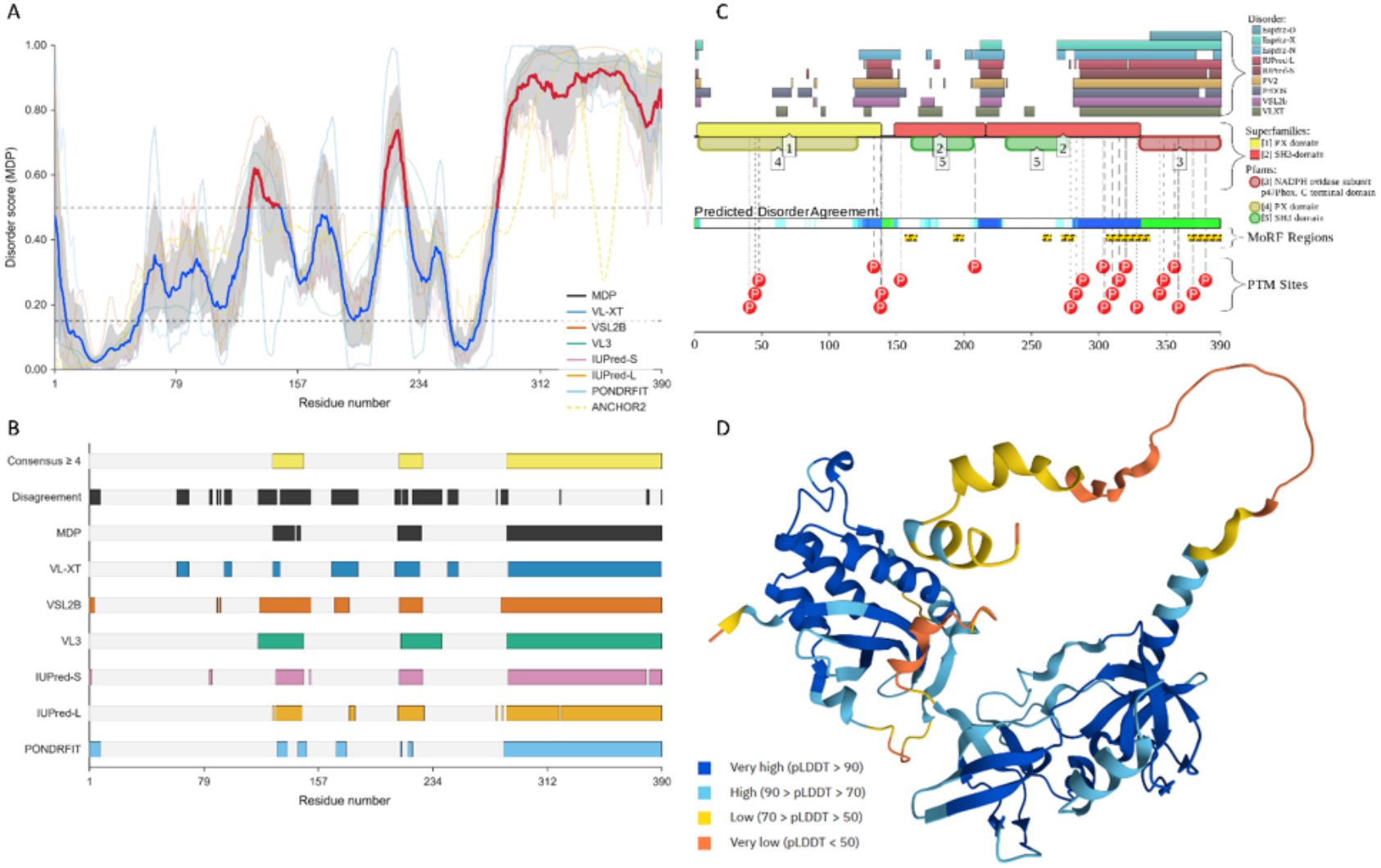
Evaluation of functional intrinsic disorder in human NCF1. **A**. Per-residue intrinsic disorder profile generated by (RIDAO) platform that assembles outputs of six commonly used PONDR® VLS2, PONDR® VL3, PONDR® VLXT, PONDR® FIT, IUPred-Long, and IUPred-Short [36, 37, 43, 44, 45, 46]. Mean disorder profile (MDP) is calculated by averaging the outputs of individual predictors as shown by dashed dark cyan curve, whereas light cyan shade represents the distribution of standard deviations. The plot also incorporates an ANCHOR2 profile generated via the IUPred2A web server [46]. Based on the pairwise energy estimation approach of the original ANCHOR framework, this curve maps the position-specific propensity of intrinsically disordered residues to gain stabilizing energy by interacting with a globular protein partner (with scores above 0.5 typically indicating putative binding sites) [38, 47]. Horizontal lines represent flexibility (solid line at disorder score 0.15) and intrinsic disorder (dashed line at disorder score *≥* 0.5) thresholds. Here, residues with disorder scores 0.5 are considered intrinsically disordered, and residues whose disorder score ranges from 0.15 to 0.5 are assumed to be flexible. **B**. Positions of the intrinsically disordered regions predicted by different disorder predictors assembled by RIDAO. **C**. Functional disorder profile generated by D2P2 platform [48]. **D**. 3D structure of a protein modeled by AlphaFold2. This 3-D structure is color-coded based on the “Predicted Local Distance Difference Test” (pLDDT) values, where segments predicted with very high (pLDDT *>* 90), high (90 *>* pLDDT *>* 70), low (70 *>* pLDDT *>* 50), and very low (pLDDT *<* 50) confidence are shown by blue, cyan, yellow, and orange colors respectively [42]. The folded PX domain and tandem SH3 domains occupy the high-confidence (blue/cyan) core, whereas the PX–SH3A linker, the inter-SH3 linker, and the C-terminal tail (including the AIR, residues 292–340, and the PRR, residues 363–368) occupy the low-confidence (yellow/orange) regions. In this model, the C-terminal tail extends away from the folded core, consistent with the surface-exposed, regulatory role of the AIR and PRR described in the text.

Consistent with this functional organization, Figures 25A and 25B demonstrate that the PX–SH3A linker, the SH3A– SH3B linker, and the regulatory C-terminal tail exhibit high predicted intrinsic disorder propensity. Overall, NCF1 is characterized by a PONDR® VSL2-predicted disorder content (PDC) of 45.38% and a mean disorder score (MDS) of 0.53*±*0.31. Therefore, despite containing ordered domains, NCF1 is classified as highly disordered based on the criteria accepted in the field, where two arbitrary cutoffs for the levels of intrinsic disorder are used to classify proteins as highly ordered (PCD<10%), moderately disordered (10% *≤* PDC<30%) and highly disordered (PDC *≥* 30%)) [41].

Furthermore, Figure 25C presents the functional disorder profile generated by the D2P2 platform, showing that NCF1 is predicted to contain six disorder-based binding sites known as molecular recognition features (MoRFs; residues 156– 164, 192–199, 258–264, 272–281, 305–337, and 366–390)—disordered sub-regions that lack stable structure in the unbound state but undergo disorder-to-order transitions upon binding specific partners—alongside multiple phosphorylation sites.

Finally, Figure 25D displays the predicted 3D structural model of human NCF1 (p47phox) generated by AlphaFold [42]. While the structured, globular modules, specifically the N-terminal PX domain and the central tandem SH3 domains, are predicted with high confidence (pLDDT > 70–90), the model reveals distinct low-confidence regions (pLDDT < 50) primarily located within the inter-domain linkers and the C-terminal tail. In AlphaFold predictions, such low pLDDT scores strongly correlate with intrinsic structural disorder and conformational flexibility.

In the AlphaFold model (Figure 25D), the folded PX domain and the tandem SH3 domains occupy the high-confidence core, whereas the PX–SH3A linker, the inter-SH3 linker, and the C-terminal tail occupy the low-confidence regions. This spatial arrangement places the AIR (residues 292–340) and the PRR (residues 363–368) on the exposed tail rather than within the folded core. Because the super-SH3 groove is formed at the SH3A–SH3B interface and the PX phosphoinositide-binding face lies on the PX domain, the model is consistent with the tail being positioned to mask these surfaces in the resting state and to release them upon phosphorylation. The model does not, however, resolve the individual contacts: the AIR and PRR are low-confidence segments, and the model therefore supports the spatial plausibility of the autoinhibitory and NCF2-tethering roles rather than demonstrating specific residue-level interfaces.

Functionally, this structural plasticity governs NCF1 activation. In the resting state, the disordered C-terminal AIR folds back to intramolecularly mask the tandem SH3 groove and PX domain, locking NCF1 in an autoinhibited conformation. Upon phagocyte stimulation, multi-site phosphorylation of clustered serine residues within the disordered C-terminal tail (spanning Ser303–Ser379, including Ser303, Ser304, Ser315, Ser320, Ser328, Ser345, Ser348, Ser359, Ser370, and Ser379) disrupts this intramolecular clamp. This triggers a conformational switch that exposes the PX and tandem SH3 domains for membrane phosphoinositide and CYBA (p22phox) binding, thereby driving membrane translocation and NOX2 complex assembly [40]. Meanwhile, the C-terminal PRR (residues 363–368) keeps NCF1 constitutively tethered to NCF2 (p67phox) even in the resting state [16].

### 4.10. Multidimensional quantitative profiling reveals individual sequence signatures of NCF1

We evaluated the structural and evolutionary proximity of NCF1 protein sequences by integrating hierarchical clustering data from nine distinct quantitative features: amino acid frequency, sequence homology, dipeptide composition, normalized polystring lengths, hydropathy n-grams (2-, 3-, and 4-grams), four-dimensional fractal dimension profiles, and hydropathy dance metrics. Synthesizing these nine clustering partitions demonstrated that the NCF1 protein lacked a single, unified cumulative proximity matrix. Instead, the cluster membership of sequences shifted radically depending on the quantitative feature applied. This lack of convergence was a critical finding: it proved that NCF1 sequences were not defined by a single linear hierarchy, but rather by multidimensional individual signatures, wherein phylogenetic history, local biophysical chemistry, and abstract topological geometry contributed independently to their proximity.

#### 4.10.1. Phylogenetic conservation governs primary sequence proximity

When clustering was driven by primary sequence features—specifically amino acid frequency (Table 4), sequence homology (Table 6), and dipeptide composition (Table 8)—NCF1 proximity strictly obeyed evolutionary phylogeny. In these dimensions, distinct taxonomic clades were highly stable. For instance, the core murine rodents (NCF1_MOUSE, Q3U9Y3_MOUSE, A0A6P5PKW1_MUSCR), primates (NCF1_HUMAN, U3EBN6_CALJA), and marine pinnipeds (A0A3Q7NHV4_CALUR, A0A6J2E9M7_ZALCA) formed exclusive, invariant clusters. Furthermore, dipeptide composition (Table 8) acted as a high-resolution lens, splitting larger macro-groups (such as all Cetartiodactyla) into highly specific sub-clusters (e.g., separating whales/dolphins from camels/alpacas), confirming deep evolutionary divergence.

#### 4.10.2. Biophysical convergence drives structural cross-taxa bridges

As the clustering metrics transitioned to structural chemistry—specifically normalized polystring lengths (Table 9) and hydropathy 2-, 3-, and 4-grams (Tables 10, 11, 12)—the strict evolutionary signal dissolved. Organisms from distinct lineages converged based on shared local polar/non-polar patterns, creating unexpected structural bridges. For example, the Red Fox (A0A3Q7U5W3_VULVU) consistently abandoned its carnivore lineage to cluster directly with core rodents and avian species based on shared hydropathy signatures. Additionally, a strong structural alignment emerged between Bovine (NCF1_BOVIN) and Cetaceans (LIPVE, BALAC). These features prove that while primary sequences diverge, the local biophysical chemistry of the NCF1 protein remains highly convergent across diverse taxa.

#### 4.10.3. Topological profiling uncovers geometric conservation

The most profound evidence for individual sequence signatures emerged from highly derived mathematical metrics: four-dimensional fractal dimension (Table 13) and hydropathy dance metrics (Table 14). Rather than adhering to traditional biological boundaries, these abstract topological lenses revealed an alternative layer of sequence organization. Under these metrics, unique cross-kingdom mega-clusters formed, grouping organisms as disparate as a primate, a dolphin, a mouse, a bat, a reptile, a whale, and an amphibian into a single proximal cluster (Cluster-1, Table 13).

Despite this profound reorganization, certain topological invariants emerged. A highly unexpected and resilient bridge was observed between NCF1_HUMAN and NCF1_BOVIN. While evolutionarily and sequentially distant, their NCF1 sequences converged consistently under 3-gram hydropathy (Table 11), fractal dimension (Table 13), and hydropathy dance metrics (Table 14). This pairing revealed a hidden geometric invariance between these two distinct mammals that only high-dimensional topology could expose.

## 5. Discussion and concluding remarks

Inborn errors of immunity (IEIs) represent a clinically heterogeneous group of Mendelian disorders, among which Chronic Granulomatous Disease (CGD) stands out for its marked severity in populations where consanguineous marriage is common and recessive alleles are therefore enriched. The NCF1 (p47phox) protein serves as a critical cytosolic organizer for the phagocyte NADPH oxidase complex, and its dysfunction leads to CGD and autoimmune conditions such as Systemic Lupus Erythematosus (SLE). To better understand how the spatial architecture of this protein dictates its function, and to establish a broad genomic baseline, the present study integrated a set of distinct quantitative features across 41 NCF1 sequences from diverse organisms. By moving from primary sequence metrics to biophysical chemistry and abstract topological geometry, we aimed to decipher the multidimensional structural signatures that govern NCF1’s integrity and evolutionary constraints.

Focusing first on primary sequence composition, the NCF1 sequences shared a highly conserved architectural bias. They exhibited high frequencies of arginine (R), glutamic acid (E), lysine (K), and proline (P), alongside a near-universal depletion of cysteine (C). The high, narrow Shannon entropy (4.098–4.198 bits) indicated strong purifying selection, preventing extreme compositional shifts across species. This specific composition was biologically deliberate, reflecting NCF1’s cytosolic function and its reliance on proline-rich and basic residues to facilitate essential SH3-domain protein interactions. Furthermore, window-wise and site-specific entropy analyses revealed a distinct “block-like” architecture. The presence of completely invariant, zero-entropy blocks (e.g., positions 195–210, 260–285, and 365–380) and highly conserved basic/proline-rich C-terminal motifs (e.g., 385–400) highlighted specific structural domains under intense purifying selection. The unusually low Degree of Dominance (DD; mean 16.7%) confirmed that NCF1 relied on a distributed balance of residues rather than isolated hotspots.

The clinical implications of this primary architecture were substantial. Given that specific pathogenic variants, such as the c.75_76delGT founder mutation, are highly prevalent in certain populations, mapping these clinical variants against our structural model provided critical insight. The vast majority of severe, disease-causing mutations—including nonsense mutations and deletions—fell directly within these rigidly conserved zero-entropy blocks. Their strict conservation confirmed that these sites were structurally indispensable. In contrast, hypomorphic missense variants linked to SLE (such as the R90H mutation) fell outside these absolute zero-entropy blocks. This spatial mapping aligned closely with clinical phenotypes: mutations in the rigid core caused fatal immunodeficiency, whereas partial, localized disruptions led to autoimmune susceptibility.

Transitioning from primary sequence to local biophysical chemistry, the strict evolutionary signal faded, revealing pronounced structural convergence. The longest polystring (repeat-tract) analysis demonstrated a stringent, global restriction against extended repetitive tracts, thereby preventing protein aggregation. Similarly, hydropathy n-gram analyses revealed a strict universal bias toward adjacent polar residues and a suppression of extended non-polar stretches. This polar-hydrophilic dominance was a hallmark of soluble, well-folded cytosolic proteins, ensuring NCF1 avoided aberrant membrane insertion before translocating to the phagosome. Because NCF1’s stability relied on this precise polar bias and avoidance of repeats, pathogenic mutations that artificially introduced hydrophobic stretches or extended tracts (e.g., via frameshifts) would likely have induced severe misfolding, providing a structural mechanism for the immune dysregulation observed in IEI patients.

Expanding our analysis to global topological constraints, highly derived mathematical metrics offered the strongest evidence of NCF1’s rigidity. The three-dimensional Chaos Game Representation (CGR) and fractal dimension analysis revealed extraordinary geometric conservation; across all organisms, the long-range spatial correlations spanned a remarkably narrow fractal dimension range of just 0.009 units. This geometric rigidity was further corroborated by hydropathy dance metrics, where sequences exhibited persistent, directed trajectories and narrow multifractal spectra. Taken together, these topological lenses confirmed that NCF1 formed structured, contiguous compositional domains that could not tolerate deviation. Because the protein required a highly specific spatial packing to orient its interaction domains for NADPH oxidase assembly, genetic variants disrupting these long-range spatial correlations would likely have caused catastrophic misfolding.

The intrinsic disorder analysis positioned NCF1 as a highly disordered protein (PDC 45.38%; MDS 0.53 *±* 0.31), with disorder concentrated in the PX–SH3A linker, the inter-SH3 linker, and the C-terminal tail, while the PX and tandem SH3 domains remained ordered and AlphaFold-confident. This architecture was functionally instructive: the disordered C-terminal tail harbored the AIR and PRR, enabling the tail to fold back and mask the SH3 groove and PX domain in the resting state, and to release this autoinhibition upon multi-site phosphorylation. The six predicted MoRFs and clustered phosphosites within these disordered segments were consistent with a disorder-to-order switching mechanism that governed membrane translocation and NOX2 assembly, while the constitutive PRR–NCF2 tethering illustrated how a single disordered region could simultaneously encode autoinhibition and partner binding. Thus, NCF1 illustrated how intrinsic disorder converted a modular enzyme organizer into a phosphorylation-responsive molecular switch. The AlphaFold model (Figure 25D) is consistent with this arrangement: the folded PX and SH3 domains form the high-confidence core, while the AIR- and PRR-containing C-terminal tail lies in a low-confidence, surface-exposed region, spatially positioned to mask and release the SH3 groove and the PX face during activation. This interpretation is structural and qualitative; residue-level interface mapping will require a dedicated analysis.

A comprehensive synthesis of these quantitative features established a critical biological insight: NCF1 lacked a single, unified cumulative proximity matrix. Instead, its structural and evolutionary relationships shifted dynamically depending on the applied mathematical lens. This lack of convergence indicated that NCF1 was defined by multidimensional individual signatures, wherein phylogenetic history, local biophysical chemistry, and abstract topological geometry operated independently. Despite this profound reorganization across metrics, several resilient invariants emerged. The core murine pair (NCF1_MOUSE, Q3U9Y3_MOUSE) and the cetacean pair (A0A340YAF9_LIPVE, A0A384AH76_BALAC) remained robustly clustered. Most surprisingly, a geometric invariant emerged between NCF1_HUMAN and NCF1_BOVIN; although evolutionarily distant, they converged consistently across 3-gram hydropathy, fractal dimension, and hydropathy dance metrics, revealing a deep structural baseline only exposed by high-dimensional topology.

The primary strength of this study lay in its comprehensive, multi-metric approach, integrating distinct quantitative features to map NCF1 from primary sequence to abstract mathematical topology. The cross-species analysis of 41 organisms allowed for the precise identification of evolutionary constraints and clinical mutation hotspots, translating complex mathematical models into a predictive framework for evaluating clinical variants. This offers value for clinical genomics in populations where consanguinity increases the likelihood of recessive IEIs. However, this study had certain limitations. Our analyses were entirely computational and sequence-based; they did not account for post-translational modifications (PTMs)—such as the phosphorylation of p47phox required for its activation—or the dynamic conformational changes that occurred during active NADPH oxidase assembly. Future studies should focus on experimental validation of these structural predictions using *in vitro* protein folding assays and targeted clinical genomic screening within affected cohorts.

In conclusion, NCF1 proximity and structural integrity were defined by a unique interplay of evolutionary, biophysical, and topological quantitative signatures rather than a single, linear ranking of features. By mapping these multidimensional structural invariants, this study strengthened the quantitative understanding of the NCF1 spatial architecture. Ultimately, these multidimensional baselines provided a predictive framework for clinical genomics. Because the spatial and compositional constraints of NCF1 were so tightly conserved, pathogenic mutations—such as those causing the inborn error of immunity CGD or autoimmune conditions like SLE—likely disrupted specific baseline signatures rather than globally destroying the protein. This offered a robust foundation for evaluating the severity of genetic variants and informing future targeted therapeutic strategies, supporting precision medicine efforts for NCF1-associated diseases.

## Author contributions statement

SSH, KH and VNU formulated and designed the study protocol. SSH and VNU conducted the theoretical experiments and obtained the results. All authors reviewed, checked, and approved the final version of the manuscript.

## Acknowledgements

The authors thank Debaleena Nawn for her help in developing the mathematical methods used in the Mathematical Genomics framework. Her contribution is gratefully acknowledged; any interpretation of the results remains the responsibility of the authors.

## Declaration of competing interest

The authors declare no conflict of interest.

## Notes

### Competing Interest Statement

The authors have declared no competing interest.

